# Inferred Developmental Origins of Brain Tumors from Single Cell RNA-Sequencing Data

**DOI:** 10.1101/2024.05.10.593553

**Authors:** Su Wang, Rachel Naomi Curry, Anders W Erickson, Claudia Kleinman, Michael D. Taylor, Ganesh Rao, Benjamin Deneen, Arif O. Harmanci, Akdes Serin Harmanci

## Abstract

The reactivation of neurodevelopmental programs in cancer highlights parallel biological processes that occur in both normal development and brain tumors. Achieving a deeper understanding of how dysregulated developmental factors play a role in the progression of brain tumors is therefore crucial for identifying potential targets for therapeutic interventions. Single-cell RNA sequencing (scRNA-Seq) provides an opportunity to understand how developmental programs are dysregulated and reinitiated in brain tumors at single-cell resolution. Here, we introduce COORS (Cell Of ORigin like CellS), as a computational tool trained on developmental human brain single-cell datasets that enables annotation of “developmental-like” cell states in brain tumor cells. Applying COORS to various brain cancer datasets, including medulloblastoma (MB), glioma, and diffuse midline glioma (DMG), we identified developmental-like cells that represent putative cells of origin in these tumors. Our work adds to our cumulative understanding of brain tumor heterogeneity and helps pave the way for tailored treatment strategies.

## Introduction

One of the greatest challenges to finding a cure for brain cancers is the robust inter- and intra-tumoral heterogeneity that characterizes these tumors^1–4^. This heterogeneity contributes to disease progression and is a key reason therapeutic approaches fail to prevent disease recurrence. Although the genetic evolution of cancer cells is a critical determinant, tumor heterogeneity is also influenced by non-genetic factors including varying developmental cellular programs, which include stem, progenitor, and senescent cell states^5,6^. Prior studies have demonstrated that aberrant expression of neurodevelopmental programs is pervasive in brain tumors and is largely driven by the reactivation of developmental transcriptional states that are acquired by genomic and epigenomic changes. Given the complexity of cell types and an array of developmental states, isolating a single cell type of origin poses a difficult task, however, a more thorough examination of brain tumor transcriptomics alongside transcriptional signatures of neurodevelopmental cell types may shed light on the origins of brain cancer. To gain a deeper understanding of which developmental cell types brain tumors most closely resemble, we hypothesized that tumor cell lineages can recapitulate cell lineages encountered in the developing brain. While tumors exhibit a multitude of dysregulated pathways, existing evidence, particularly in pediatric tumors, supports this hypothesis^6–10^. We, therefore, focused on employing developmental expression modeling trained on human brain atlases that span various developmental time points. This modeling approach allows us to characterize tumor cells by overlaying their gene expression patterns onto those of early neurodevelopmental stages. By identifying and studying these myriad cellular states from development, our goal is to uncover insights into the origins and behavior of brain tumors, ultimately paving the way for more effective treatment strategies and improved patient outcomes.

Single-cell RNA sequencing (scRNA-Seq) provides an opportunity to dissect the complex cellular states during development and in health and disease^11^. However, it is computationally challenging to decipher the spectrum of heterogeneous developmental cell states in tumor cells using scRNA-Seq. Accurate identification of developmental-like cell states necessitates a comprehensive understanding of the interactions among all genes, which, in turn, requires a substantial amount of gene expression data. In this study, we developed COORS, a computational tool to annotate each developmental cell state in tumor cells at single cell resolution. COORS uses multilayer perceptron model for cell of origin classification and cell age regression using developing brain scRNA-Seq datasets from previously published scRNA-seq datasets, comprising approximately 1M million cells from developing human and mouse brains^6,12–16^. We used COORS to predict developmental-like analogs in pediatric and adult tumors using public and in-house MB^17^, DMG^14^ and glioma scRNA-Seq data, which revealed unique developmental cell types as putative cells of origin in each brain cancer subtype.

## Results

### The overall workflow of COORS algorithm

The basic motivation of COORS algorithm is to utilize existing primitive cell types identified in developing normal brain tissues to annotate the potential origins of cells in tumor samples. In this regard, COORS aims to capitalize on the large amount of healthy developmental transcriptional profiles to identify similar programs in tumor tissues. This is in essence different from existing cell type annotation methods with two aspects: 1) Existing methods for cell type annotations in healthy tissues rely on exact matching of the cell types. 2) Existing methods for cell type annotations in tumor samples focus mainly on detecting tumor cells and do not focus on annotating the origin states for cells.

For training COORS models, we train a multilayer perceptron model for cell of origin classification and cell age regression using developing brain scRNA-Seq datasets^6,12–16^ (**Table S1**). Assuming we have reference data with two origin-like cell types A and B, we train a neural network-based cell of origin classifier using this reference data, saving the model in our repository **(Figure 1A-B)**. Concurrently, we train two neural network-based cell age regressors, one for cell origin A and another for cell origin B, also saving these trained models in the repository. In the assignment step, each tumor needs to be matched to correct set of origin-like cell type assignment models. This is justified since each cancer originates at different regions of brain (neocortex vs cerebellum vs pons) which can be used to select the COORS models. After selecting the relevant origin-like cell models, we map tumor cells to developing healthy brain cells by using the pre-trained models. We predict the cell of origin for the testing dataset using the pre-trained cell of origin classifier. For each developmental-like cell type, we further predict cell age using the corresponding pre-trained cell age regressors. Finally, we conduct SHapley Additive exPlanations (SHAP) analysis^18^ to extract essential features from our machine-learning neural network models, identifying tumor-specific developmental-like gene markers for each cell type and age within our training dataset **(Figure 1A-B).**

**Figure 1.**
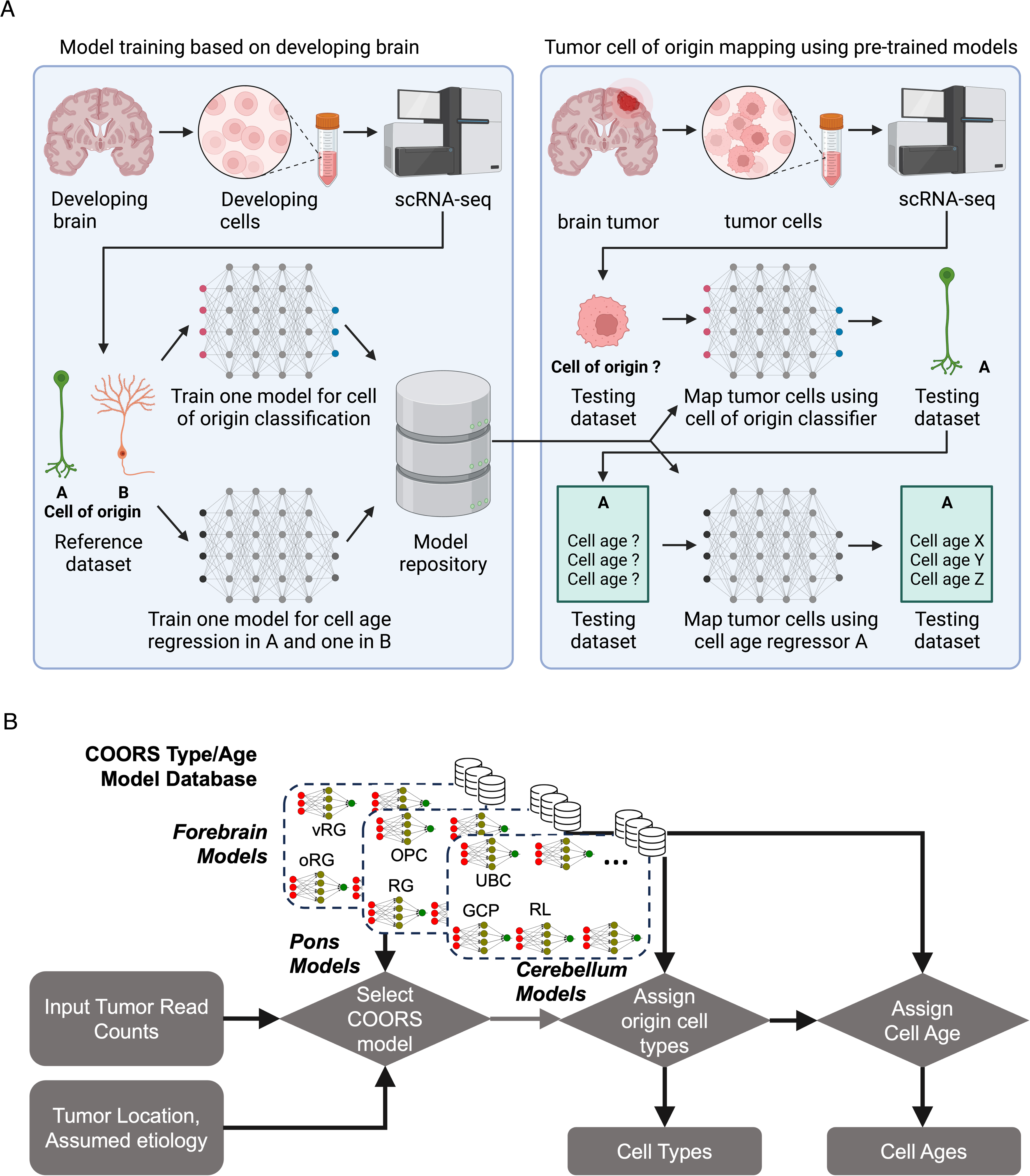
Overview of COORS algorithm. (A) In the first step, neural network models are trained for cell of origin classification and cell age regression using developing brain scRNA-Seq datasets, and the models saved in the repository. In the second step, these pre-trained models are used to map scRNA-Seq tumor cells to developing brain cells, predicting cell origin and age while conducting SHAP analysis to identify tumor-specific gene markers. (B) Tumors are matched with specific origin-like cell type assignment models based on their region of origin within the brain (e.g., neocortex vs cerebellum vs pons), enhancing the precision of the COORS application. Post-model selection, the mapping of tumor cells to developing healthy brain cells is performed through the application of these pre-trained models, as depicted in the schematic.

### Validation of COORS algorithm on scRNA-Seq Medulloblastoma data

Medulloblastoma (MB) is a pediatric brain tumor that is closely associated with early hindbrain development and can be classified into four main molecular subgroups^6,7,10,19–22^. The WNT-activated subgroup is defined by mutations in the WNT signaling pathway and generally displays a favorable prognosis. In contrast, the Sonic Hedgehog (SHH)-activated subgroup results from mutations in the SHH pathway and may have varying clinical outcomes. Group 3 (GP3) MBs have a distinct gene expression profile and are typically associated with a poorer prognosis. Finally, Group 4 (GP4) tumors, marked by a specific gene expression pattern, tend to have intermediate clinical outcomes. Understanding the origin of these four tumor subtypes might lead to the development of improved treatment strategies.

We applied the COORS algorithm to previously published MB scRNA-Seq data ^17^, where subgroup annotations are available for each sample, containing 29 samples and approximately ~40K cells in total (**Figure 2A**). Using COORS algorithm, we have used the pre-trained cell type and cell age models, derived from scRNA-Seq data of developing cerebellum, to map tumor MB cells (**Figure 2B**)^13^. We have not focused on WNT subgroup tumor cells because the WNT subgroup is known to originate from the lower rhombic lip (LRL) adjacent to the brainstem, rather than from the upper rhombic lip (URL) in the cerebellum^23^. COORS maps SHH subgroup tumor cells to granule cell precursor (GCP), GP4 subgroup tumor cells to unipolar brush cell (UBC-CN), Granule Neurons (GN) predominantly to SHH and secondarily to GP4 and GP3 subgroup tumor cells, and Rhombic Lip (RL) subgroup tumor cells to GP4 and GP3 (**Figure 2B-F**). These findings align with recent publications on MB subgroup cell origins^8,9,24,25^. Earlier research identified GN as progenitor cells for SHH-induced MB, while another study proposed that the rhombic lip subventricular zone (RLSVZ) progenitor cells are the source of GP3 and GP4 MB^8,9,24,25^. Additionally, one study suggested that GP4 originates from the UBC lineage^8^. In addition, we conducted SHAP analysis^18^ to extract critical features from our machine-learning neural network models, which returned new and known marker genes associated with upper rhombic lip-derived cell types and the MB subgroups to which they correspond. For example, markers associated with the external granule layer or GCP identity such as *NDST3*^26^, *CBFA2T2*^24^, and *UNC13C*^27^; GC identity and maturation such as *RBFOX3*^28,29^, *GRIK2*^24^, *ROBO1* whose paralogs are essential in GC migration^30^, with *MSI2* likely as it marks GCPs in the external granule layer but not postmitotic granule cells in the internal granule layer^31^; UBC identity such as *LMX1A*^32^, *CACNA2D1*^8,33^, *RELN*^34^, known GP4 oncogene *ERBB4*^24,35^ and *JMJD1C/KDM3C*, a putative H3K9 demethylase whose paralogs are recurrently somatically altered in GP4 MB^20,24,36^; and RL identity such as *OTX2*^13^, *HIST1H4C, -3B, and -1C*^37^, and *SLIT2*^30^ (**Figure 2G-J**).

**Figure 2.**
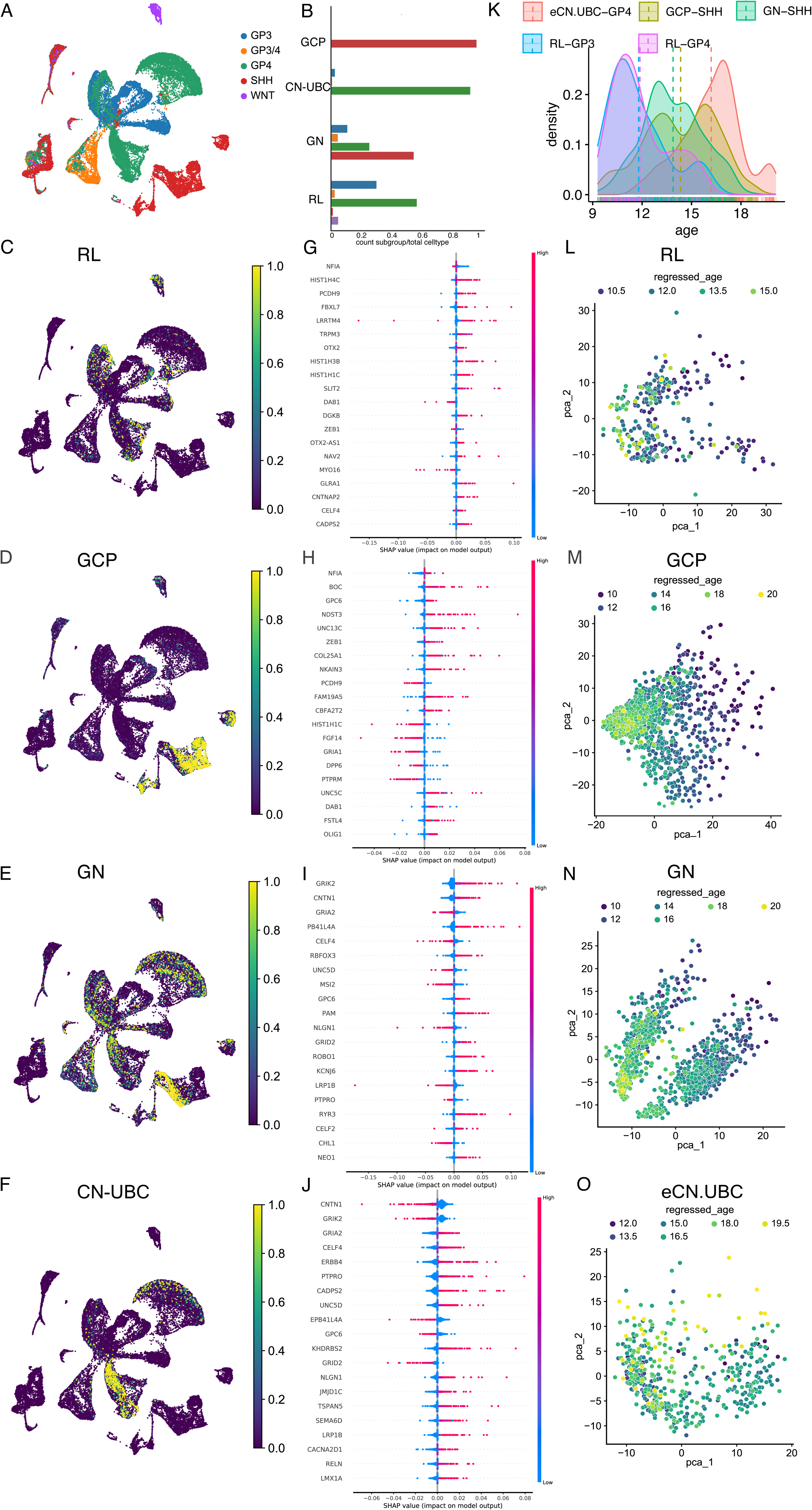
Characterization of developmental-like cell states in MB scRNA-Seq data. (A) Tumor subgroups are shown for MB scRNA-seq dataset. (B) Barplots showing the distribution of MB cells within each tumor subgroup mapped to individual developing cell types. (C-F) Developmental cell type probability scores are shown for RL, GCP, GN and UBC cell types. (G-J) The figure displays the results of SHAP analysis, showing the top impactful genes from each cell type, RL, GCP, GN and UBC respectively, in our training dataset. (K) Distribution of age mapping within each tumor subgroup and their respective mapped cell of origin pairs. (L-O) PCA plots showing the mapping of developmental ages for tumor cells mapped to various developmental origins, GCP, CN-UBC, GN, and RL respectively.

Next, we predicted the cell age of each identified developmental cell type within MB tumor cells using our pretrained cell age regressor models **(Figure 2K-O).** Interestingly, GP4 subgroup cells mapping to UBC mostly correspond to the later weeks in development by 17 post-conceptional weeks (PCW), while GP3 and GP4 subgroup cells mapping to RL mostly correspond to the earlier weeks by 11 PCW in development (**Figure 2K)**. The DEGs between tumor cells and their respective cells of origin, as well as the DEGs between the cell of origin and the following developmental stage cell type are listed in **Table S2 (Figure S1-8)**.

### Application of COORS algorithm on pediatric diffuse midline glioma (DMG)

Next, we applied the COORS algorithm on previously published H3.1/H3.2 and H3.3 histone 3 K27M-mutant DMG scRNA-Seq data containing 13 samples and approximately 47K cells in total (**Figure 3**)^6,14^. Using COORS algorithm, similar to our previous application in MB data, we have used the pre-trained cell type and cell age models, derived from scRNA-Seq data of developing mouse pons brain^6^, to map pediatric glioma cell origins (**Figure 3A-C**). Consistent with previous studies we have found that H3.3K27M gliomas mapped to oligodendrocyte-like and neuron-like cells, as previously reported^3,6,14,38^. More specifically, H3.1/2K27M tumors mapped to ependymal-like cells whereas H3.3 mapped to neuronal intermediate progenitor cells (IPCs).

**Figure 3.**
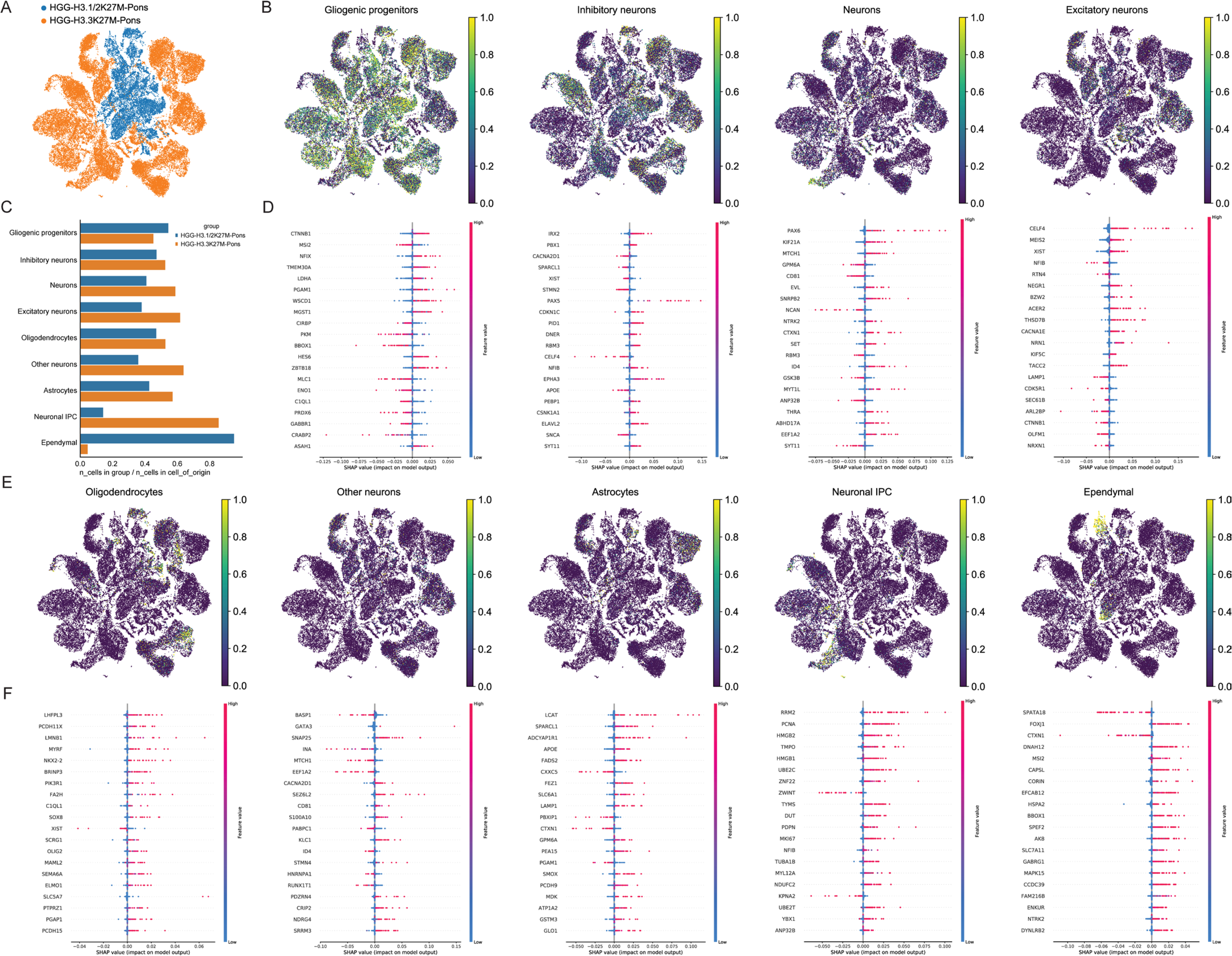
Characterization of developmental-like cell states in DMG scRNA-Seq data. **(A)** Tumor subgroups are shown for DMG scRNA-seq dataset. **(B)** Developmental cell type probability scores are shown for Gliogenic Progenitor, Inhibitory Neurons, Neurons, and Excitatory Neurons cell types. **(C)** Barplots showing the distribution of DMG cells within each tumor subgroup mapped to individual developing cell types. **(D)** The figure displays the results of SHAP analysis, showing the top impactful genes from each cell type Gliogenic Progenitor, Inhibitory Neurons, Neurons, and Excitatory Neurons respectively, in our training dataset. **(E)** Developmental cell type probability scores are shown for Oligodendrocytes, Other neurons, Astrocytes and Neuronal IPC, and Ependymal cell types **(F)** The figure displays the results of SHAP analysis, showing the top impactful genes from each cell type Oligodendrocytes, Other neurons, Astrocytes and Neuronal IPC and Ependymal respectively, in our training dataset.

Additionally, SHAP analysis identified that *FOXJ*, a well-known ependymal transcription factor^39^, contributed to the mapping of H3.1/2 tumor cells to ependymal origins, while NFIB, a recognized transcription factor for neuronal progenitors^40^, contributed to the assignment of H3.3 tumor cells to neuronal IPCs. Next, we predicted the cell age of each identified developmental cell type within DMG tumor cells using our pretrained cell age regressor models **(Figure S9).** H3.3 tumor cells mapping to neuronal IPCs mostly correspond to the earlier weeks in development, whereas H3.1/2 tumor cells mapping to ependymal-like cells mostly correspond to the later weeks in development (**Figure S9**). The DEGs between tumor cells and their respective cells of origin, as well as the DEGs between the cell of origin and the subsequent developmental cell type, are listed in **Table S3 (Figure S10-27)**.

### Application of COORS algorithm on inhouse scRNA-Seq glioma data

Next, we applied the COORS algorithm to our inhouse glioma scRNA-Seq data, containing 21 samples and approximately ~234K cells in total (**Figure 4A**)^41^. Using COORS algorithm, we have used the pre-trained cell type and cell age models, derived from Jessa et al. developing mouse forebrain^6^ and three human developing brain scRNA-Seq datasets from Zeng et al^15^, Polioudakis et al^128^ and Bhaduri et al^16^ to map adult glioma cells (**Figure 4B-D**). Pretrained models from both mouse forebrain developmental dataset^12^ and the dataset by Bhaduri et al.^41,42^ maps IDH^WT^ subgroup tumor cells to the radial glia (RG) cells. RG cells are neural stem cells found in the developing human brain, particularly during the embryonic stages of brain development^42,43^ and alterations in their regulatory pathways or genetic mutations can lead to their transformation into glioma stem-like cells (GSCs), which are thought to drive tumor initiation and progression in glioblastoma^44–46^. In another dataset on the developing human brain by Zeng et al.^15^, IDH^WT^ tumor cells are again mapped to Neural Stem Cells (NSC-cluster 12). Poliodakis et al.^41,42^ dataset more specifically maps IDH^WT^ tumor cells to ventricular zone radial glia (vRG) which are known to reside in the ventricular zone (VZ) of the developing brain^42,43^. On the other hand, IDH^Mut^ cells consistently maps to oligodendrocytes, OPC and also to neuronal subtypes using multiple pretrained models from Bhaduri et al.^16^, Poliodakis et al.^12^, Jessa et al.^6^ datasets. In another dataset on the developing human brain by Zeng et al.^46,47^, IDH^Mut^ tumor cells mostly mapped to specific populations of GABA cells in the developing brain (GABA-cluster 9) derived from the ectoderm layer. Previous studies also suggest that GABAergic neurons and OPCs are derived from common neurodevelopmental origins; predominantly, they both originate from Nkx2.1-expressing precursors located in the medial, lateral, and caudal ganglionic eminences^47,48^. Moreover, GABAergic neurons and OPCs converge at a shared transcriptional state with expression of *OLIG2*^49^, *GABARs*^50^, and *PDGFRA*^51^. In our recent study, we demonstrated that a subset of IDH^Mut^ glioma cells fire single, short action potentials (APs) and are defined by mixed characteristics of GABAergic neurons and OPC^52^.

**Figure 4.**
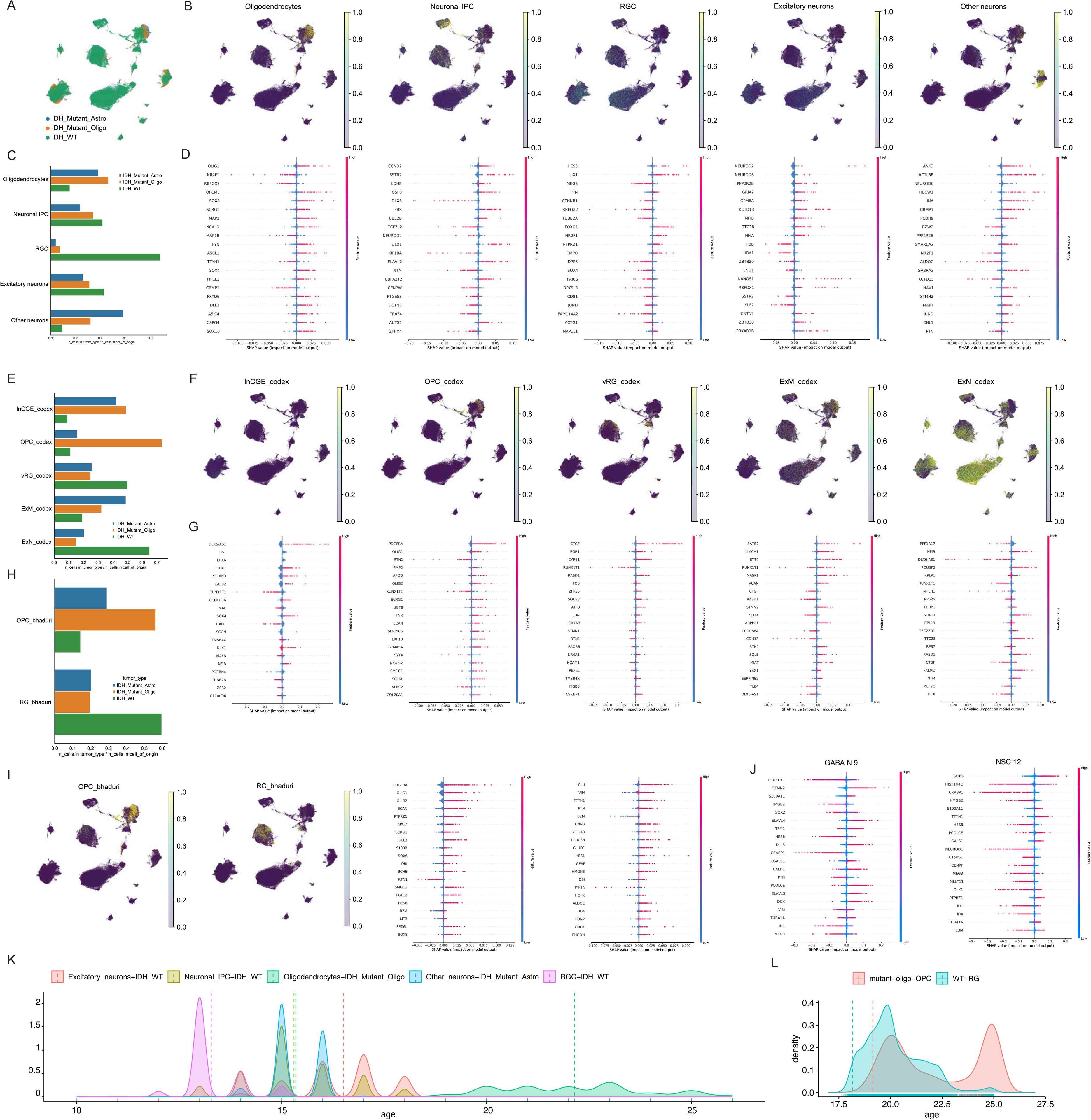
Characterization of developmental-like cell states in glioma scRNA-Seq data. **(A)** Tumor subgroups are shown for inhouse glioma scRNA-seq dataset. **(B)** Developmental cell type probability scores are shown for Oligodendrocytes, Neuronal IPC, RGC, Excitatory Neurons and Other neurons predicted from Jessa et al pretrained models^14^. **(C)** Barplots showing the distribution of glioma cells within each tumor subgroup mapped to individual developing cell types in Jessa et al dataset^14^. **(D)** The figure displays the results of SHAP analysis, showing the top impactful genes from each cell type, Oligodendrocytes, Neuronal IPC, RGC, Excitatory Neurons and Other neurons respectively, using Jessa et al pretrained models^14^. **(E)** Barplots showing the distribution of glioma cells within each tumor subgroup mapped to individual developing cell types predicted from Poliudokis et al. pretrained models^12^. **(F)** Developmental cell type probability scores are shown for InCGE (caudal ganglionic eminence derived interneurons), OPC, vRG, ExM (migrating excitatory neuron) and ExN (newborn excitatory neuron) cell types predicted from Poliudokis et al. pretrained models^12^. **(G)** The figure displays the results of SHAP analysis, showing the top impactful genes from each cell type in InCGE, OPC, vRG, ExM and ExN celltypes using Poliudokis et al. pretrained models^12^. **(H)** Barplots showing the distribution of glioma cells within each tumor subgroup mapped to individual developing cell types in Bhaduri et al. dataset^16^. **(I)** Developmental cell type probability scores are shown for OPC, RG cell types in Bhaduri et al. dataset^16^ and SHAP analysis, showing the top impactful genes from each cell type, OPC, RG respectively, using Bhaduri et al. pretrained models^16^. **(J)** The figure displays the results of SHAP analysis, showing the top impactful genes from each cell type, GABA N 9 and NSC 12 respectively, using Zeng et al. pretrained models^15^. **(K)** Distribution of age mapping within each tumor subgroup and their respective mapped cell of origin pairs from Jessa et al pretrained models^14^. **(L)** Distribution of age mapping within each tumor subgroup and their respective mapped cell of origin pairs from Bhaduri et al. pretrained models^16^.

In addition, we conducted SHAP analysis to extract critical features from our machine-learning neural network models, enabling the identification of developmental-like gene markers specific to glioma for each mapped cell type. SHAP analysis identified that markers commonly associated with oligodendrocyte precursor cells (OPCs) such as *OLIG1*^53^, *SOX10*^54^, *PDGFRA*^51^ *NKX2-2*^55^ and *OLIG2*^56,57^ predominantly contributed to the mapping of IDH mutant tumor cells to OPCs. GABAA receptor subunit *GABRA2*^58^, known to be also expressed in oligodendrocytes contributed to the mapping of neuronal subtypes in IDH^Mut^ using Jessa et al.^6^ pretrained model. In the dataset from Zeng et al.^15^, GABA-cluster 9 exhibits a high expression of genes such as *STMN2*, *ELAVL4*, *ELAVL3*, and *DCX*, all of which are known to have crucial role in neuronal development and differentiation, predominantly contributed to the mapping of IDH^Mut^ tumor cells to neuronal subtypes. SHAP analysis identified that markers commonly associated with neural stem cells and radial glia cells *HES5*^59^*and HES1*^59^, contributed to the mapping of IDH^WT^ tumor cells to radial glia cells. NSC-cluster 12 exhibits high expression of genes such as *SOX2*, and *TTYH1*, all of which are known markers of neural stem cells^60,61^, contributing to the mapping of IDH^WT^ tumor cells to NSCs.

To estimate the cell age of each identified developmental cell type within glioma tumor cells, we applied our pre-trained cell age regressor models (**Figure 4A-C**). Age mapping was performed exclusively on the datasets from Bhaduri et al.^16^, Zeng et al.^15^, and Jessa et al.^6^ due to their wide range of developmental age data. Notably, both Bhaduri et al.^16^ and Jessa et al.^6^ datasets’ models consistently revealed that IDH^Mut^ cells corresponding to OPCs exhibit a bimodal age distribution, indicating stages early and late in development. In contrast, IDH^WT^ cells aligning with RG mostly represent earlier developmental weeks. The DEGs between tumor cells and their respective cells of origin, as well as the DEGs between the cell of origin and the following developmental cell type, are listed in **Table S4-5 (Figure S28-51)**.

## Discussion

Here we presented a hierarchical machine learning-based approach, named COORS, for the identification and characterization of tumor cells that exhibit gene expression patterns reminiscent of early developmental stages in the brain. COORS achieves this by employing NNMs trained on diverse scRNA-Seq datasets from developing human brain tissues. We applied our method to predict developmental-like cells in various brain cancer datasets, including MB, DMG, and glioma, with validation against well-characterized MB data. COORS identified vRG developmental cells within IDH^WT^ glioma cells whereas OPC and neuronal-like cells in IDH^Mut^. Interestingly, IDH^Mut^ subgroup cells that map to OPC show bimodal distributions, that are both early and late weeks in development, while IDH^WT^ subgroup cells mapping to RG mostly correspond to the earlier weeks in development. Furthermore, COORS offers a valuable resource by providing information on the DEGs between tumor cells and their respective cells of origin, as well as between the cell of origin and the subsequent developmental cell type. These DEGs hold promise as potential therapeutic targets, offering new avenues for the development of targeted therapies for brain tumors. In conclusion, the development and application of COORS represent a significant advancement in our ability to accurately annotate developmental-like cell states in brain cancer datasets and potentially extend this approach to other cancer types.

In the past, efforts to induce the differentiation of cancer cells into more mature, less aggressive cell types, without damaging normal cells were met with limited success in solid tumors, likely due to insufficient understanding of the precise progenitor cells involved^62,63^. However, advancements in our comprehension of specific time points and cell types have paved the way for a more nuanced approach. By examining the subsequent steps in the lineage, such as the differentiation of OPCs into mature oligodendrocytes, we can identify key genes involved in this process. For instance, oligodendrocytes are characterized by decreased proliferative capacity compared to OPCs, suggesting a regulatory role for certain genes in cell fate determination. Targeting genes like *OLIG2*, which maintains OPC identity, while activating those involved in mature oligodendrocyte function, such as myelin basic protein (*MBP*), holds promise for directing OPCs toward differentiation into oligodendrocytes. This approach presents a potential avenue for differentiation therapy, wherein manipulating gene expression could drive tumor cells toward a more benign phenotype, offering a novel strategy for cancer treatment.

Overall, our approach relies on a separate model for each origin-like cell type. While one could use a single model for assignment of all cell types jointly (e.g. multitask learning) to make use of all data at once, our experiments did not show significant benefit of one model compared to building simpler single models for each cell type. We hypothesize that this may be inherently a result of data size requirements and complexities of jointly learning hundreds of origin-like cell types from unbalanced datasets. In addition, our approach provides more flexibility for selecting biologically meaningful models of origin-like cell types in different tumors.

## Methods

### Data preprocessing

We conducted the standard pipeline of single-cell RNA sequencing data preprocessing for both reference and testing datasets using Scanpy 1.7.2 in Python 3.6.8. For cell of origin classification, the preprocessing of reference and testing data were performed starting from the whole datasets. On the other hand, for cell age regression, we first grouped the reference and testing datasets by cells of origin and then preprocessed each group separately. Each cell was normalized to have the same total read count and the matrices were transformed into natural logarithm domain. We annotated the top 2,000 highly variable genes in reference dataset, scaled both datasets to unit variance and zero mean, and truncated to 10. We kept all the other parameters in default values.

A common gene set is needed between the reference and testing datasets as they will be fed into one same model. We took the union of reference marker genes and highly variable genes to intersect with the testing genes as the common gene set and trimmed both datasets. The volumes of reference cell types vary. We first excluded cell types with number of cells fewer than 20. To balance the number of cells among the rest cell types, we set as baseline the cell type that is the 25% quantile in terms of numbers of cells and randomly subsampled the others to this baseline. Those cell types with number of cells fewer than the baseline were not subsampled.

We randomly split each reference cell type into two subsets, one with 80% cells for model training and the other with 20% cells for model validation. The training subsets of all the cell types were concatenated as training data, the validation subsets as validation data.

We conducted one-hot encoding of cell types in both training and validation data. We scaled training, validation, and testing data into the range from 0 to 1 using Scikit-learn 0.24.2.

### MLP-based prediction model

We developed multilayer perceptron networks for cell of origin classification and cell age regression.

#### Cell of origin classifier

The cell of origin classifier has one input layer, variable numbers of hidden layers, and one output layer. The input layer has the same number of nodes as the input genes. The number of hidden layers varies from one to four, and the number of nodes in one hidden layer is set to be 256, 128, 64, or 32, which is determined after hyperparameter optimization. Following the dense connection within each hidden layer, there are batch normalization, activation, and dropout functions. We use the popular Rectified Linear Unit (ReLU) for hidden layer activation and set dropout rate to be 0.1 or 0.2. The output layer uses Softmax activation function so that each node outputs a non-negative value smaller than 1 and all the values sum up to 1. Therefore, each output corresponds to the probability of one cell type. We compile the model using categorical crossentropy as loss function, Adam as optimizer, and accuracy as metrics.

#### Cell age regressor

Similar as cell of origin classifier, cell age regressor consists of one input layer, a group of hidden layers, and one output layer. While the input layer and hidden layers are structurally the same as cell of origin classifier, the output layer of cell age regressor has only one node with Sigmoid activation function that corresponds to the predicted cell age. The model is compiled using mean squared error as loss function, Adam as an optimizer, and loss as metrics. Since more than one cell age regressors exist corresponding to each cell of origin classifier, these regressors can have specific hyperparameters of hidden layers that are not necessarily the same.

#### Model training prerequisites

We implemented cell of origin classifier and cell age regressor using Keras 2.6.0 with Tensorflow 2.6.2 as backend in Python 3.6.8. Prerequisite packages for data preprocessing and model training include Numpy 1.19.5, Pandas 1.1.5, Scanpy 1.7.2, Anndata 0.7.8, Scipy 1.5.4, and Scikit-learn 0.24.2.

#### Hyperparameter optimization

We systematically optimized hyperparameters of cell of origin classifier and cell age regressor using grid search cross validation implemented by Scikit-learn 0.24.2, focusing on tuning the number of hidden layers and nodes, dropout rate of hidden layers, and learning rate of the optimizer. For each model, we varied the number of hidden layers from one to four and the number of nodes in each layer that could be 256, 128, 64, or 32. We followed convention to use Rectified Linear Unit (ReLU) as activation function in hidden layers. Training epochs were fixed to be 100 and batch size 32 as they did not show significant affections in our case. Along with the iteration of every possible hidden layer structure, we explored dropout rate of 0.1 or 0.2 and learning rate of 0.1, 0.01, 0.001, or three decaying learning rates that were scheduled to exponentially reduce during model training based on initial rate of 0.1 or 0.01, final rate of 0.01 or 0.001, training epochs, and batch size.

## Source Code Availability

Source code of COORS is publicly available at https://github.com/Su-Wang-UTH/COORS

## Supporting information

Supplemental Meterial

