## Supplemental Meterial for "Inferred Developmental Origins of Brain Tumors from Single Cell RNA-Sequencing Data"

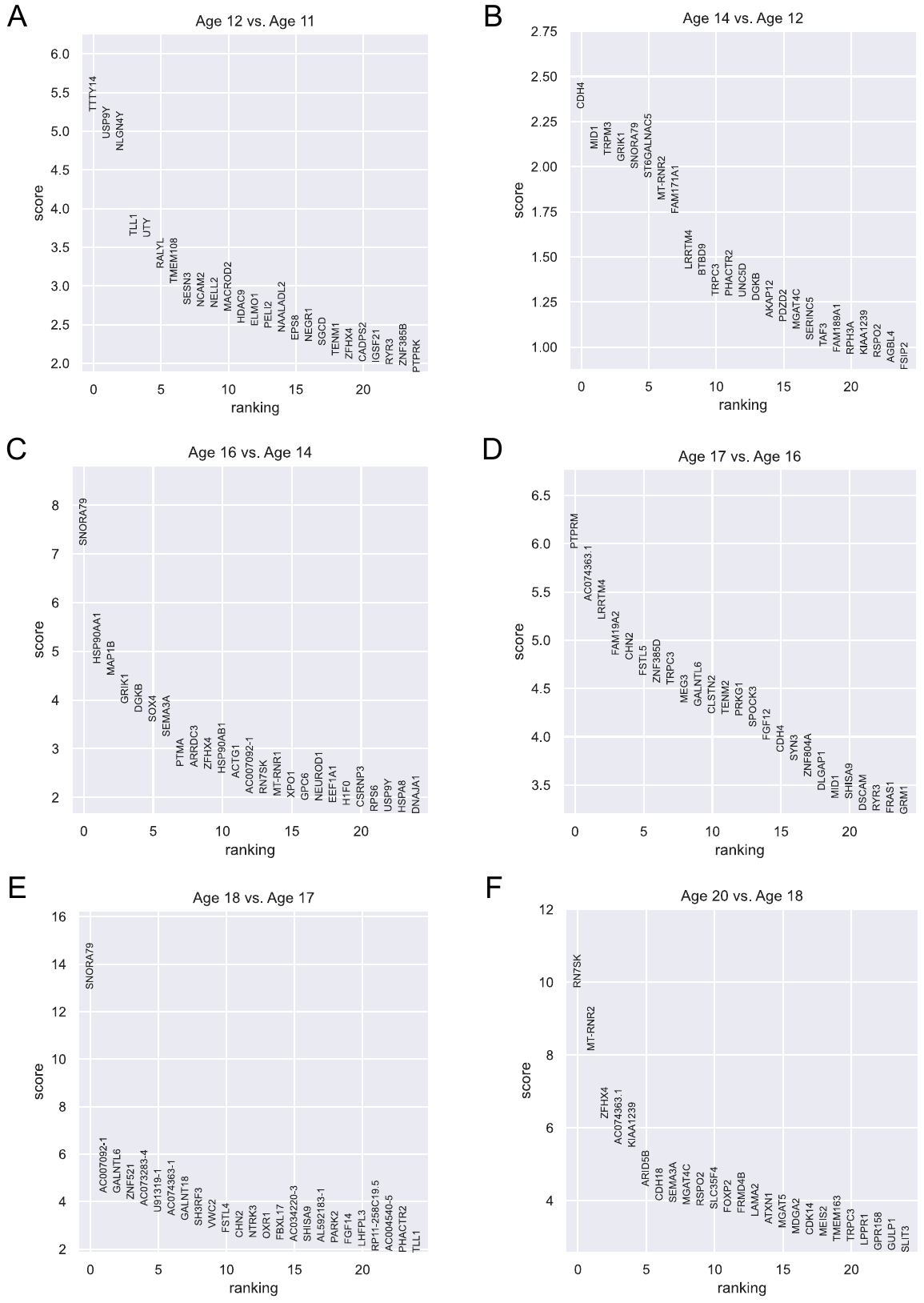

**Figure S1. Differentially expressed genes (DEGs) between adjacent age groups of CN-UBC in Aldinger’s dataset.** Differential expression analysis using Wilcoxon rank-sum is conducted for each pair of adjacent age groups. Genes are ranked according to the z-score underlying the p-value computation and the top 25 rankings are displayed. (A) DEGs between 12 PCW and 11 PCW. (B) DEGs between 14 PCW and 12 PCW. (C) DEGs between 16 PCW and 14 PCW. (D) DEGs between 17 PCW and 16 PCW. (E) DEGs between 18 PCW and 17 PCW. (F) DEGs between 20 PCW and 18 PCW.

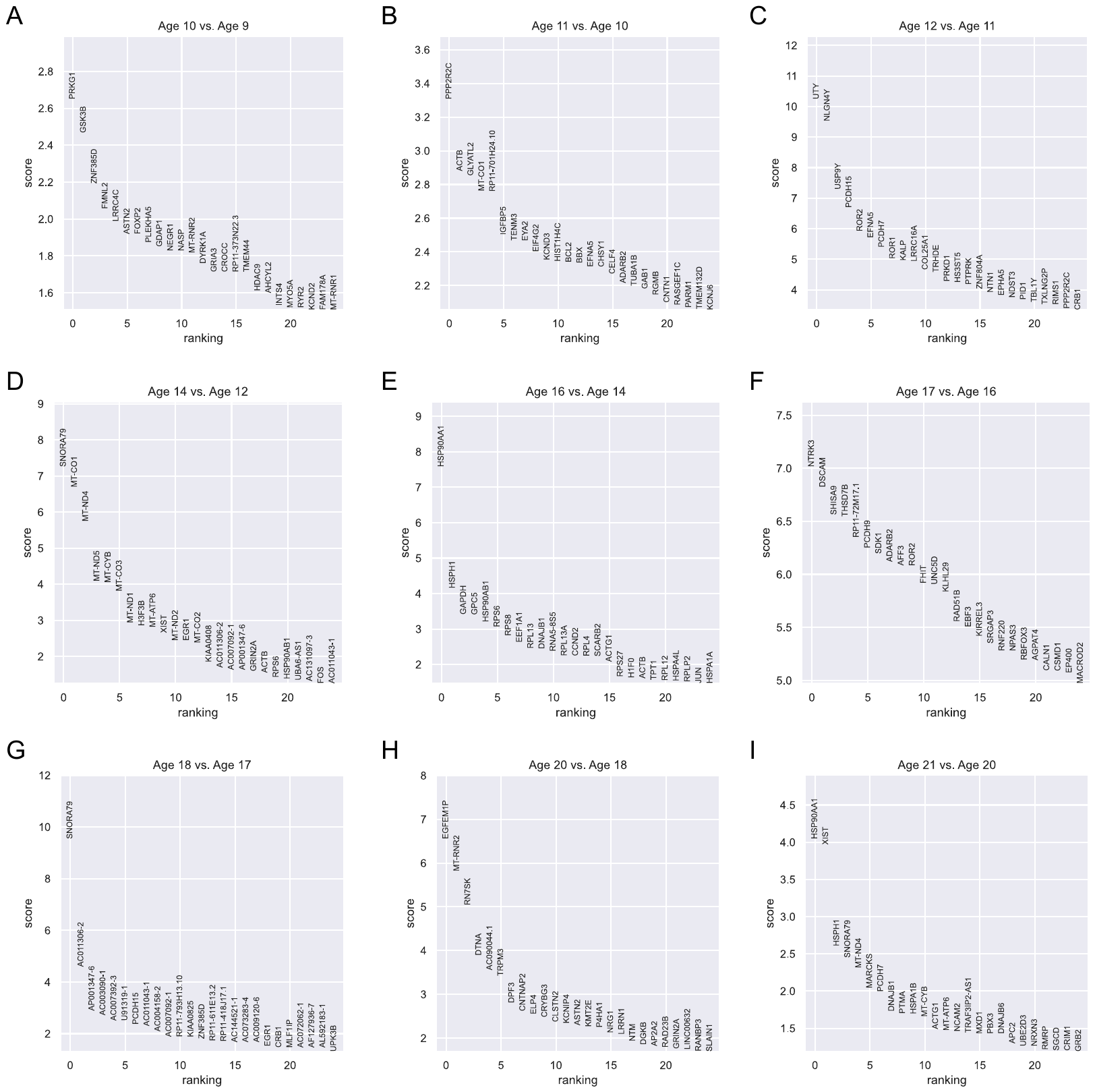

**Figure S2. Differentially expressed genes (DEGs) between adjacent age groups of GCP in Aldinger’s dataset.** Differential expression analysis using Wilcoxon rank-sum is conducted for each pair of adjacent age groups. Genes are ranked according to the z-score underlying the p-value computation and the top 25 rankings are displayed. (A) DEGs between 10 PCW and 9 PCW. (B) DEGs between 11 PCW and 10 PCW. (C) DEGs between 12 PCW and 11 PCW. (D) DEGs between 14 PCW and 12 PCW. (E) DEGs between 16 PCW and 14 PCW. (F) DEGs between 17 PCW and 16 PCW. (G) DEGs between 18 PCW and 17 PCW. (H) DEGs between 20 PCW and 18 PCW. (I) DEGs between 21 PCW and 20 PCW.

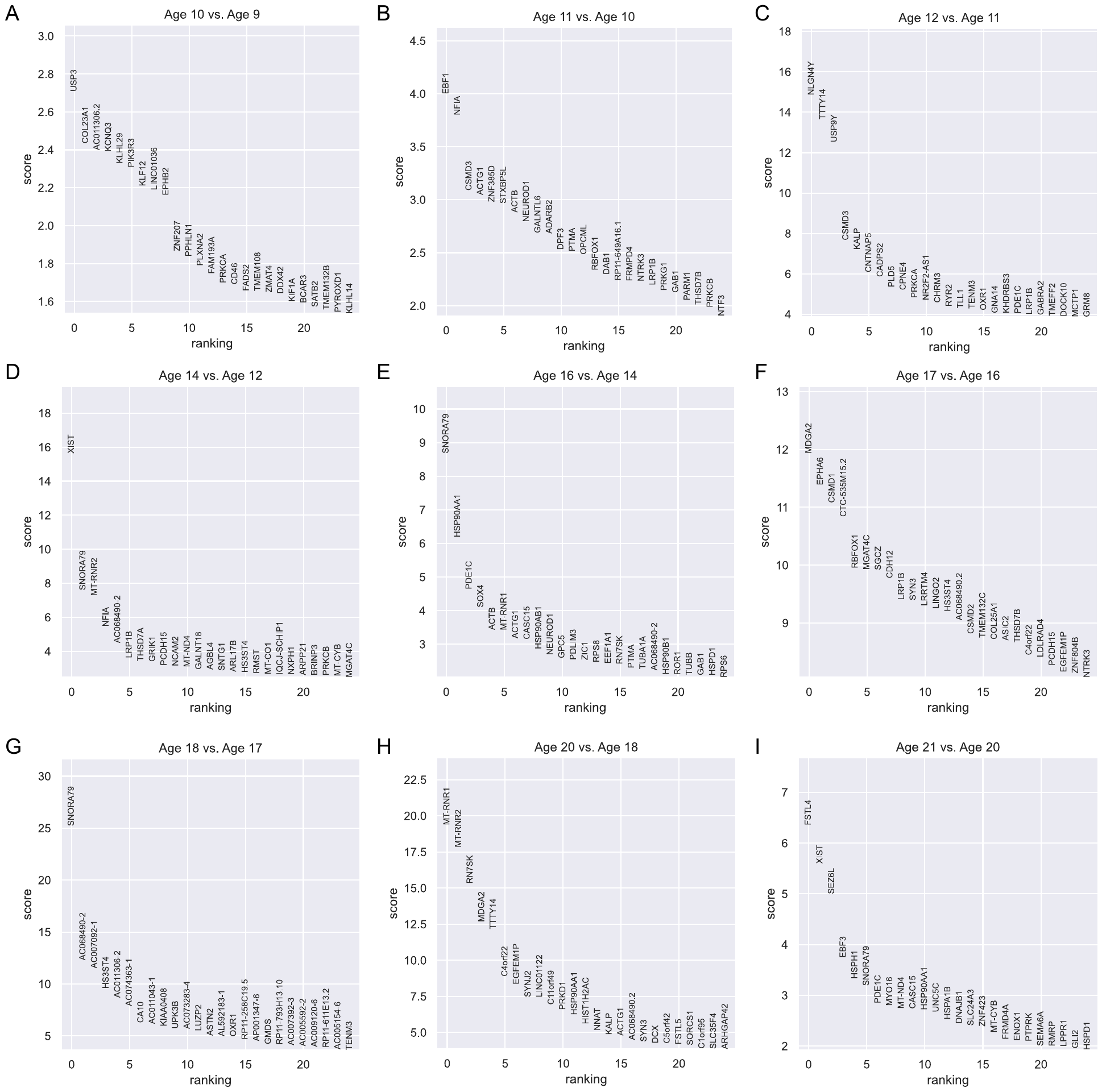

**Figure S3. Differentially expressed genes (DEGs) between adjacent age groups of GN in Aldinger’s dataset.** Differential expression analysis using Wilcoxon rank-sum is conducted for each pair of adjacent age groups. Genes are ranked according to the z-score underlying the p-value computation and the top 25 rankings are displayed. (A) DEGs between 10 PCW and 9 PCW. (B) DEGs between 11 PCW and 10 PCW. (C) DEGs between 12 PCW and 11 PCW. (D) DEGs between 14 PCW and 12 PCW. (E) DEGs between 16 PCW and 14 PCW. (F) DEGs between 17 PCW and 16 PCW. (G) DEGs between 18 PCW and 17 PCW. (H) DEGs between 20 PCW and 18 PCW. (I) DEGs between 21 PCW and 20 PCW.

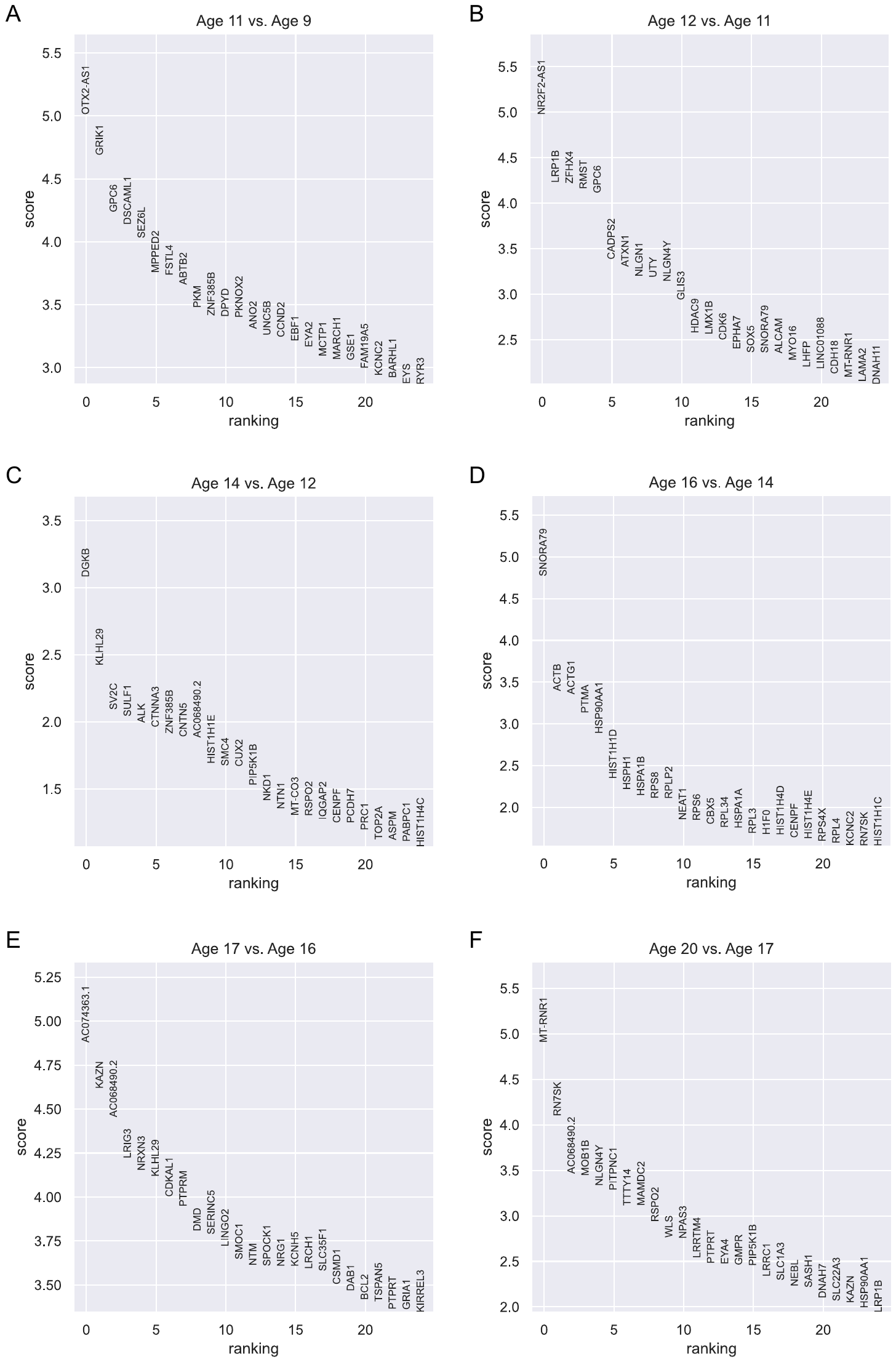

**Figure S4. Differentially expressed genes (DEGs) between adjacent age groups of RL in Aldinger’s dataset.** Differential expression analysis using Wilcoxon rank-sum is conducted for each pair of adjacent age groups. Genes are ranked according to the z-score underlying the p-value computation and the top 25 rankings are displayed. (A) DEGs between 11 PCW and 9 PCW. (B) DEGs between 12 PCW and 11 PCW. (C) DEGs between 14 PCW and 12 PCW. (D) DEGs between 16 PCW and 14 PCW. (E) DEGs between 17 PCW and 16 PCW. (F) DEGs between 20 PCW and 17 PCW.

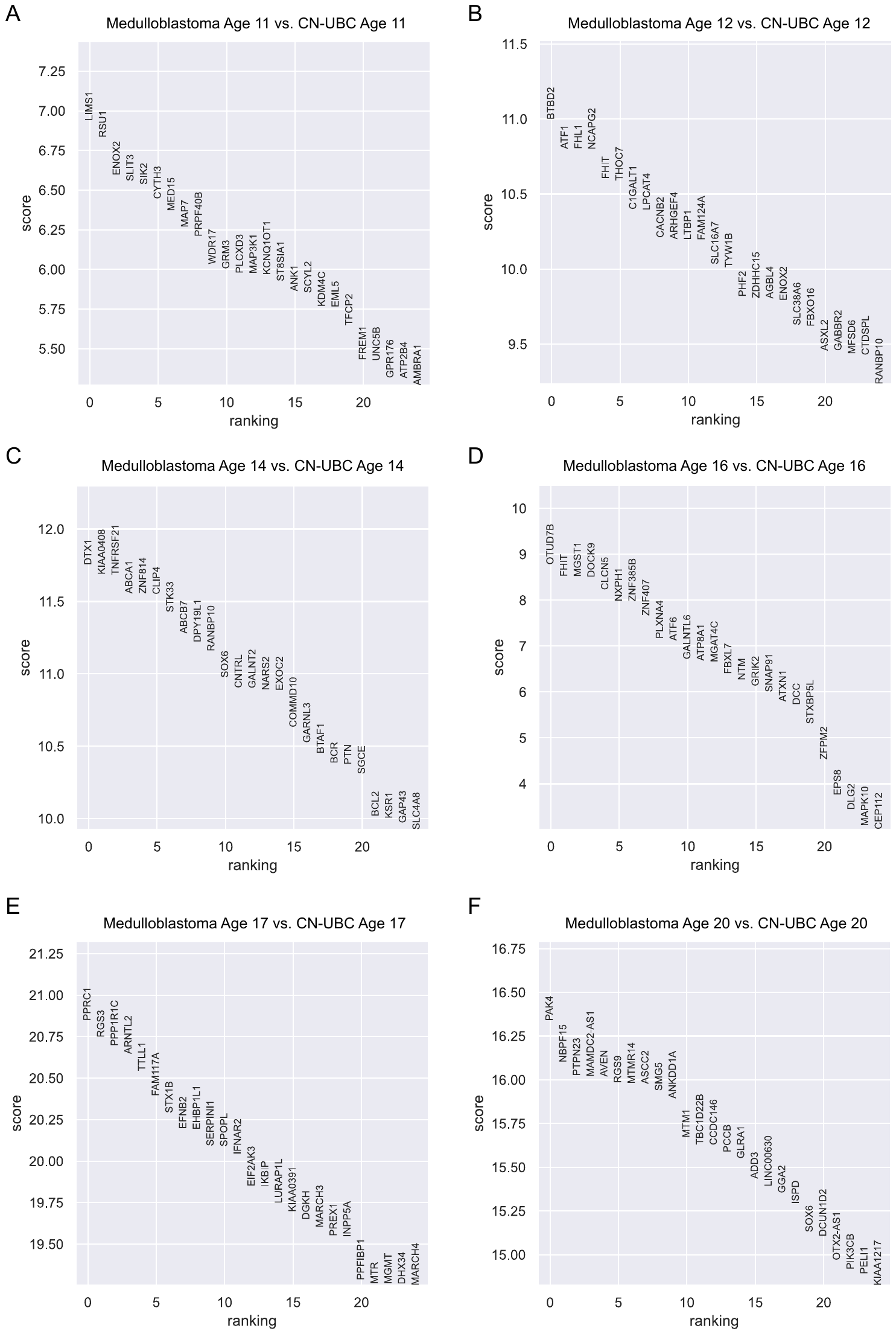

**Figure S5. Differentially expressed genes (DEGs) between medulloblastoma and CN-UBC in Aldinger’s dataset.** Differential expression analysis using Wilcoxon rank-sum is conducted for each age group. Genes are ranked according to the z-score underlying the p-value computation and the top 25 rankings are displayed. (A) DEGs for 11 PCW. (B) DEGs for 12 PCW. (C) DEGs for 14 PCW. (D) DEGs for 16 PCW. (E) DEGs for 17 PCW. (F) DEGs for 20 PCW.

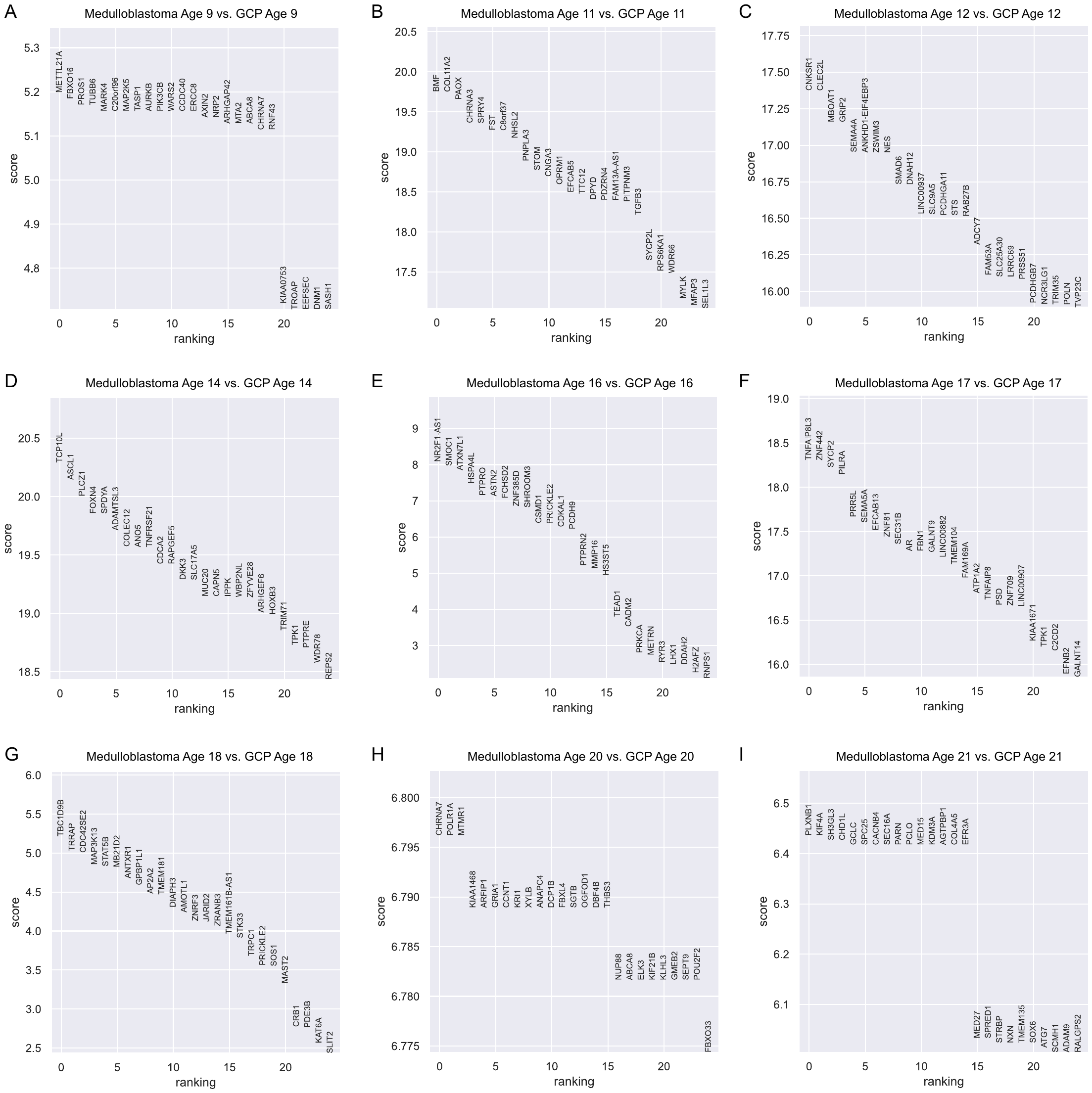

**Figure S6. Differentially expressed genes (DEGs) between medulloblastoma and GCP in Aldinger’s dataset.** Differential expression analysis using Wilcoxon rank-sum is conducted for each age group. Genes are ranked according to the z-score underlying the p-value computation and the top 25 rankings are displayed. (A) DEGs for 9 PCW. (B) DEGs for 11 PCW. (C) DEGs for 12 PCW. (D) DEGs for 14 PCW. (E) DEGs for 16 PCW. (F) DEGs for 17 PCW. (G) DEGs for 18 PCW. (H) DEGs for 20 PCW. (I) DEGs for 21 PCW.

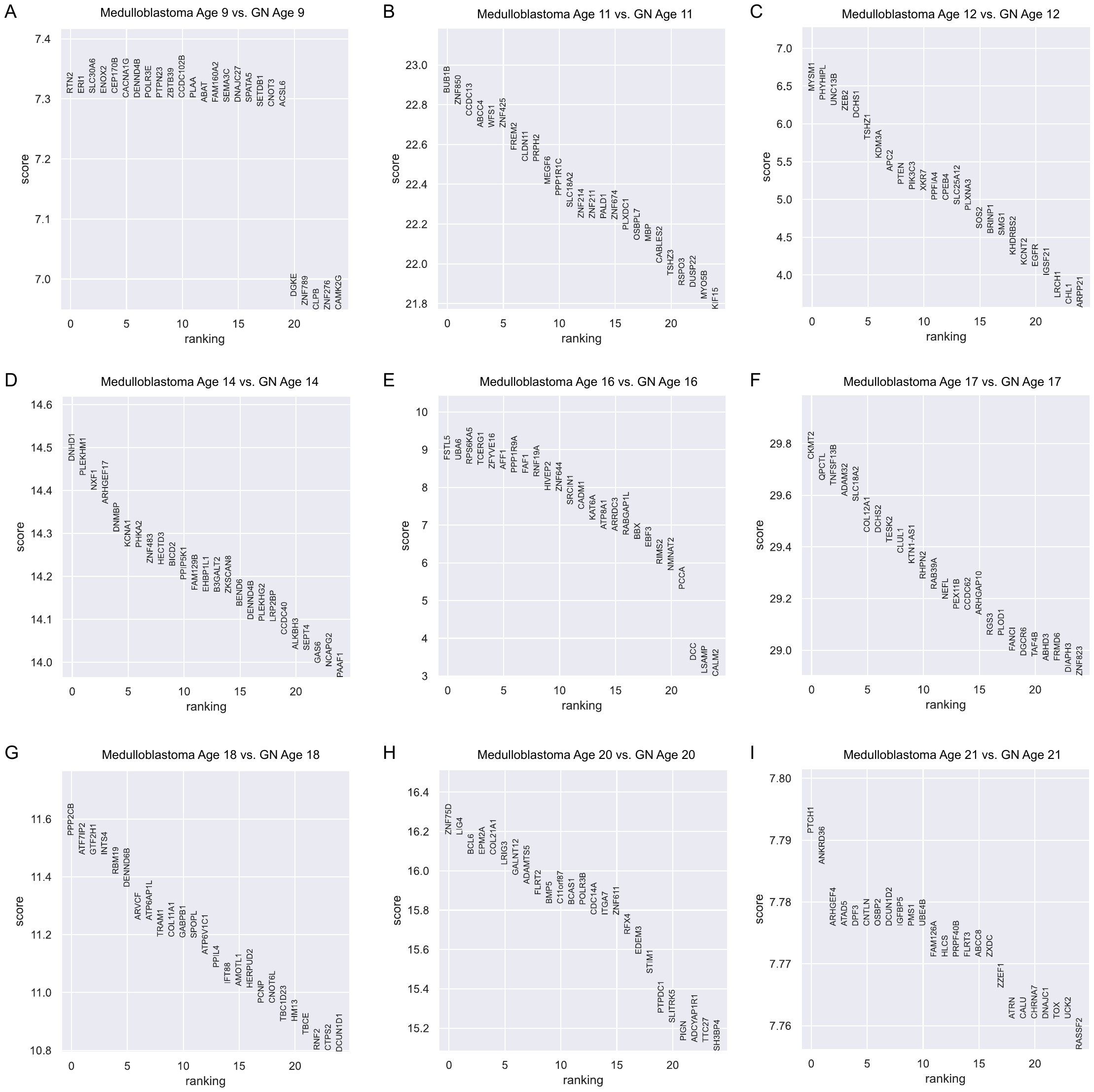

**Figure S7. Differentially expressed genes (DEGs) between medulloblastoma and GN in Aldinger’s dataset.** Differential expression analysis using Wilcoxon rank-sum is conducted for each age group. Genes are ranked according to the z-score underlying the p-value computation and the top 25 rankings are displayed. (A) DEGs for 9 PCW. (B) DEGs for 11 PCW. (C) DEGs for 12 PCW. (D) DEGs for 14 PCW. (E) DEGs for 16 PCW. (F) DEGs for 17 PCW. (G) DEGs for 18 PCW. (H) DEGs for 20 PCW. (I) DEGs for 21 PCW.

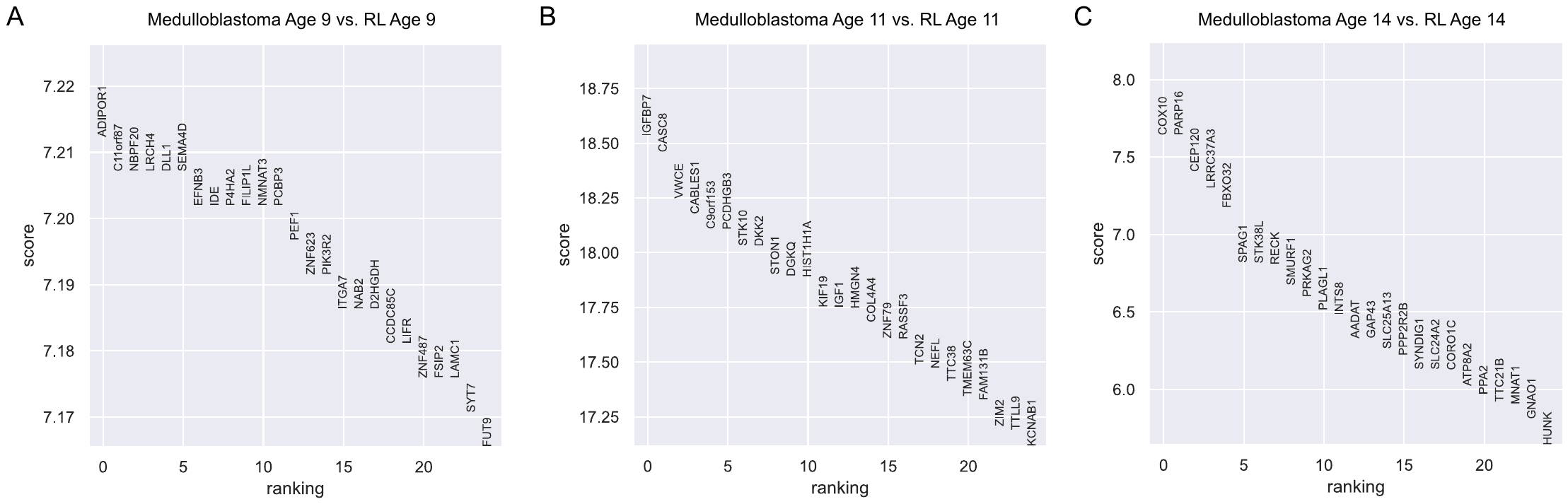

**Figure S8. Differentially expressed genes (DEGs) between medulloblastoma and RL in Aldinger’s dataset.** Differential expression analysis using Wilcoxon rank-sum is conducted for each age group. Genes are ranked according to the z-score underlying the p-value computation and the top 25 rankings are displayed. (A) DEGs for 9 PCW. (B) DEGs for 11 PCW. (C) DEGs for 14 PCW.

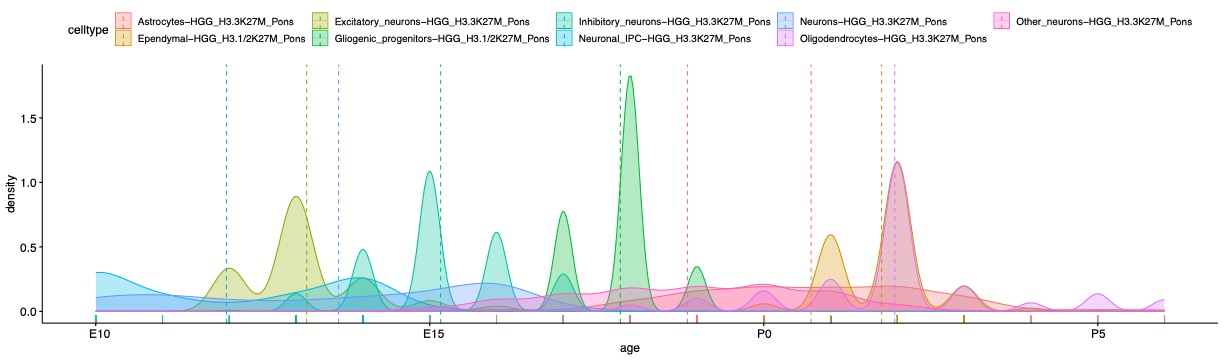

**Figure S9.** Distribution of age mapping within each tumor subgroup and their respective mapped cell of origin pairs.

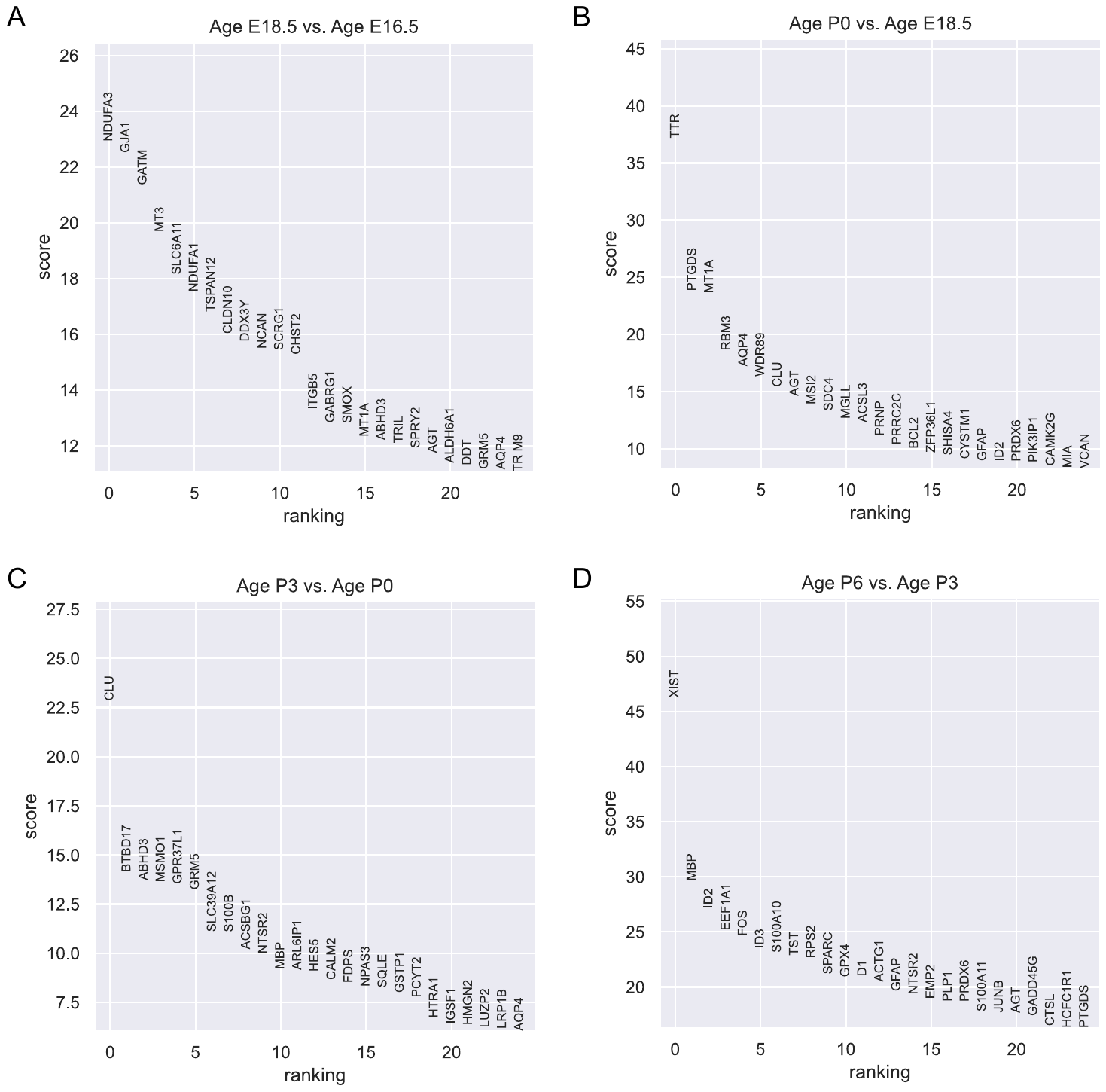

**Figure S10. Differentially expressed genes (DEGs) between adjacent age groups of Astrocytes in Pons dataset.** Differential expression analysis using Wilcoxon rank-sum is conducted for each pair of adjacent age groups. Genes are ranked according to the z-score underlying the p-value computation and the top 25 rankings are displayed. (A) DEGs between E18.5 and E16.5. (B) DEGs between P0 and E18.5. (C) DEGs between P3 and P0. (D) DEGs between P6 and P3.

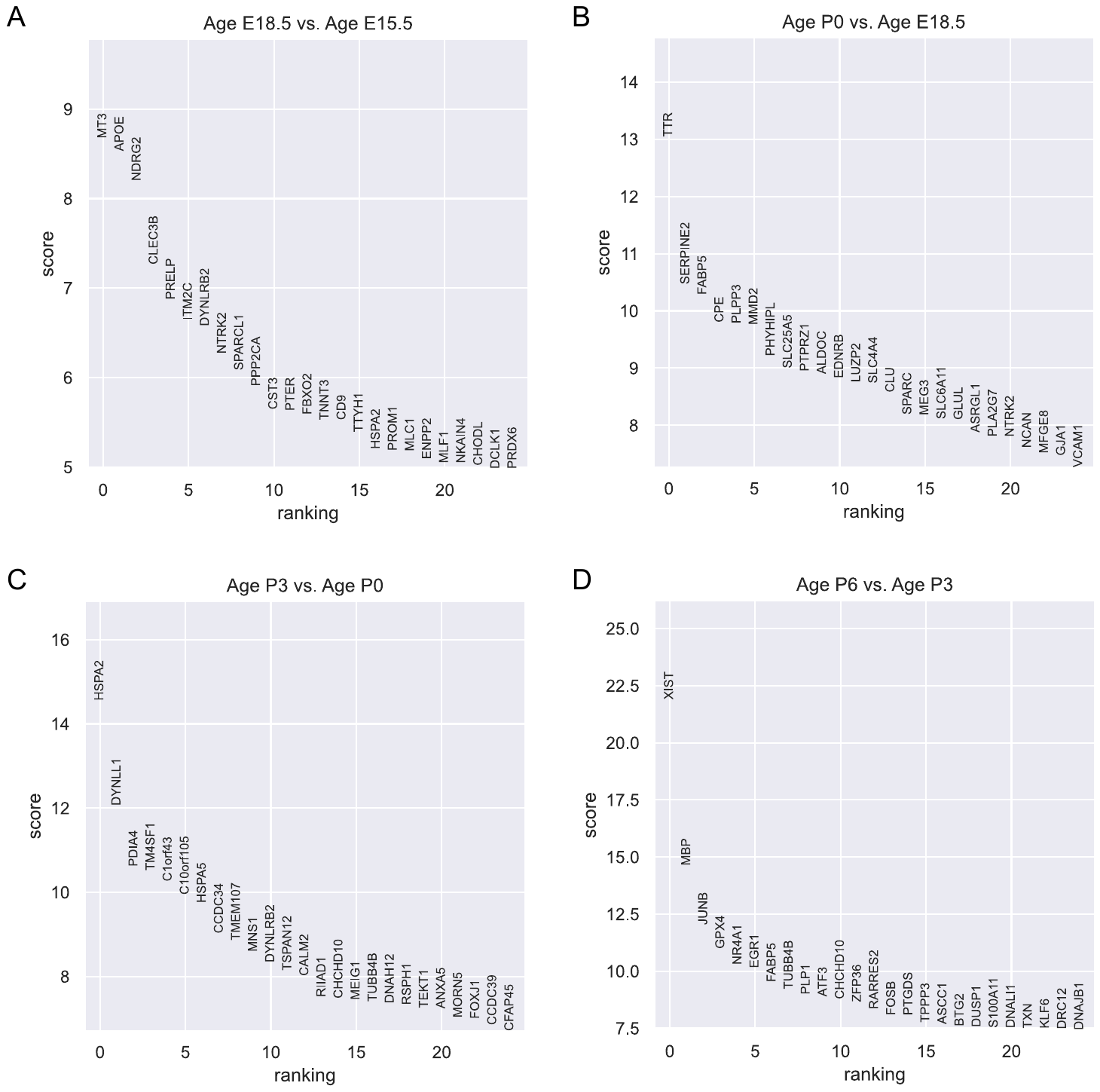

**Figure S11. Differentially expressed genes (DEGs) between adjacent age groups of Ependymal in Pons dataset.** Differential expression analysis using Wilcoxon rank-sum is conducted for each pair of adjacent age groups. Genes are ranked according to the z-score underlying the p-value computation and the top 25 rankings are displayed. (A) DEGs between E18.5 and E15.5. (B) DEGs between P0 and E18.5. (C) DEGs between P3 and P0. (D) DEGs between P6 and P3.

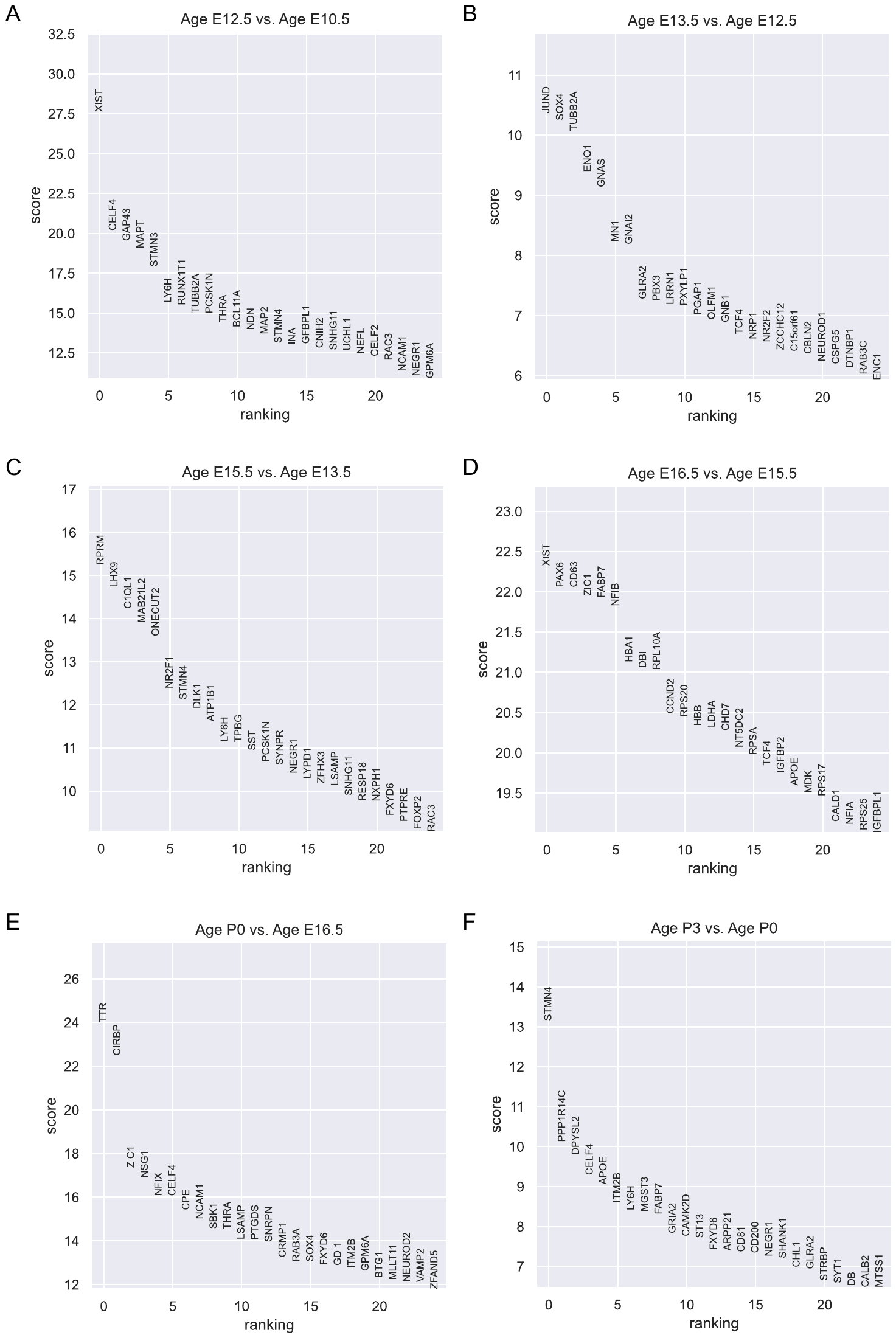

**Figure S12. Differentially expressed genes (DEGs) between adjacent age groups of Excitatory neurons in Pons dataset.** Differential expression analysis using Wilcoxon rank-sum is conducted for each pair of adjacent age groups. Genes are ranked according to the z-score underlying the p-value computation and the top 25 rankings are displayed. (A) DEGs between E12.5 and E10.5. (B) DEGs between E13.5 and E12.5. (C) DEGs between E15.5 and E13.5. (D) DEGs between E16.5 and E15.5. (E) DEGs between P0 and E16.5. (F) DEGs between P3 and P0.

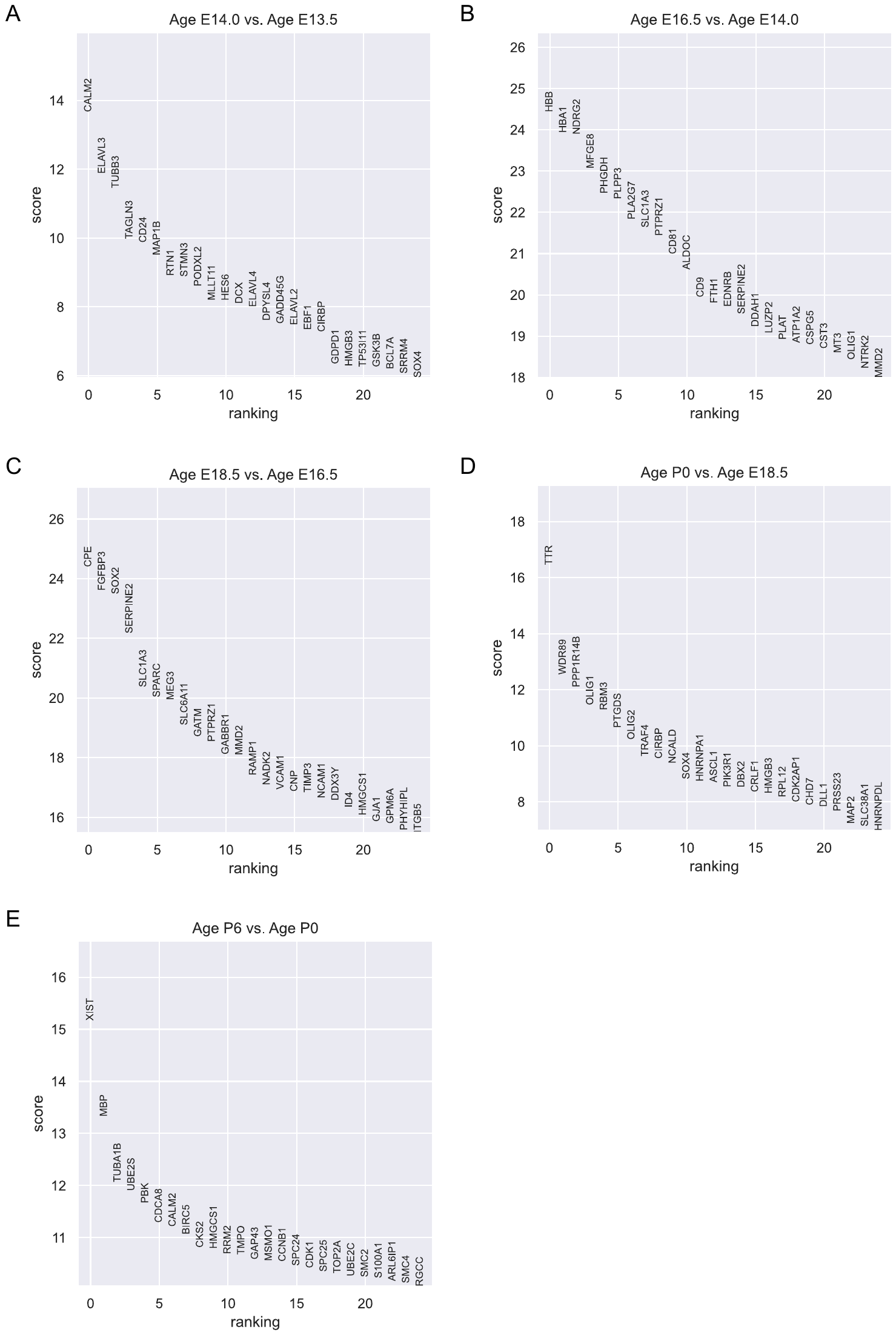

**Figure S13. Differentially expressed genes (DEGs) between adjacent age groups of Gliogenic progenitors in Pons dataset.** Differential expression analysis using Wilcoxon rank-sum is conducted for each pair of adjacent age groups. Genes are ranked according to the z-score underlying the p-value computation and the top 25 rankings are displayed. (A) DEGs between E14.0 and E13.5. (B) DEGs between E16.5 and E14.0. (C) DEGs between E18.5 and E16.5. (D) DEGs between P0 and E18.5. (E) DEGs between P6 and P0.

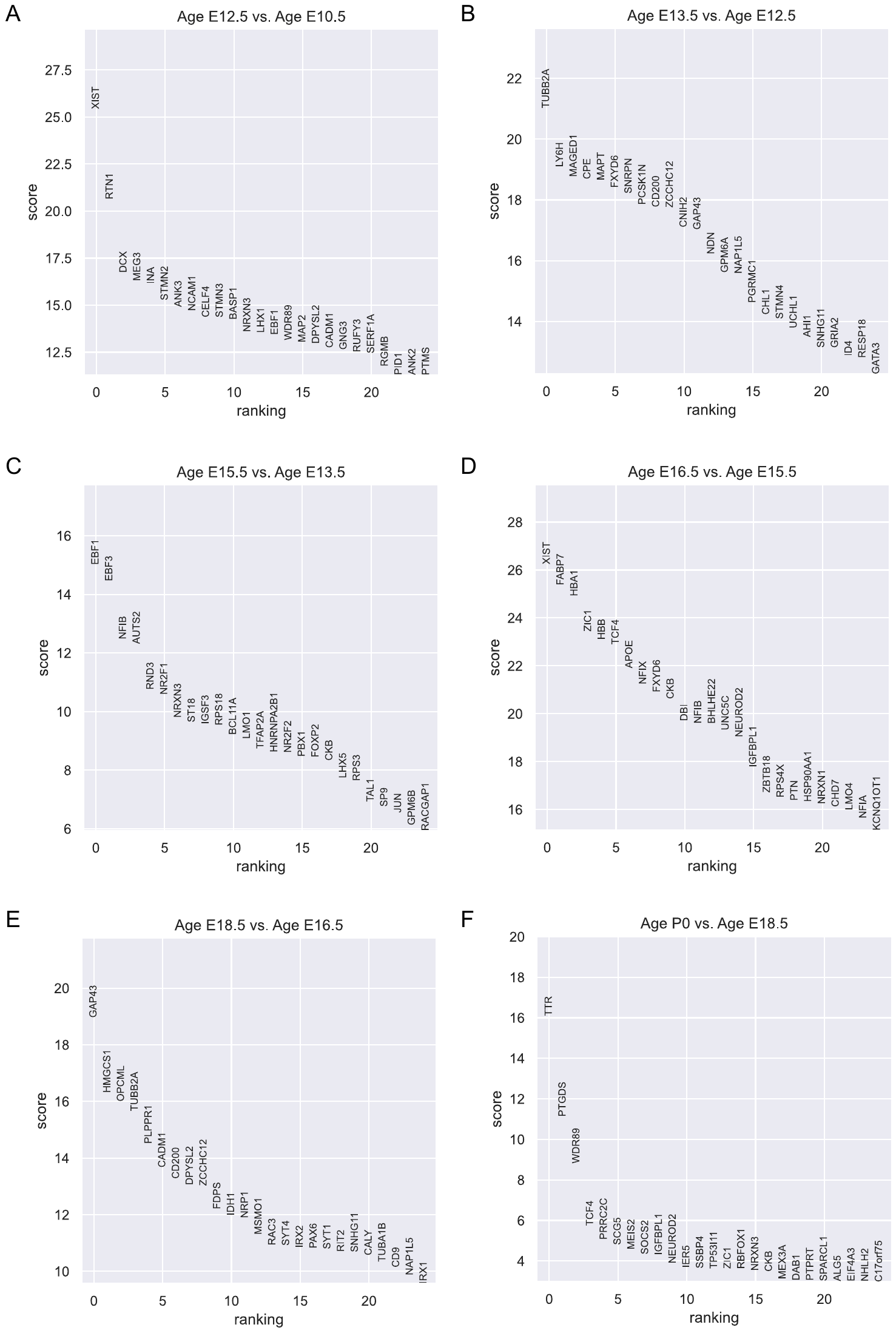

**Figure S14. Differentially expressed genes (DEGs) between adjacent age groups of Inhibitory neurons in Pons dataset.** Differential expression analysis using Wilcoxon rank-sum is conducted for each pair of adjacent age groups. Genes are ranked according to the z-score underlying the p-value computation and the top 25 rankings are displayed. (A) DEGs between E12.5 and E10.5. (B) DEGs between E13.5 and E12.5. (C) DEGs between E15.5 and E13.5. (D) DEGs between E16.5 and E15.5. (E) DEGs between E18.5 and E16.5. (F) DEGs between P0 and E18.5.

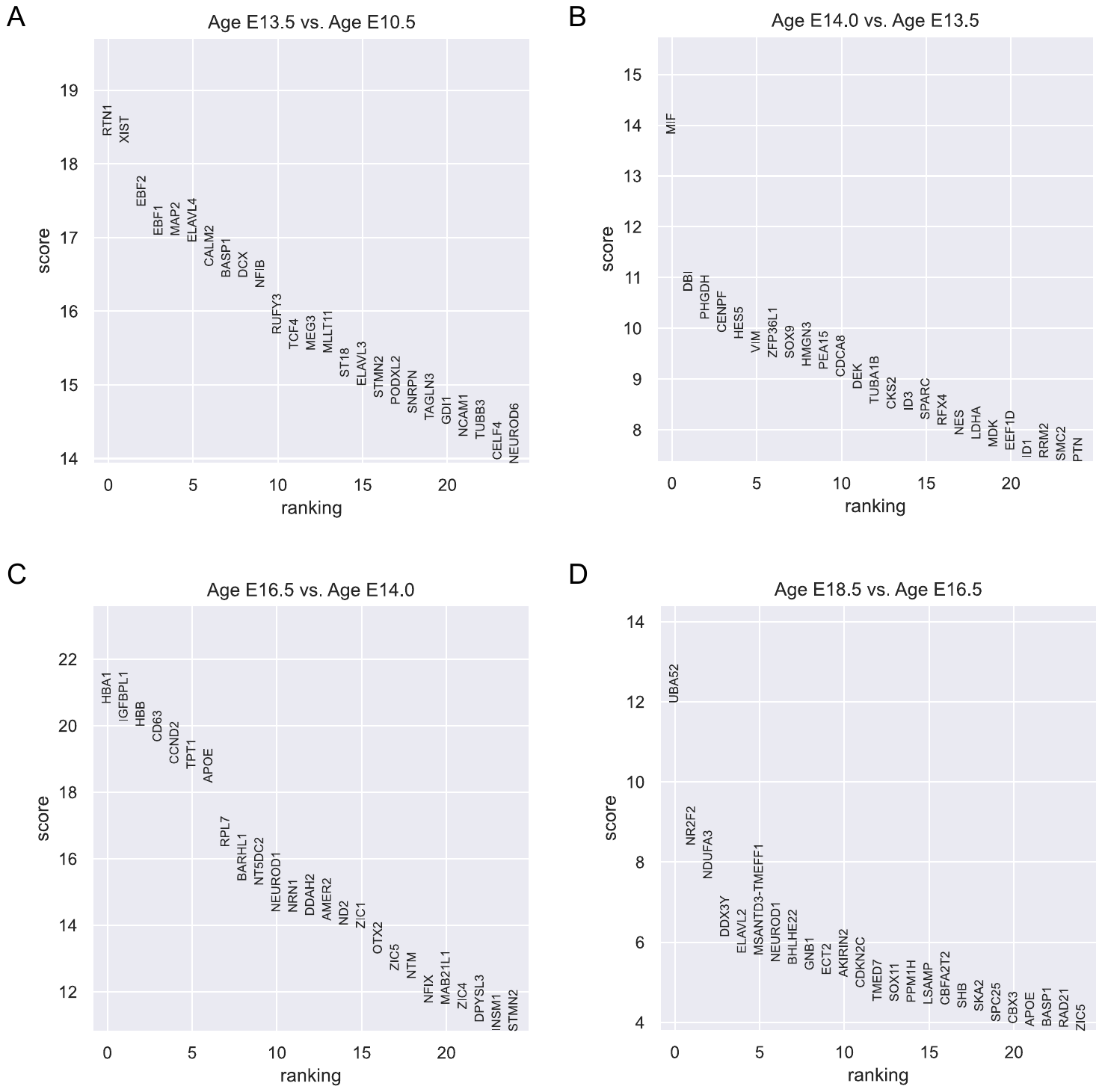

**Figure S15. Differentially expressed genes (DEGs) between adjacent age groups of Neuronal IPC in Pons dataset.** Differential expression analysis using Wilcoxon rank-sum is conducted for each pair of adjacent age groups. Genes are ranked according to the z-score underlying the p-value computation and the top 25 rankings are displayed. (A) DEGs between E13.5 and E10.5. (B) DEGs between E14.0 and E13.5. (C) DEGs between E16.5 and E14.0. (D) DEGs between E18.5 and E16.5.

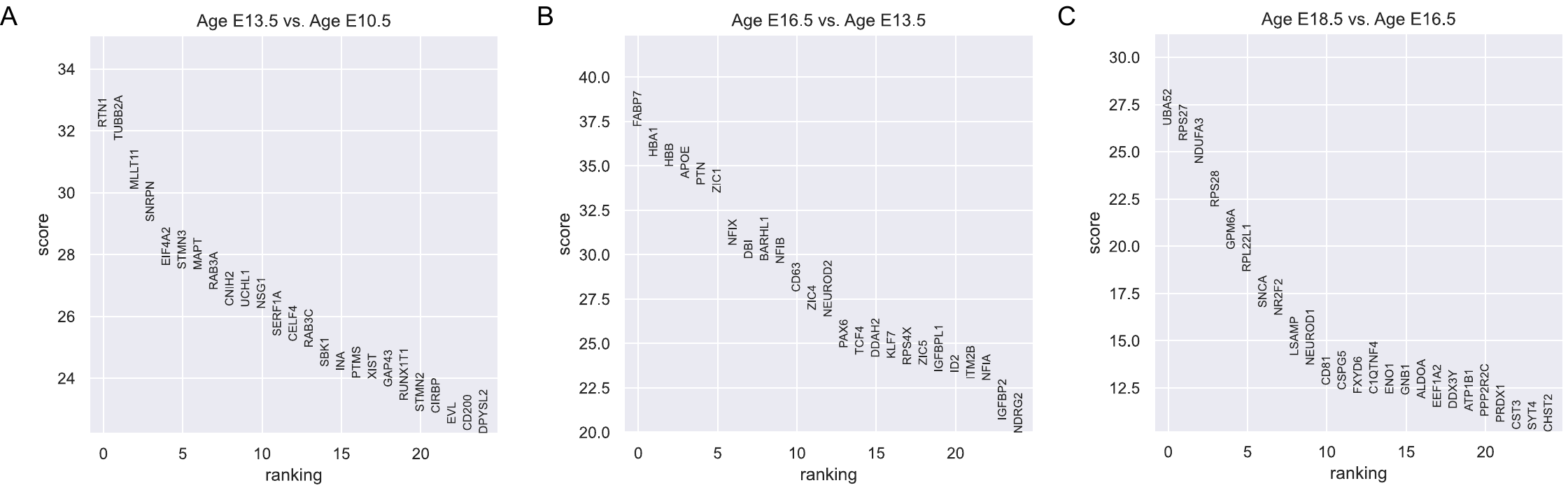

**Figure S16. Differentially expressed genes (DEGs) between adjacent age groups of Neurons in Pons dataset.** Differential expression analysis using Wilcoxon rank-sum is conducted for each pair of adjacent age groups. Genes are ranked according to the z-score underlying the p-value computation and the top 25 rankings are displayed. (A) DEGs between E13.5 and E10.5. (B) DEGs between E16.5 and E13.5. (C) DEGs between E18.5 and E16.5.

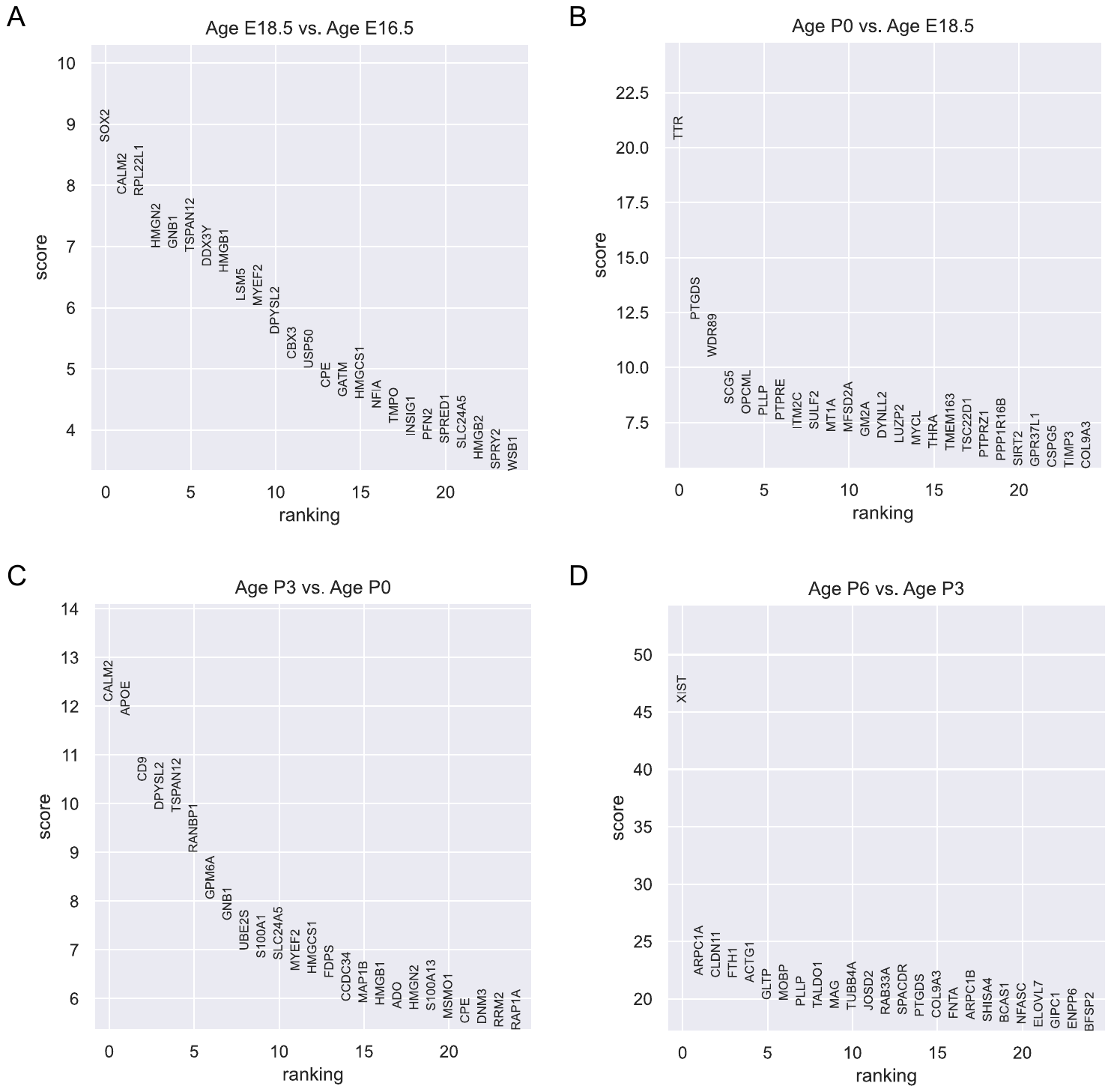

**Figure S17. Differentially expressed genes (DEGs) between adjacent age groups of Oligodendrocytes in Pons dataset.** Differential expression analysis using Wilcoxon rank-sum is conducted for each pair of adjacent age groups. Genes are ranked according to the z-score underlying the p-value computation and the top 25 rankings are displayed. (A) DEGs between E18.5 and E16.5. (B) DEGs between P0 and E18.5. (C) DEGs between P3 and P0. (D) DEGs between P6 and P3.

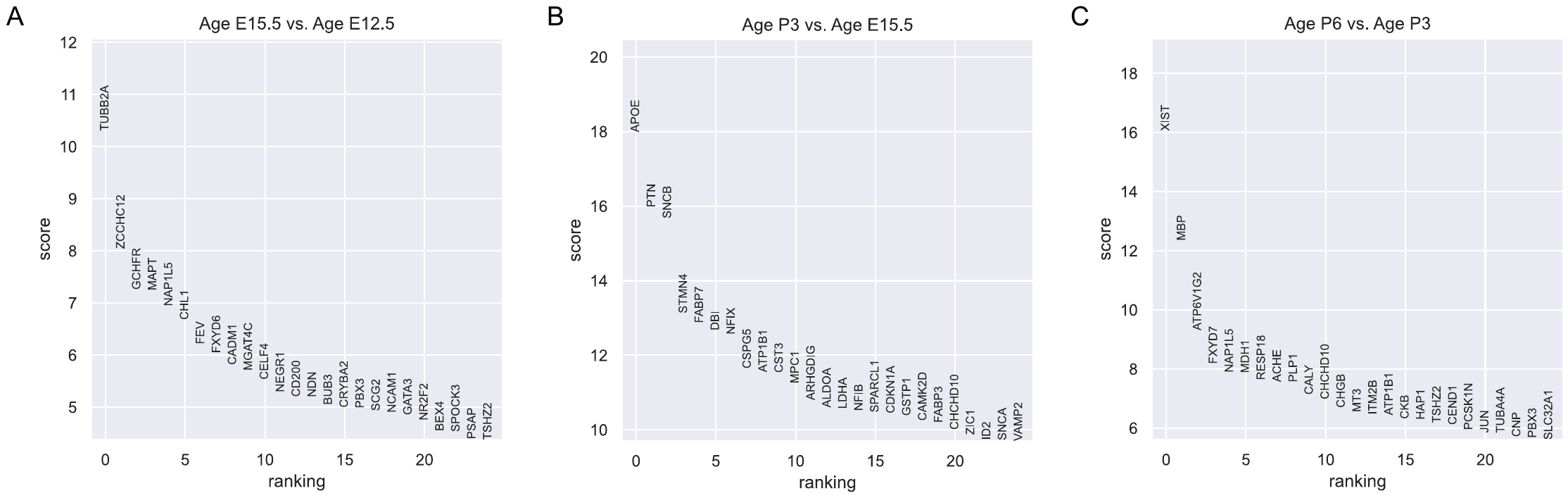

**Figure S18. Differentially expressed genes (DEGs) between adjacent age groups of Other neurons in Pons dataset.** Differential expression analysis using Wilcoxon rank-sum is conducted for each pair of adjacent age groups. Genes are ranked according to the z-score underlying the p-value computation and the top 25 rankings are displayed. (A) DEGs between E15.5 and E12.5. (B) DEGs between P3 and E15.5. (C) DEGs between P6 and P3.

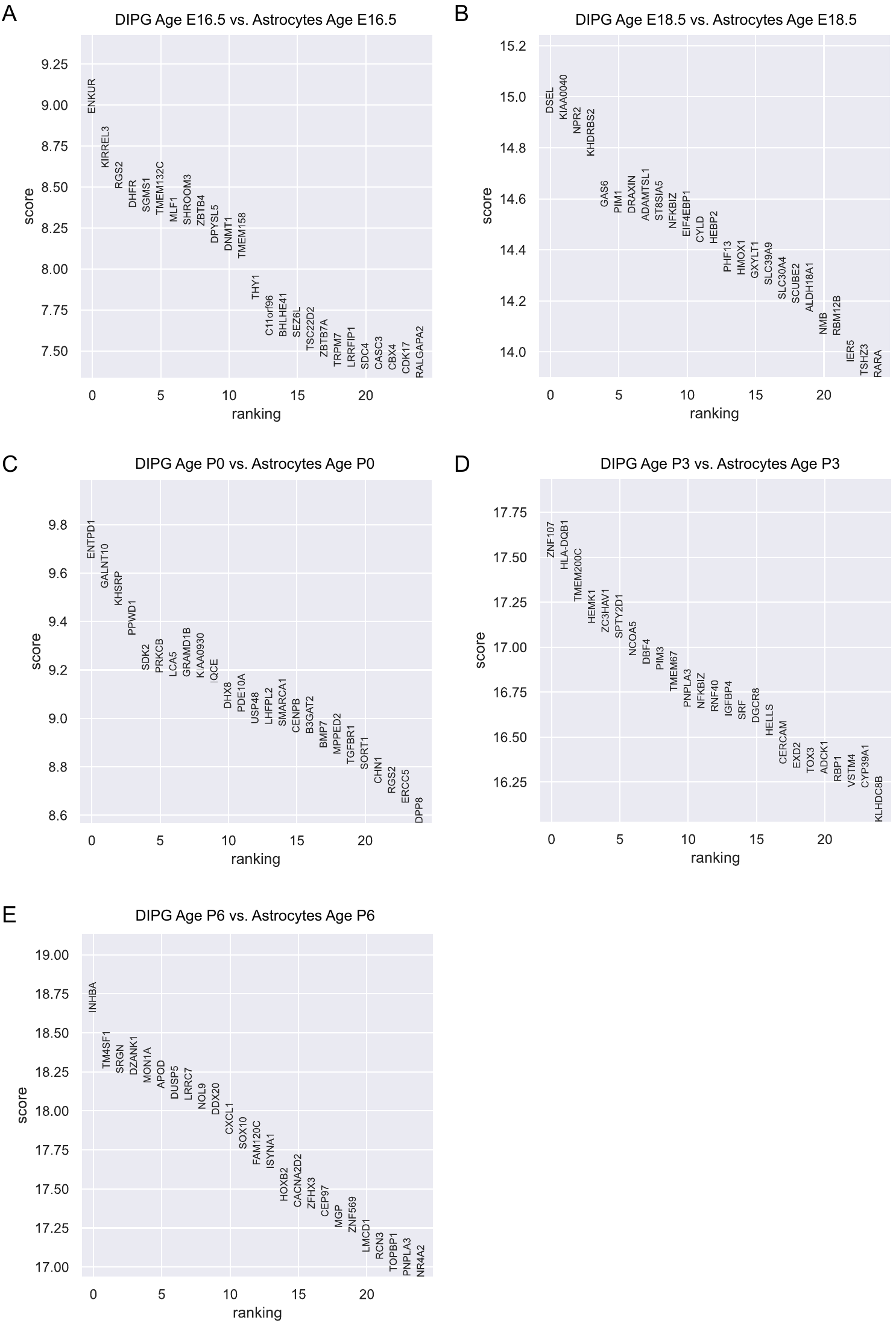

**Figure S19. Differentially expressed genes (DEGs) between DIPG and Astrocytes in Pons dataset.** Differential expression analysis using Wilcoxon rank-sum is conducted for each age group. Genes are ranked according to the z-score underlying the p-value computation and the top 25 rankings are displayed. (A) DEGs for E16.5. (B) DEGs for E18.5. (C) DEGs for P0. (D) DEGs for P3. (E) DEGs for P6.

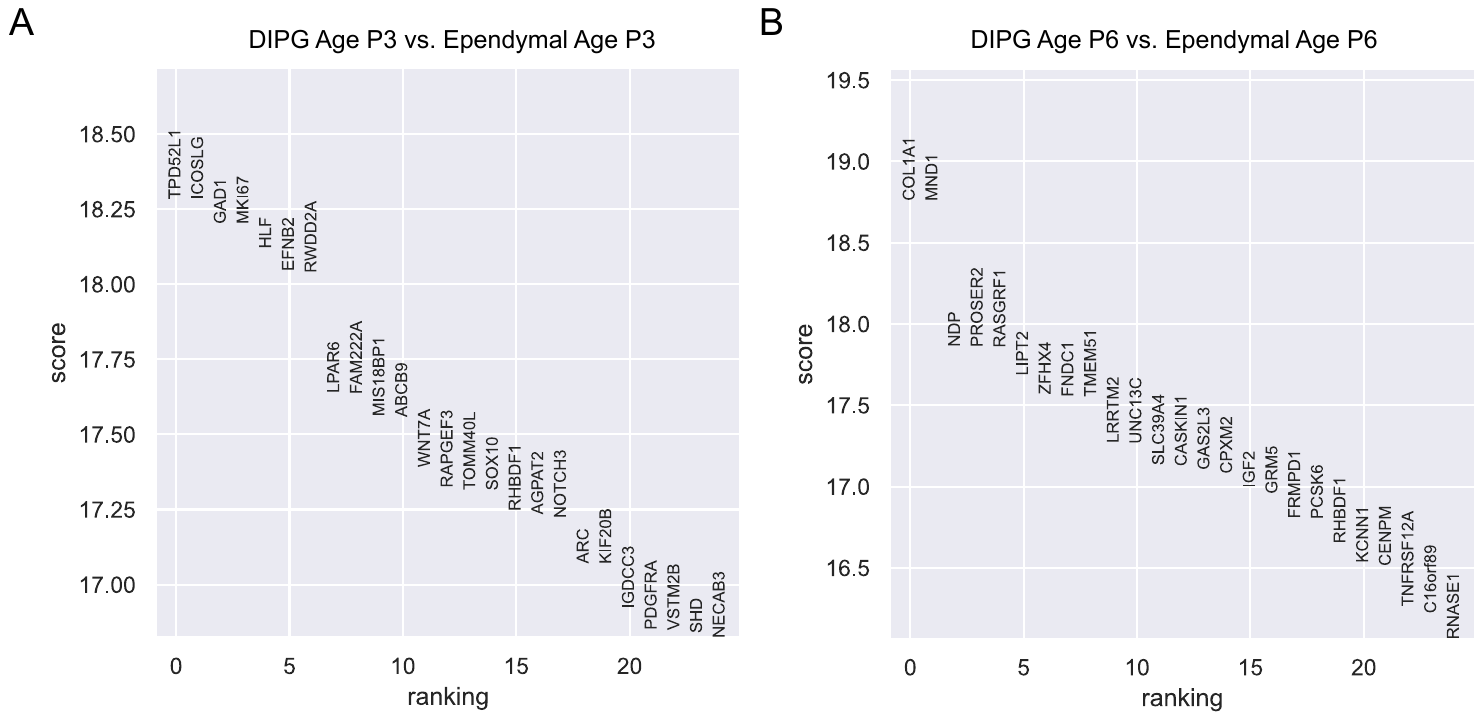

**Figure S20. Differentially expressed genes (DEGs) between DIPG and Ependymal in Pons dataset.** Differential expression analysis using Wilcoxon rank-sum is conducted for each age group. Genes are ranked according to the z-score underlying the p-value computation and the top 25 rankings are displayed. (A) DEGs for P3. (B) DEGs for P6.

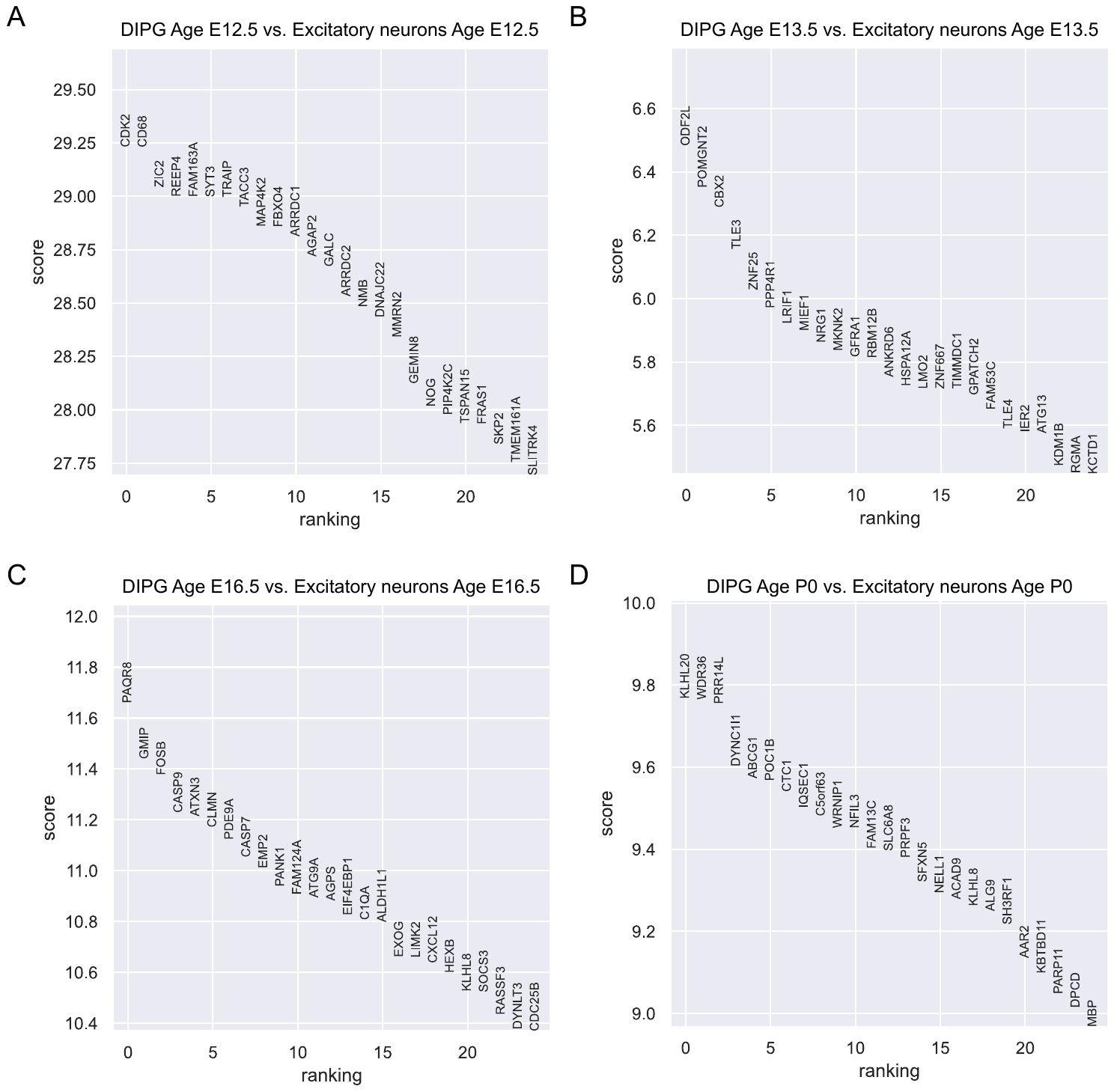

**Figure S21. Differentially expressed genes (DEGs) between DIPG and Excitatory neurons in Pons dataset.** Differential expression analysis using Wilcoxon rank-sum is conducted for each age group. Genes are ranked according to the z-score underlying the p-value computation and the top 25 rankings are displayed. (A) DEGs for E12.5. (B) DEGs for E13.5. (C) DEGs for E16.5. (D) DEGs for P0.

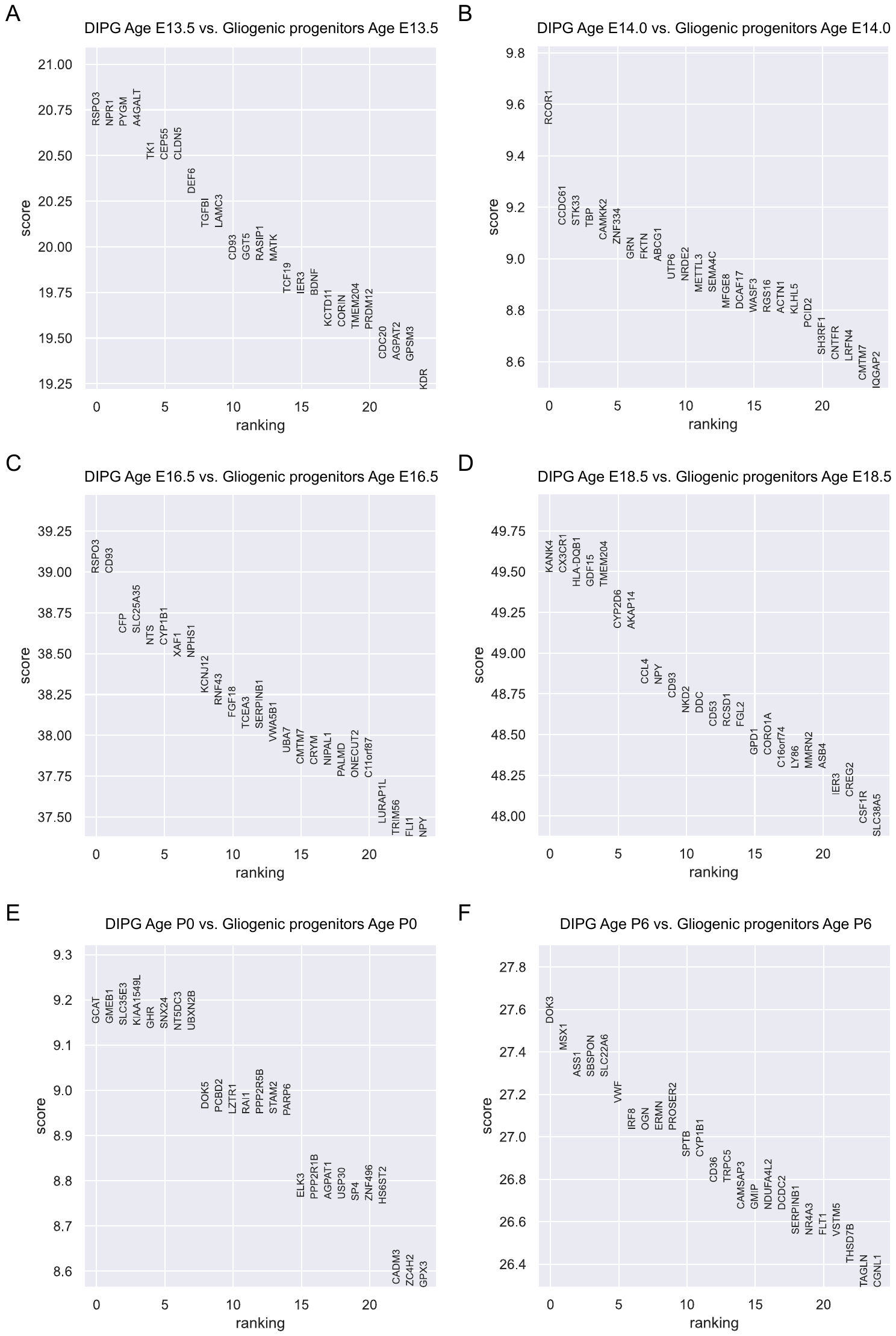

**Figure S22. Differentially expressed genes (DEGs) between DIPG and Gliogenic progenitors in Pons dataset.** Differential expression analysis using Wilcoxon rank-sum is conducted for each age group. Genes are ranked according to the z-score underlying the p-value computation and the top 25 rankings are displayed. (A) DEGs for E13.5. (B) DEGs for E14.0. (C) DEGs for E16.5. (D) DEGs for E18.5. (E) DEGs for P0. (F) DEGs for P6.

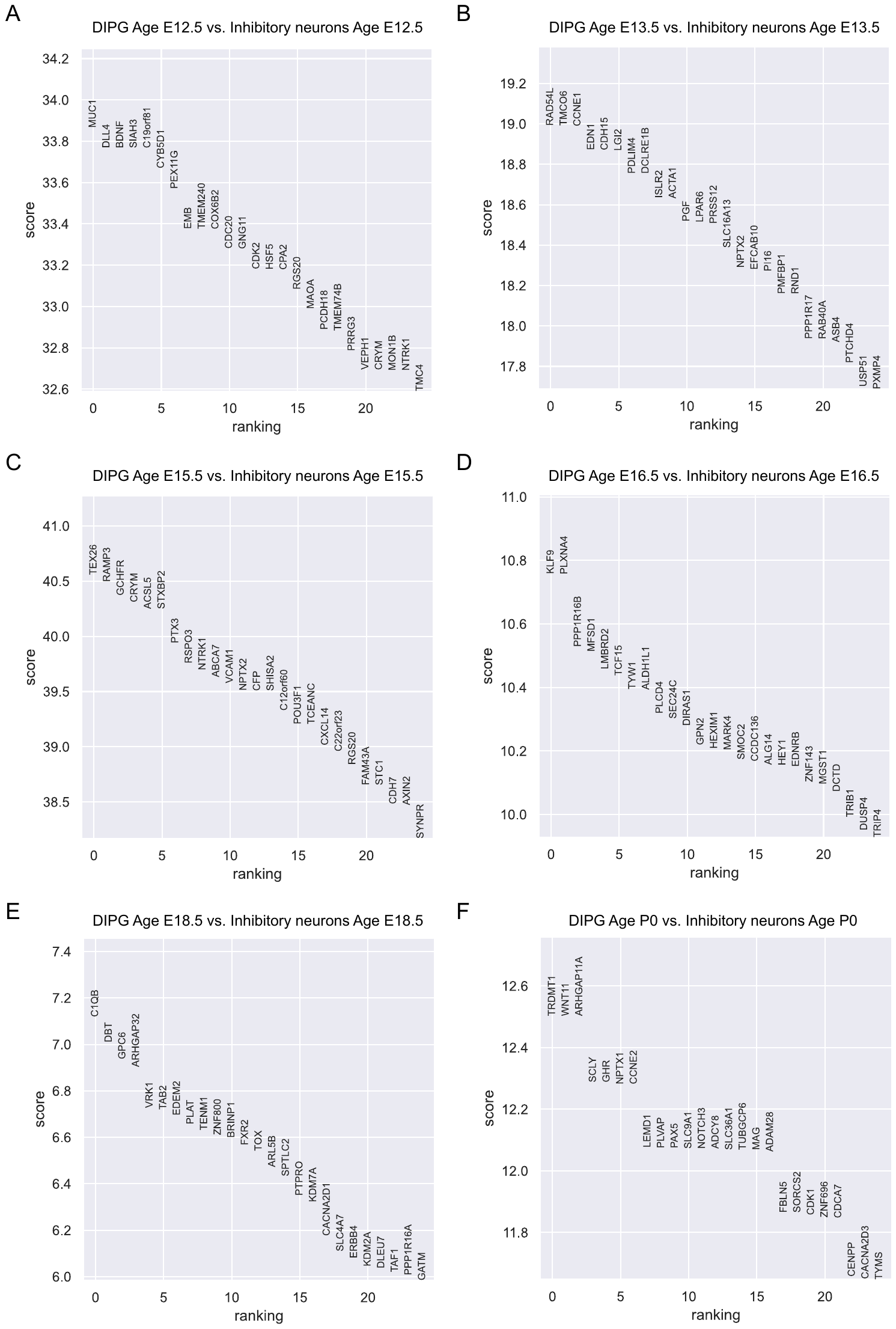

**Figure S23. Differentially expressed genes (DEGs) between DIPG and Inhibitory neurons in Pons dataset.** Differential expression analysis using Wilcoxon rank-sum is conducted for each age group. Genes are ranked according to the z-score underlying the p-value computation and the top 25 rankings are displayed. (A) DEGs for E12.5. (B) DEGs for E13.5. (C) DEGs for E15.5. (D) DEGs for E16.5. (E) DEGs for E18.5. (F) DEGs for P0.

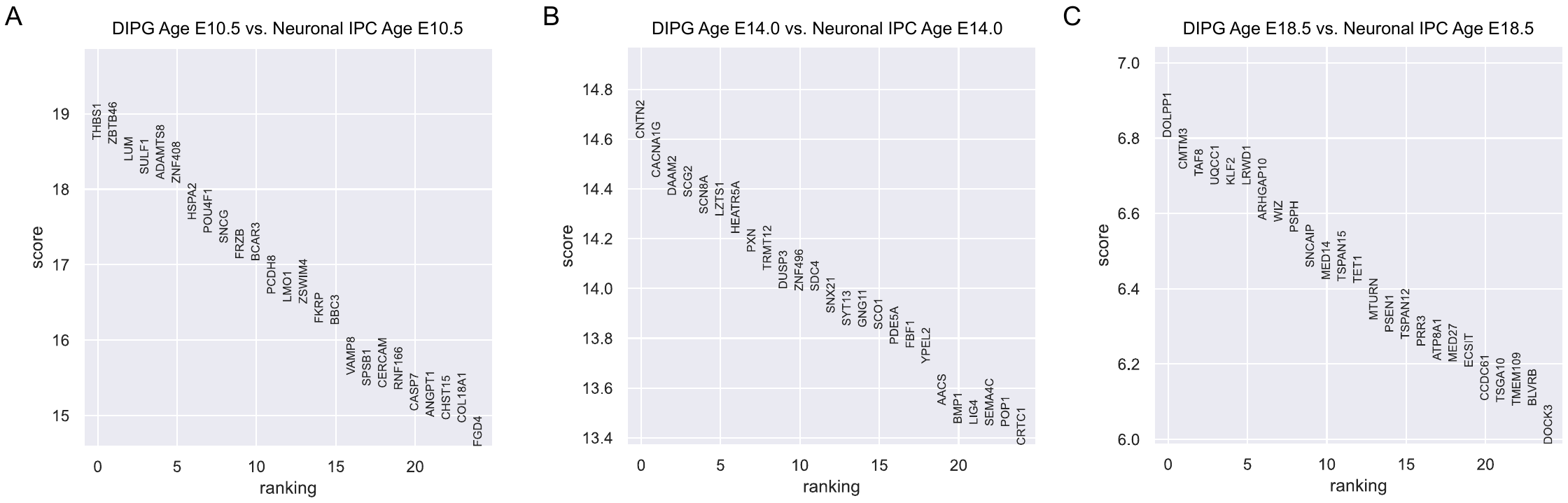

**Figure S24. Differentially expressed genes (DEGs) between DIPG and Neuronal IPC in Pons dataset.** Differential expression analysis using Wilcoxon rank-sum is conducted for each age group. Genes are ranked according to the z-score underlying the p-value computation and the top 25 rankings are displayed. (A) DEGs for E10.5. (B) DEGs for E14.0. (C) DEGs for E18.5.

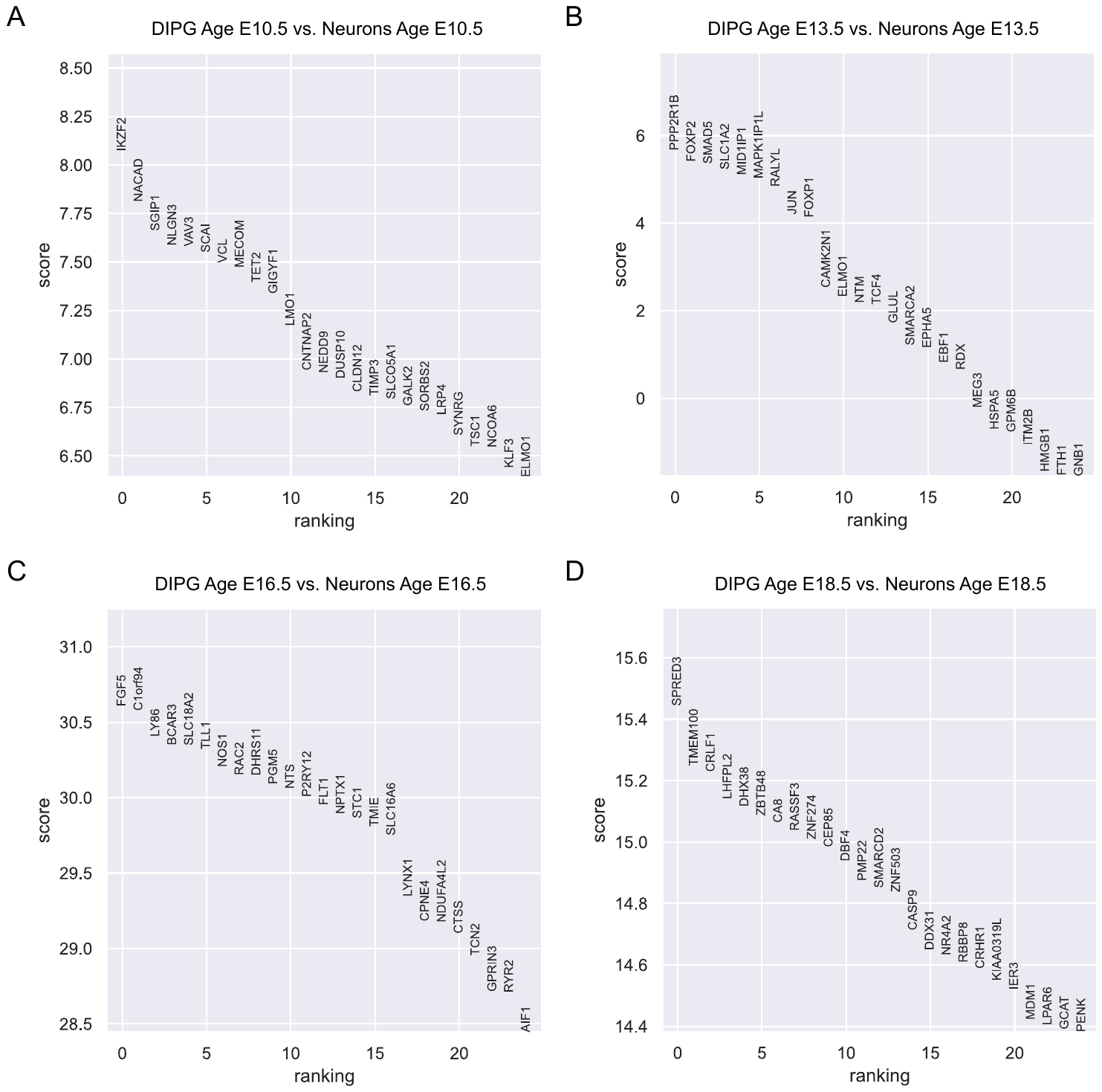

**Figure S25. Differentially expressed genes (DEGs) between DIPG and Neurons in Pons dataset.** Differential expression analysis using Wilcoxon rank-sum is conducted for each age group. Genes are ranked according to the z-score underlying the p-value computation and the top 25 rankings are displayed. (A) DEGs for E10.5. (B) DEGs for E13.5. (C) DEGs for E16.5. (D) DEGs for E18.5.

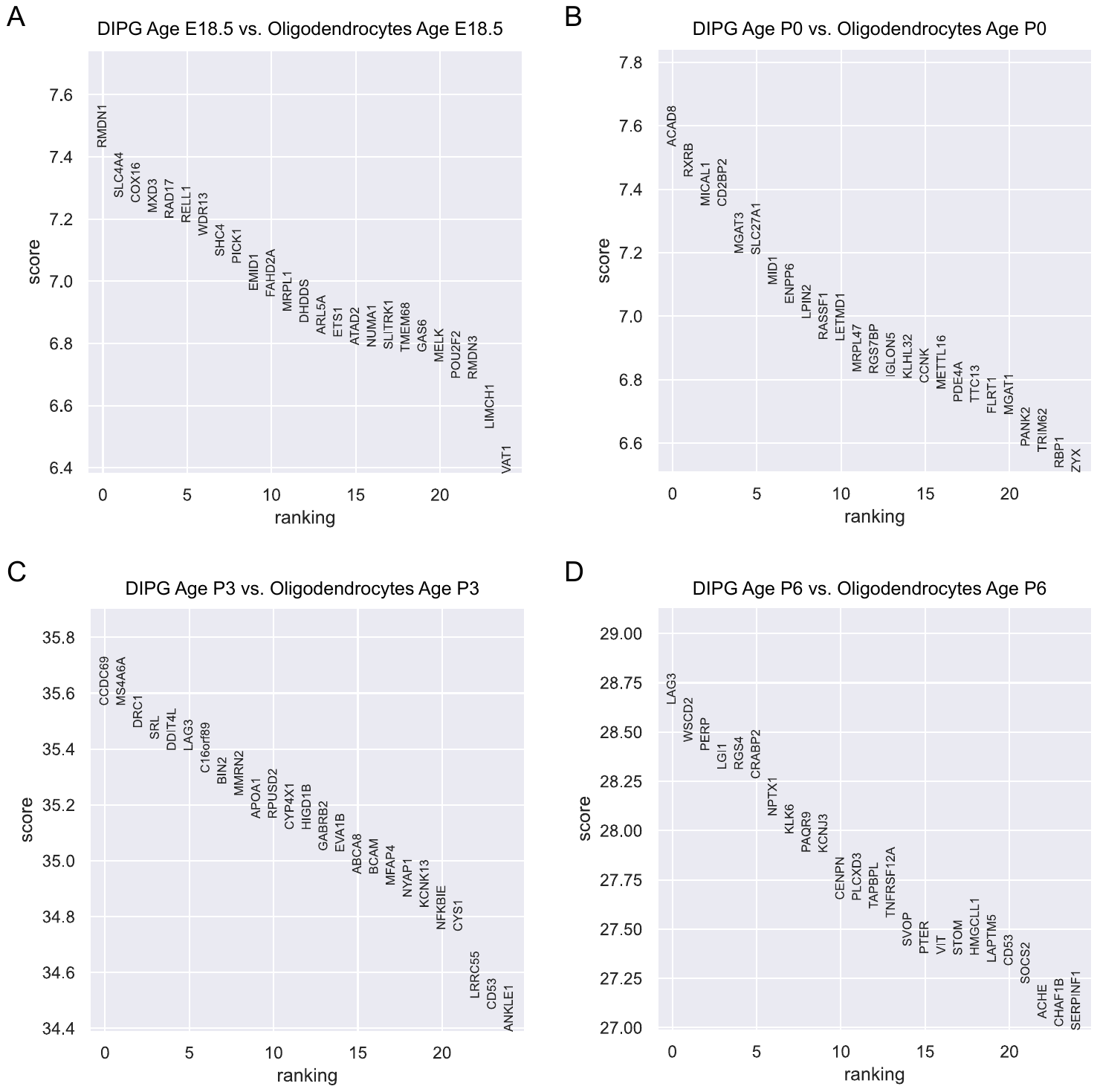

**Figure S26. Differentially expressed genes (DEGs) between DIPG and Oligodendrocytes in Pons dataset.** Differential expression analysis using Wilcoxon rank-sum is conducted for each age group. Genes are ranked according to the z-score underlying the p-value computation and the top 25 rankings are displayed. (A) DEGs for E18.5. (B) DEGs for P0. (C) DEGs for P3. (D) DEGs for P6.

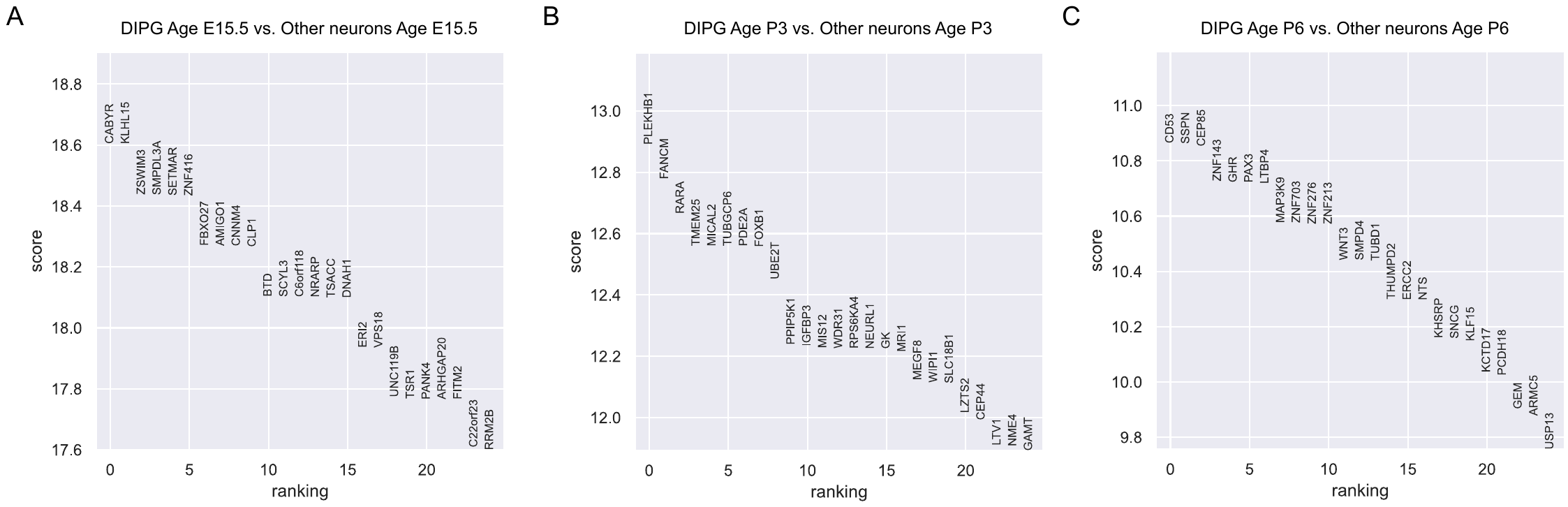

**Figure S27. Differentially expressed genes (DEGs) between DIPG and Other neurons in Pons dataset.** Differential expression analysis using Wilcoxon rank-sum is conducted for each age group. Genes are ranked according to the z-score underlying the p-value computation and the top 25 rankings are displayed. (A) DEGs for E15.5. (B) DEGs for P3. (C) DEGs for P6.

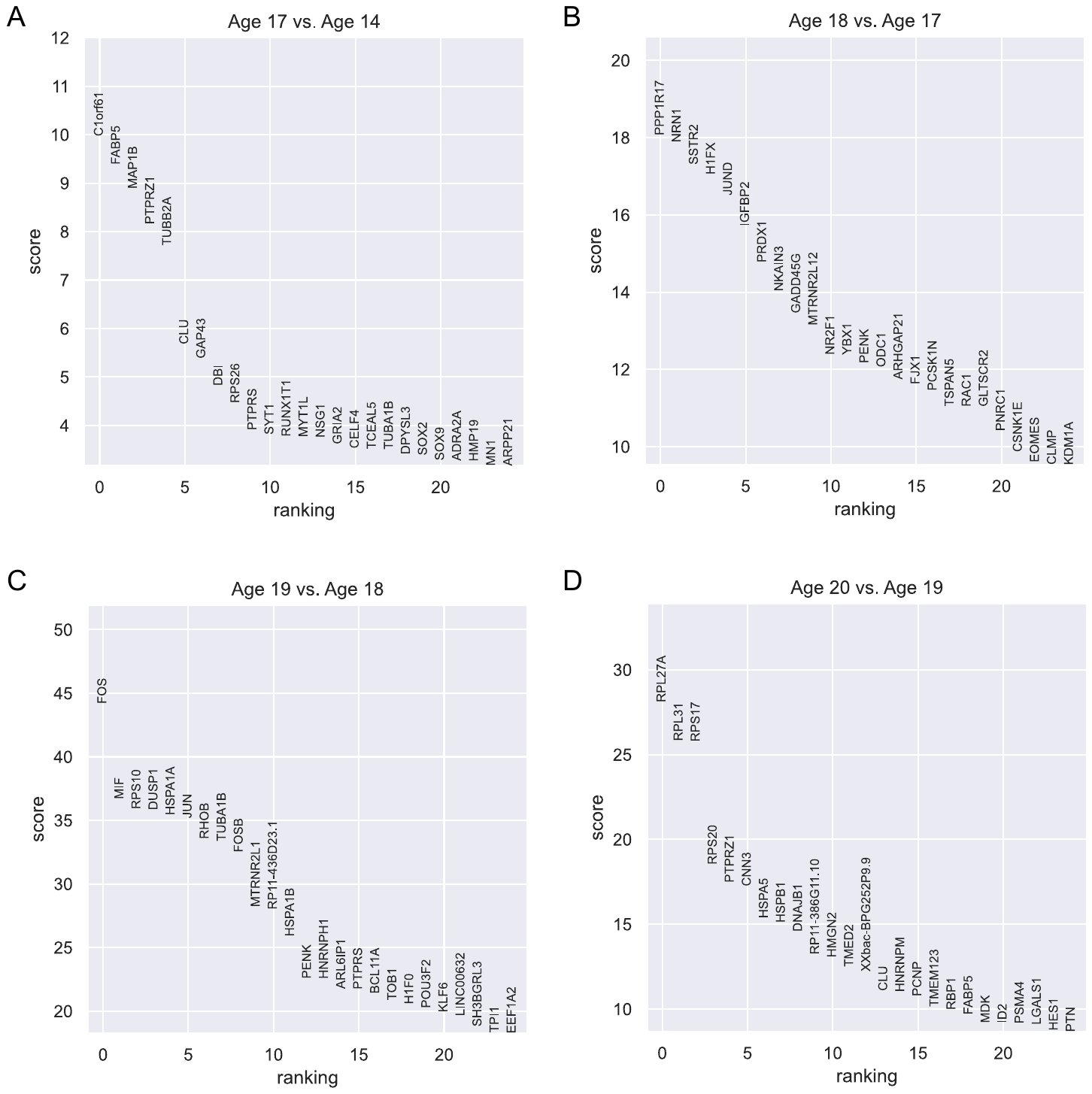

**Figure S28. Differentially expressed genes (DEGs) between adjacent age groups of IPC in Bhaduri’s dataset.** Differential expression analysis using Wilcoxon rank-sum is conducted for each pair of adjacent age groups. Genes are ranked according to the z-score underlying the p-value computation and the top 25 rankings are displayed. (A) DEGs between Age 17 and Age 14. (B) DEGs between Age 18 and Age 17. (C) DEGs between Age 19 and Age 18. (D) DEGs between Age 20 and Age 19.

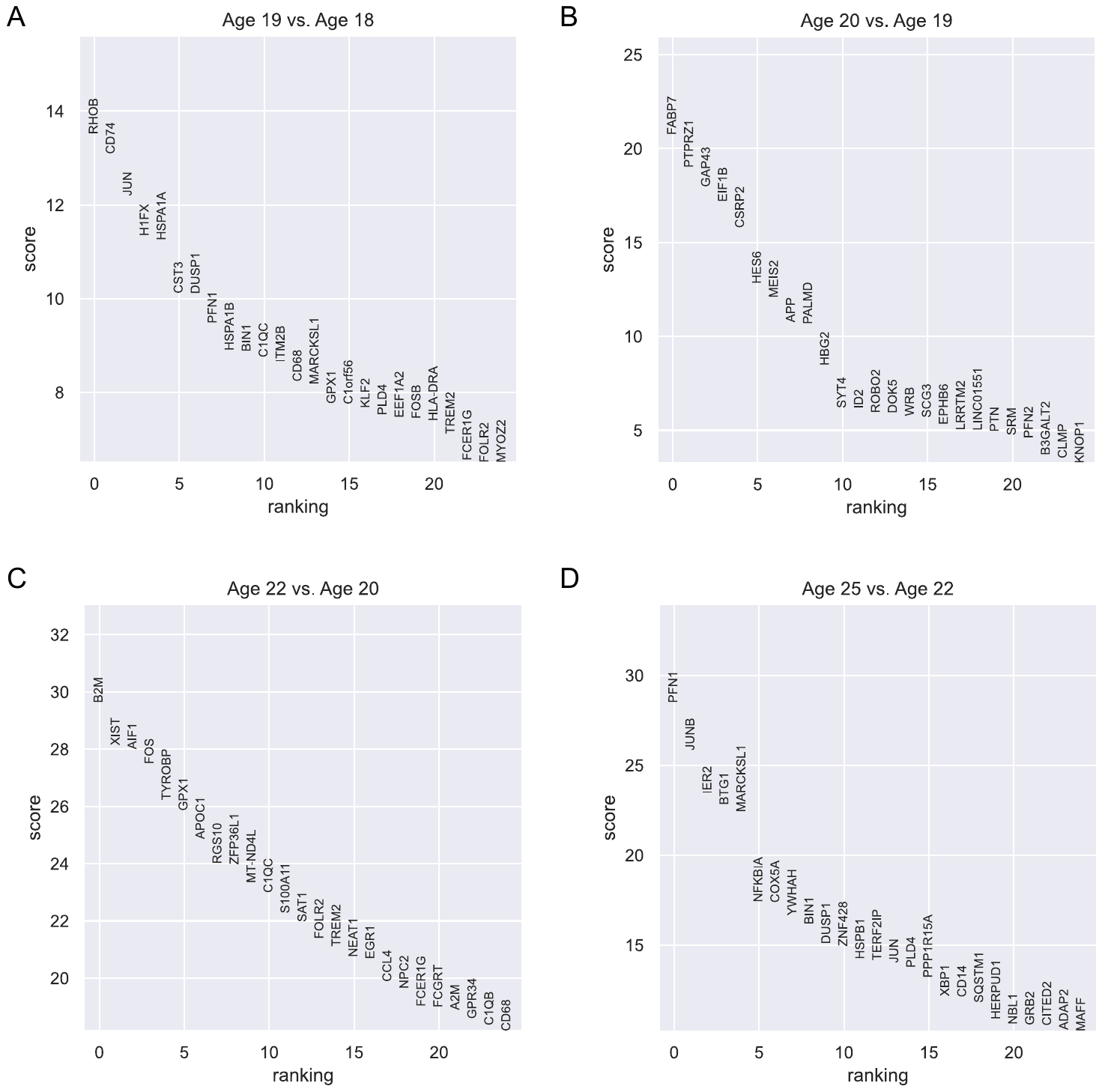

**Figure S29. Differentially expressed genes (DEGs) between adjacent age groups of Microglia in Bhaduri’s dataset.** Differential expression analysis using Wilcoxon rank-sum is conducted for each pair of adjacent age groups. Genes are ranked according to the z-score underlying the p-value computation and the top 25 rankings are displayed. (A) DEGs between Age 19 and Age 18. (B) DEGs between Age 20 and Age 19. (C) DEGs between Age 22 and Age 20. (D) DEGs between Age 25 and Age 22.

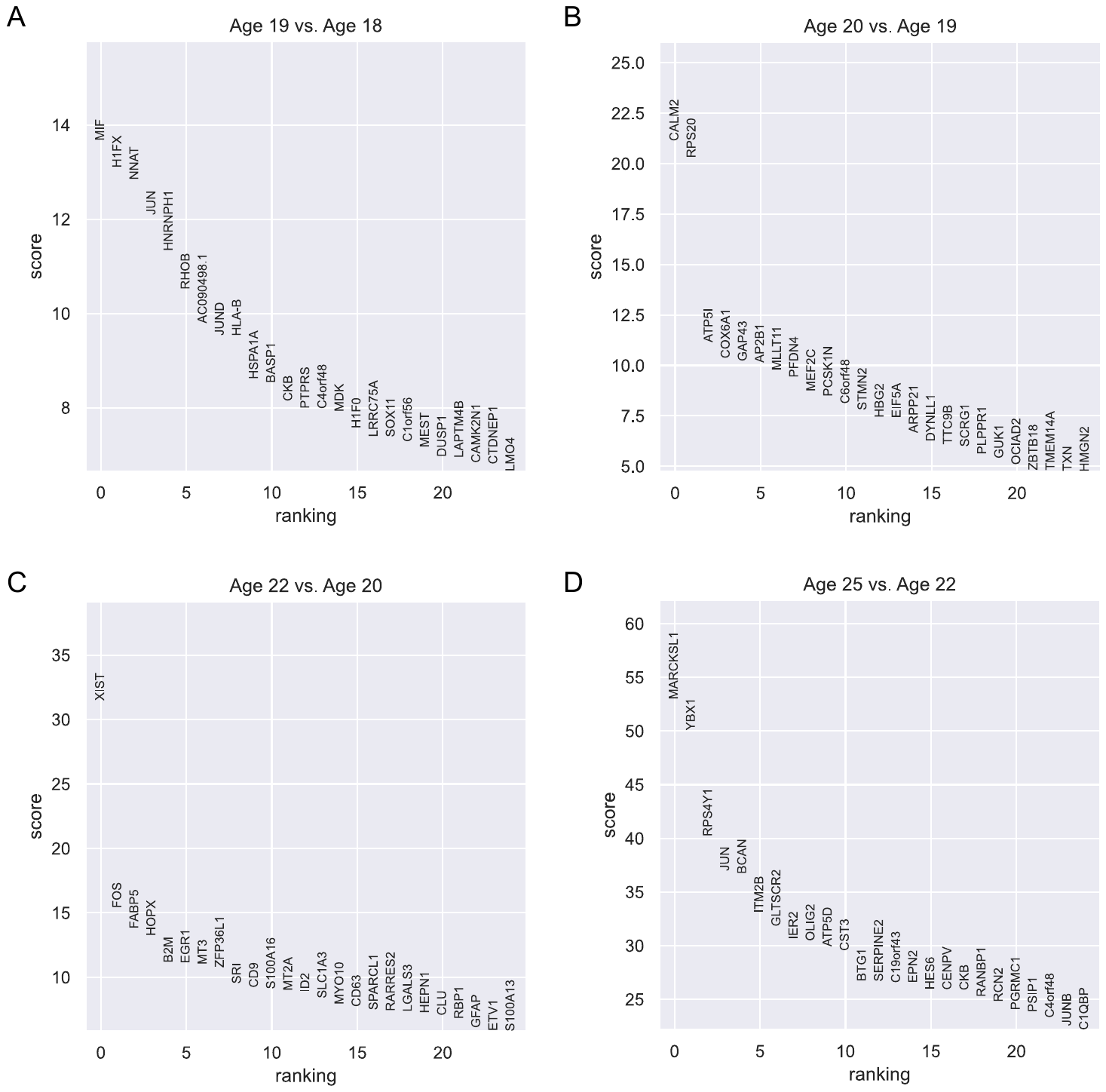

**Figure S30. Differentially expressed genes (DEGs) between adjacent age groups of OPC in Bhaduri’s dataset.** Differential expression analysis using Wilcoxon rank-sum is conducted for each pair of adjacent age groups. Genes are ranked according to the z-score underlying the p-value computation and the top 25 rankings are displayed. (A) DEGs between Age 19 and Age 18. (B) DEGs between Age 20 and Age 19. (C) DEGs between Age 22 and Age 20. (D) DEGs between Age 25 and Age 22.

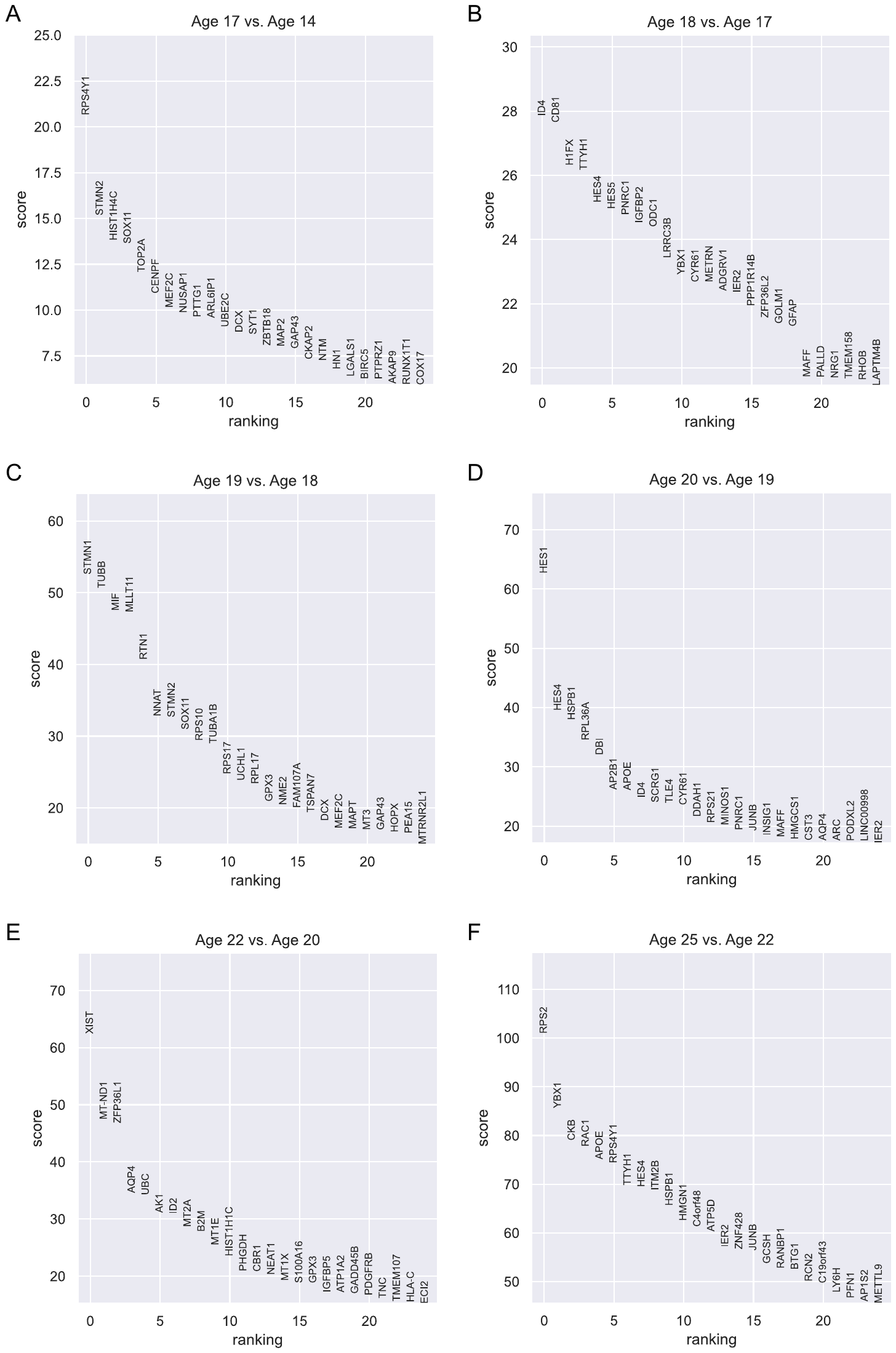

**Figure S31. Differentially expressed genes (DEGs) between adjacent age groups of RG in Bhaduri’s dataset.** Differential expression analysis using Wilcoxon rank-sum is conducted for each pair of adjacent age groups. Genes are ranked according to the z-score underlying the p-value computation and the top 25 rankings are displayed. (A) DEGs between Age 17 and Age 14. (B) DEGs between Age 18 and Age 17. (C) DEGs between Age 19 and Age 18. (D) DEGs between Age 20 and Age 19. (E) DEGs between Age 22 and Age 20. (F) DEGs between Age 25 and Age 22.

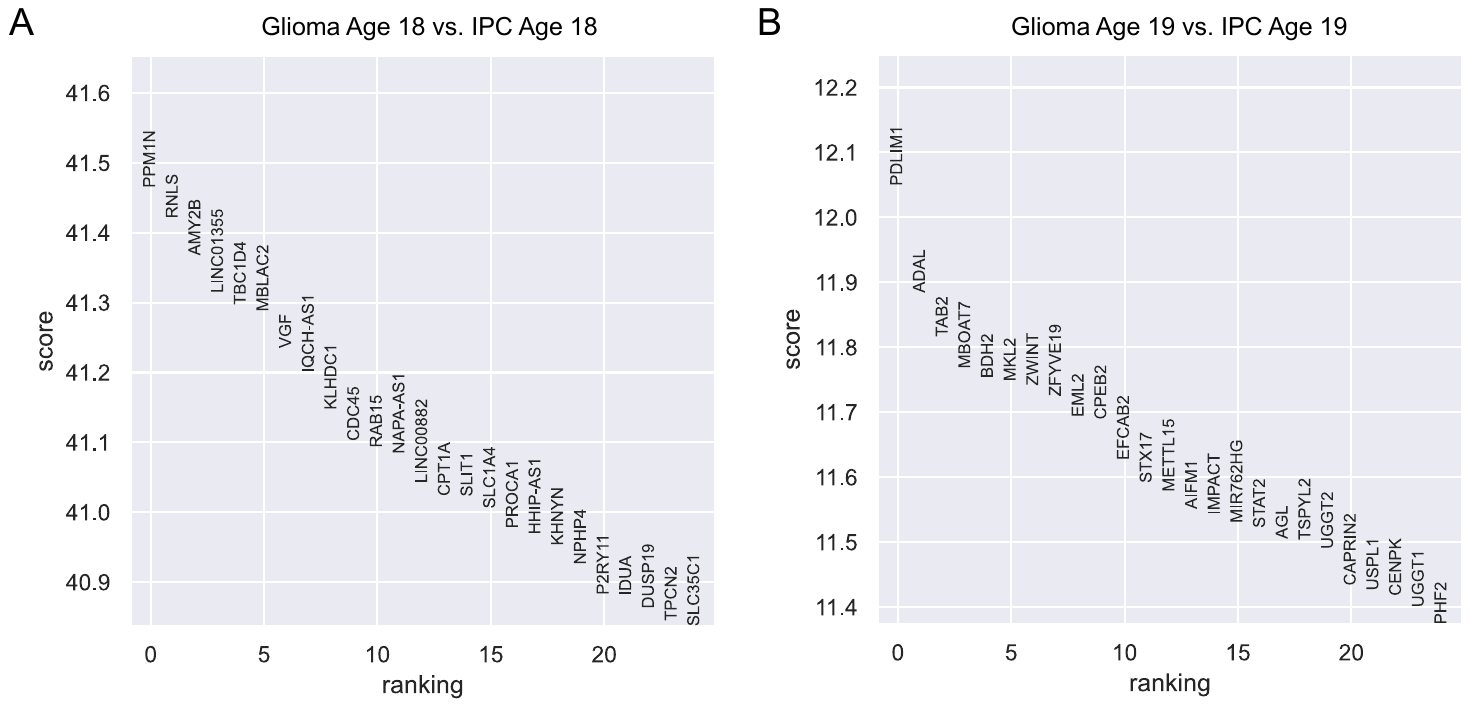

**Figure S32. Differentially expressed genes (DEGs) between glioma and IPC in Bhaduri’s dataset.** Differential expression analysis using Wilcoxon rank-sum is conducted for each age group. Genes are ranked according to the z-score underlying the p-value computation and the top 25 rankings are displayed. (A) DEGs for Age 18. (B) DEGs for Age 19.

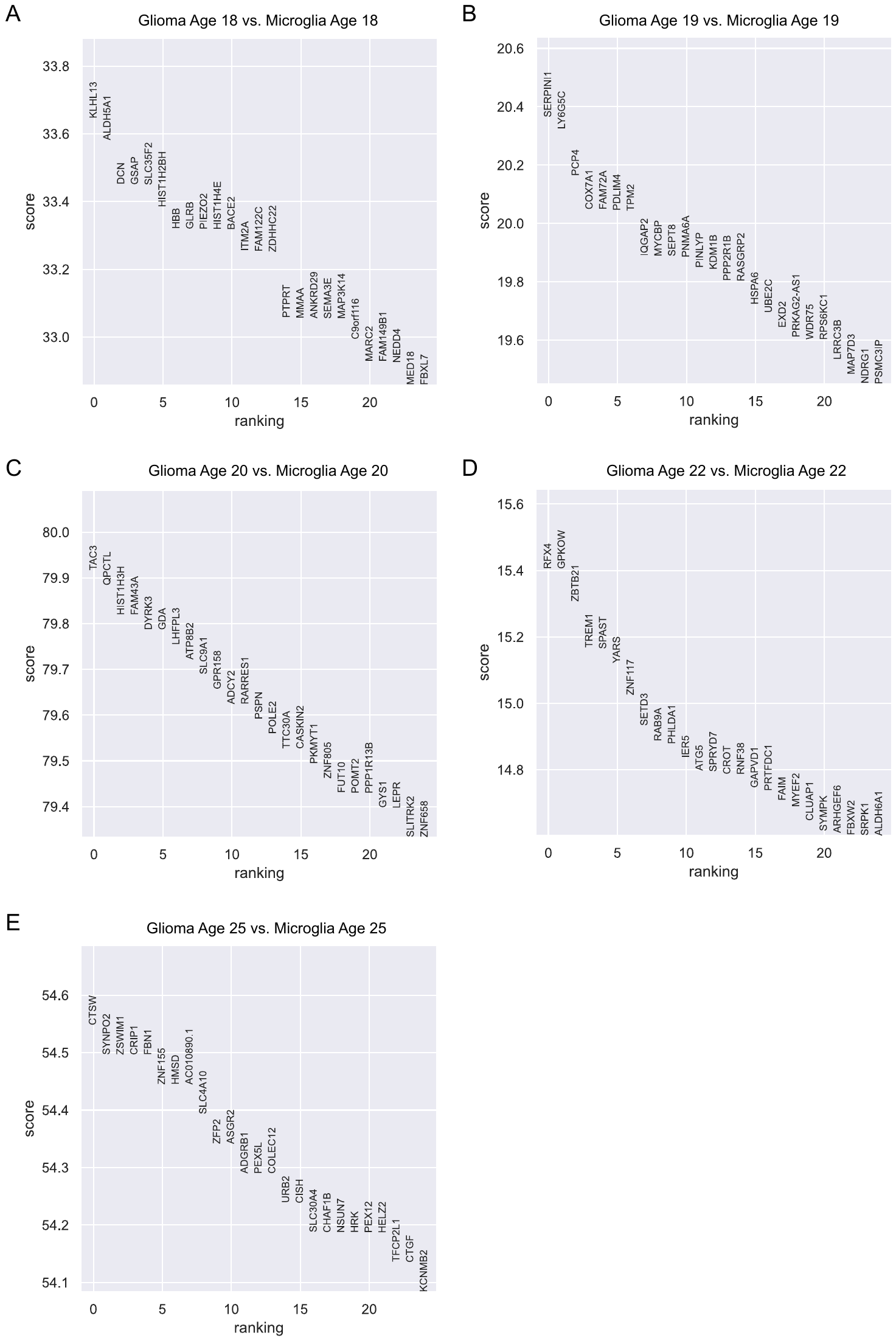

**Figure S33. Differentially expressed genes (DEGs) between glioma and Microglia in Bhaduri’s dataset.** Differential expression analysis using Wilcoxon rank-sum is conducted for each age group. Genes are ranked according to the z-score underlying the p-value computation and the top 25 rankings are displayed. (A) DEGs for Age 18. (B) DEGs for Age 19. (C) DEGs for Age 20. (D) DEGs for Age 22. (E) DEGs for Age 25.

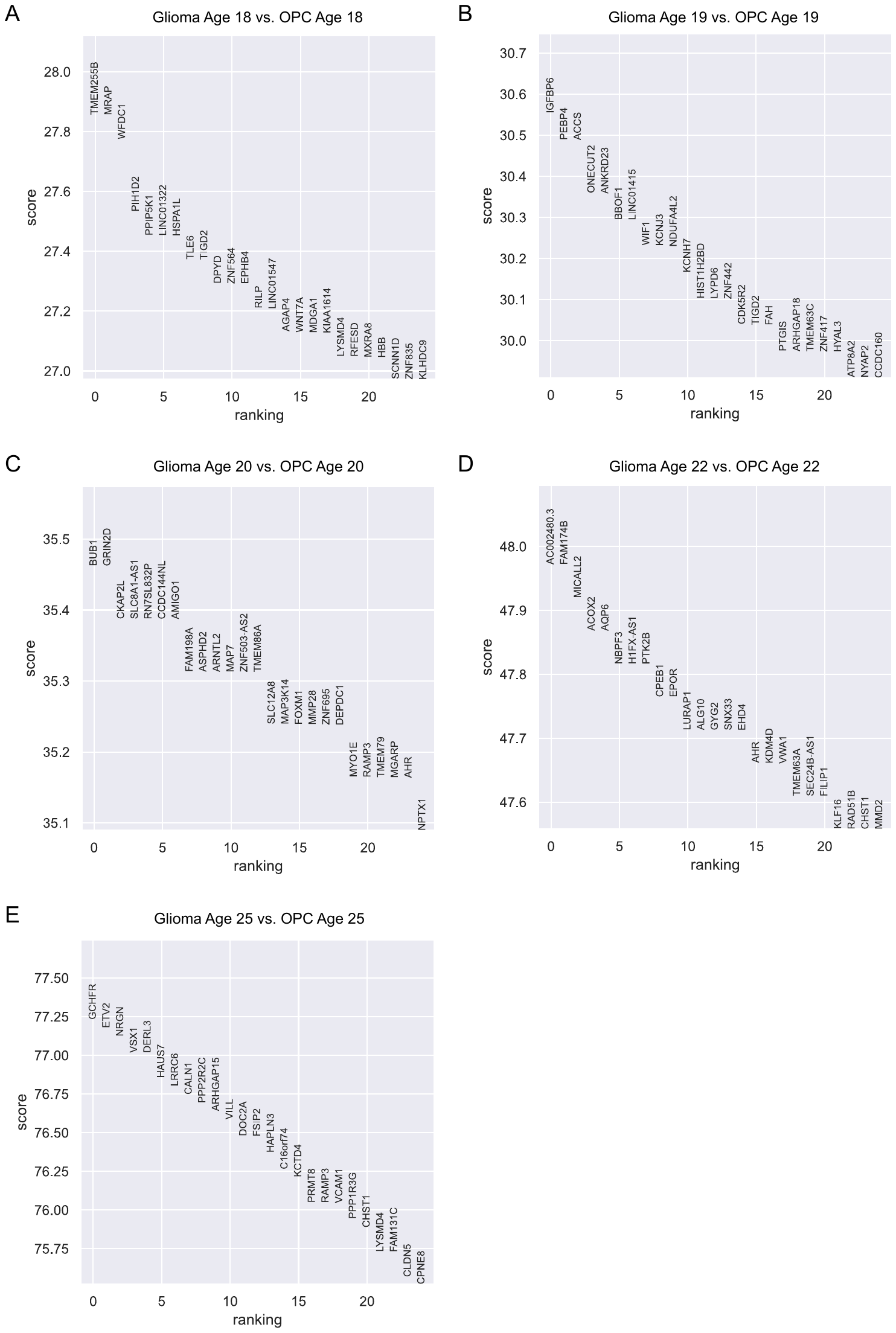

**Figure S34. Differentially expressed genes (DEGs) between glioma and OPC in Bhaduri’s dataset.** Differential expression analysis using Wilcoxon rank-sum is conducted for each age group. Genes are ranked according to the z-score underlying the p-value computation and the top 25 rankings are displayed. (A) DEGs for Age 18. (B) DEGs for Age 19. (C) DEGs for Age 20. (D) DEGs for Age 22. (E) DEGs for Age 25.

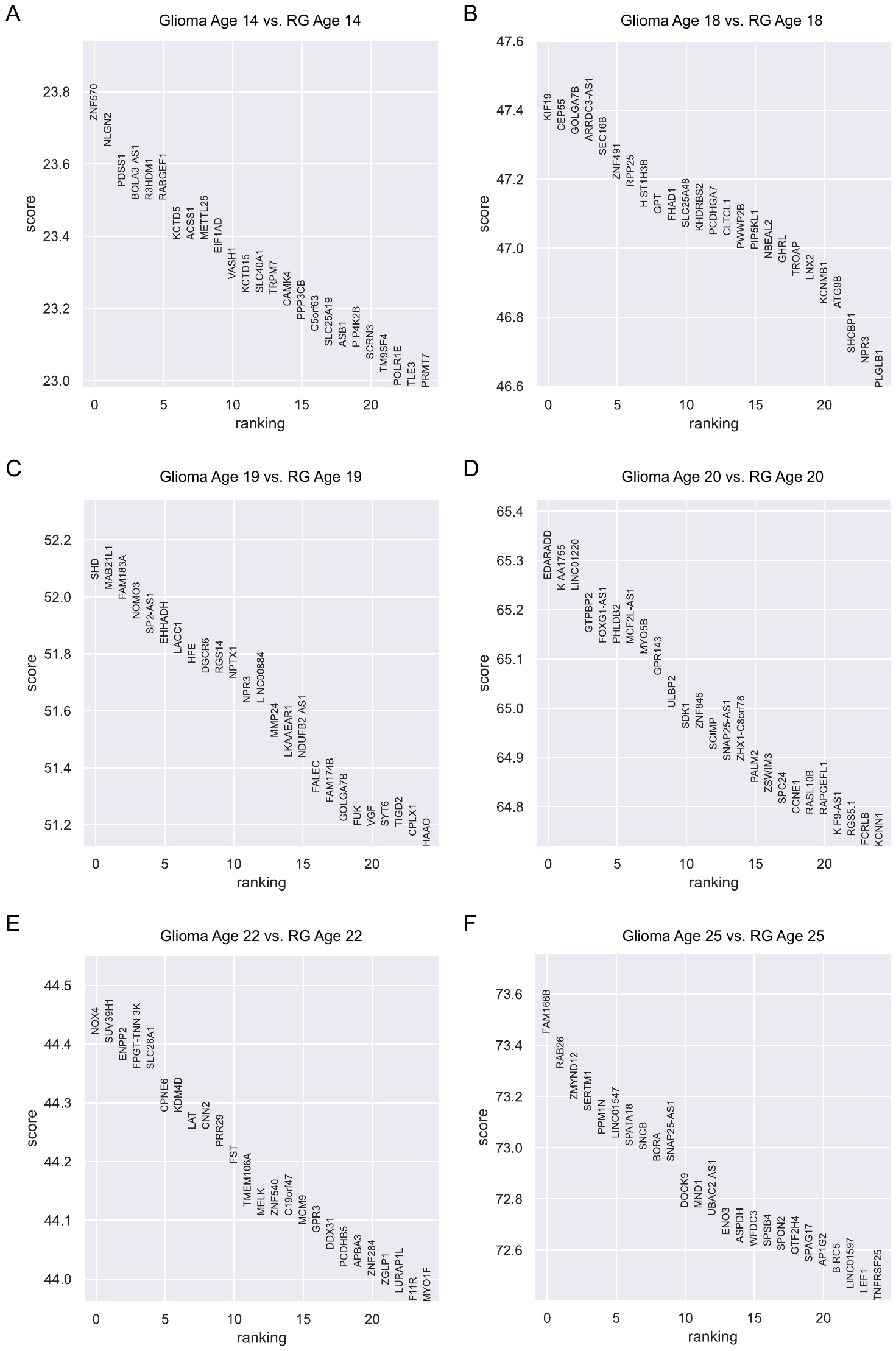

**Figure S35. Differentially expressed genes (DEGs) between glioma and RG in Bhaduri’s dataset.** Differential expression analysis using Wilcoxon rank-sum is conducted for each age group. Genes are ranked according to the z-score underlying the p-value computation and the top 25 rankings are displayed. (A) DEGs for Age 14. (B) DEGs for Age 18. (C) DEGs for Age 19. (D) DEGs for Age 20. (E) DEGs for Age 22. (F) DEGs for Age 25.

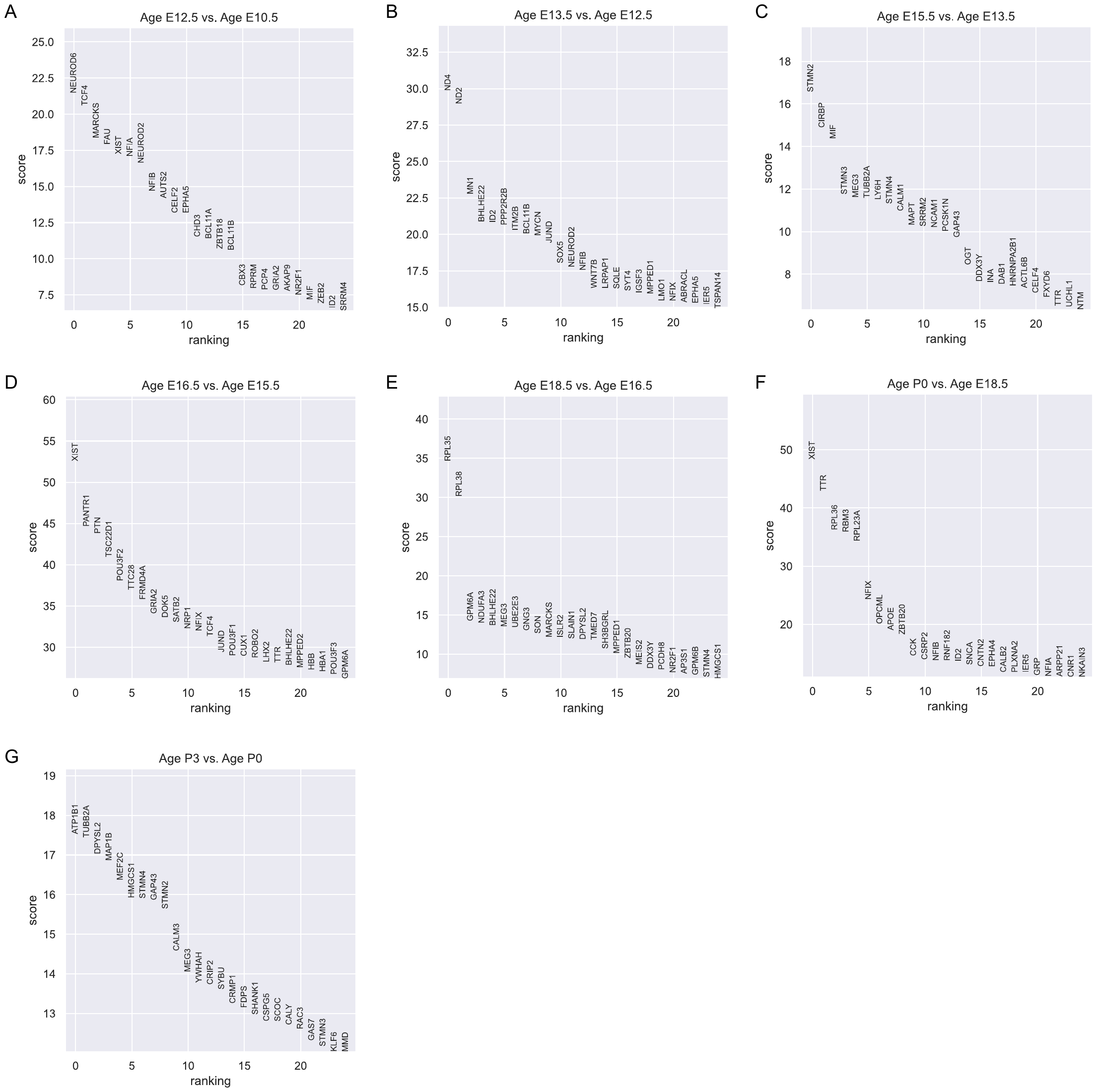

**Figure S36. Differentially expressed genes (DEGs) between adjacent age groups of Excitatory neurons in Forebrain dataset.** Differential expression analysis using Wilcoxon rank-sum is conducted for each pair of adjacent age groups. Genes are ranked according to the z-score underlying the p-value computation and the top 25 rankings are displayed. (A) DEGs between E12.5 and E10.5. (B) DEGs between E13.5 and E12.5. (C) DEGs between E15.5 and E13.5. (D) DEGs between E16.5 and E15.5. (E) DEGs between E18.5 and E16.5. (F) DEGs between P0 and E18.5. (G) DEGs between P3 and P0.

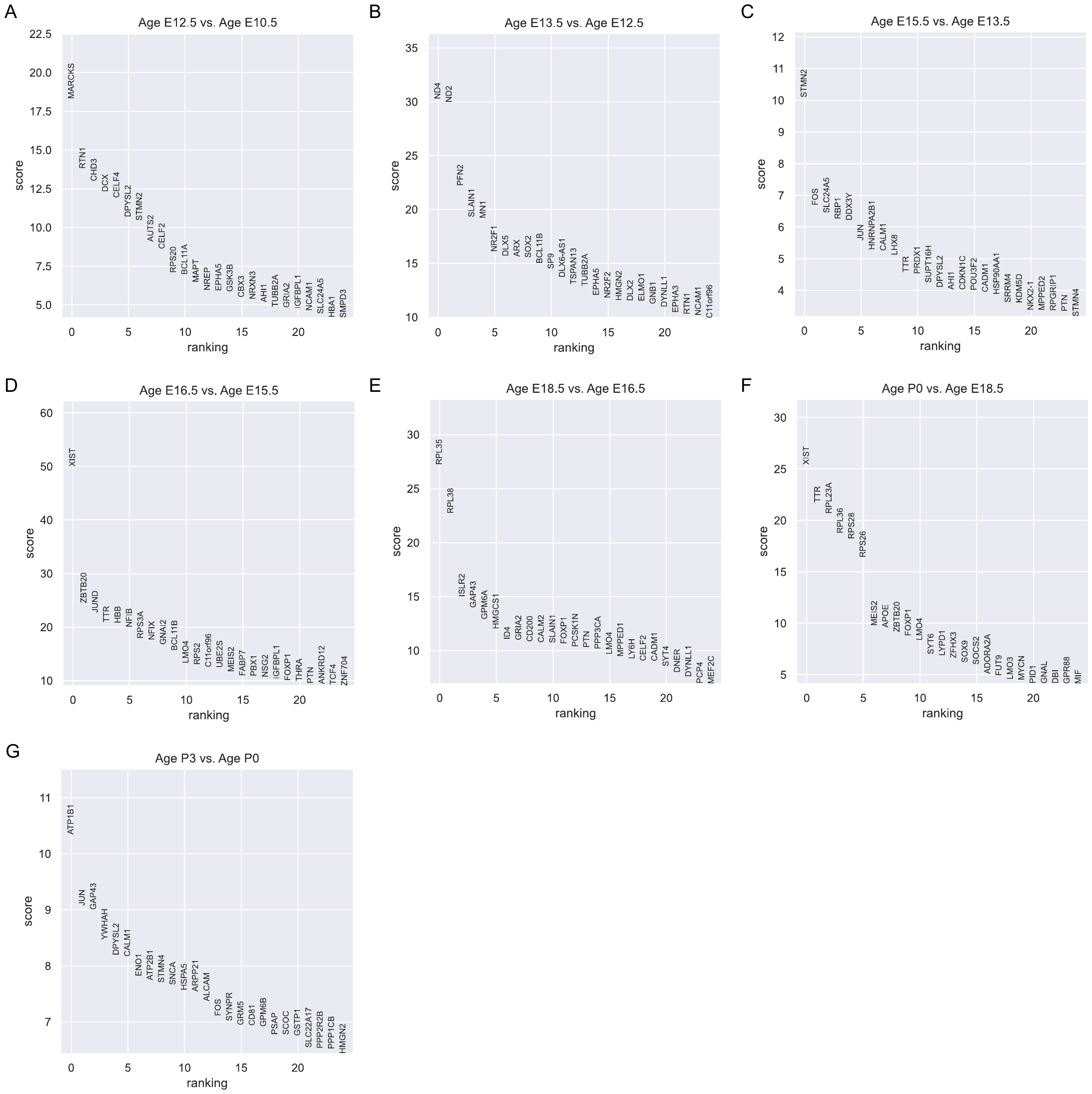

**Figure S37. Differentially expressed genes (DEGs) between adjacent age groups of Inhibitory neurons in Forebrain dataset.** Differential expression analysis using Wilcoxon rank-sum is conducted for each pair of adjacent age groups. Genes are ranked according to the z-score underlying the p-value computation and the top 25 rankings are displayed. (A) DEGs between E12.5 and E10.5. (B) DEGs between E13.5 and E12.5. (C) DEGs between E15.5 and E13.5. (D) DEGs between E16.5 and E15.5. (E) DEGs between E18.5 and E16.5. (F) DEGs between P0 and E18.5. (G) DEGs between P3 and P0.

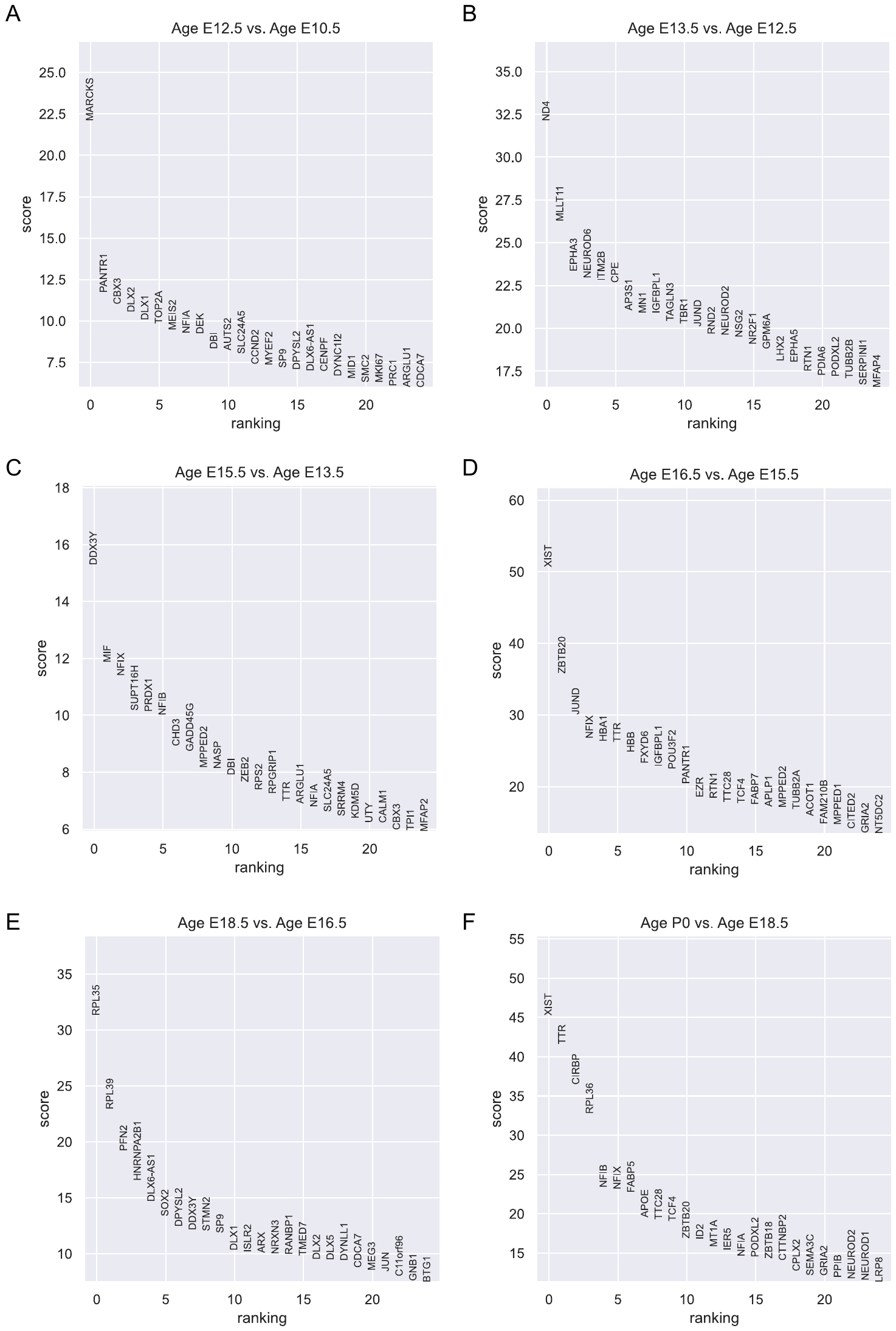

**Figure S38. Differentially expressed genes (DEGs) between adjacent age groups of Neuronal IPC in Forebrain dataset.** Differential expression analysis using Wilcoxon rank-sum is conducted for each pair of adjacent age groups. Genes are ranked according to the z-score underlying the p-value computation and the top 25 rankings are displayed. (A) DEGs between E12.5 and E10.5. (B) DEGs between E13.5 and E12.5. (C) DEGs between E15.5 and E13.5. (D) DEGs between E16.5 and E15.5. (E) DEGs between E18.5 and E16.5. (F) DEGs between P0 and E18.5.

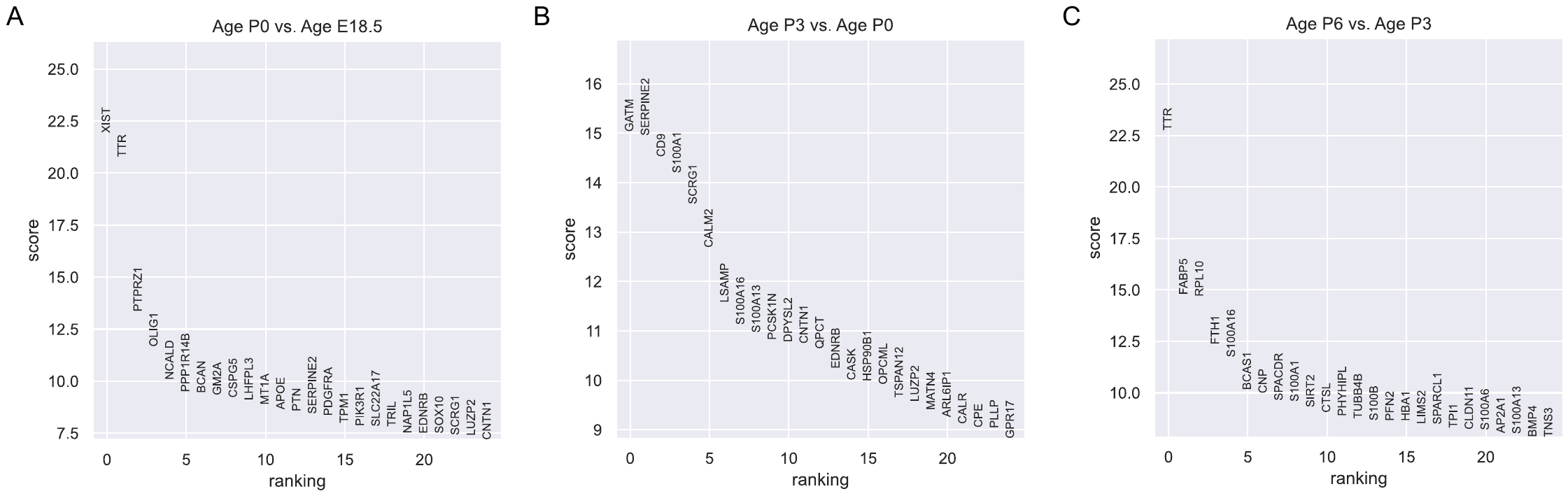

**Figure S39. Differentially expressed genes (DEGs) between adjacent age groups of Oligodendrocytes in Forebrain dataset**. Differential expression analysis using Wilcoxon rank-sum is conducted for each pair of adjacent age groups. Genes are ranked according to the z-score underlying the p-value computation and the top 25 rankings are displayed. (A) DEGs between P0 and E18.5. (B) DEGs between P3 and P0. (C) DEGs between P6 and P3.

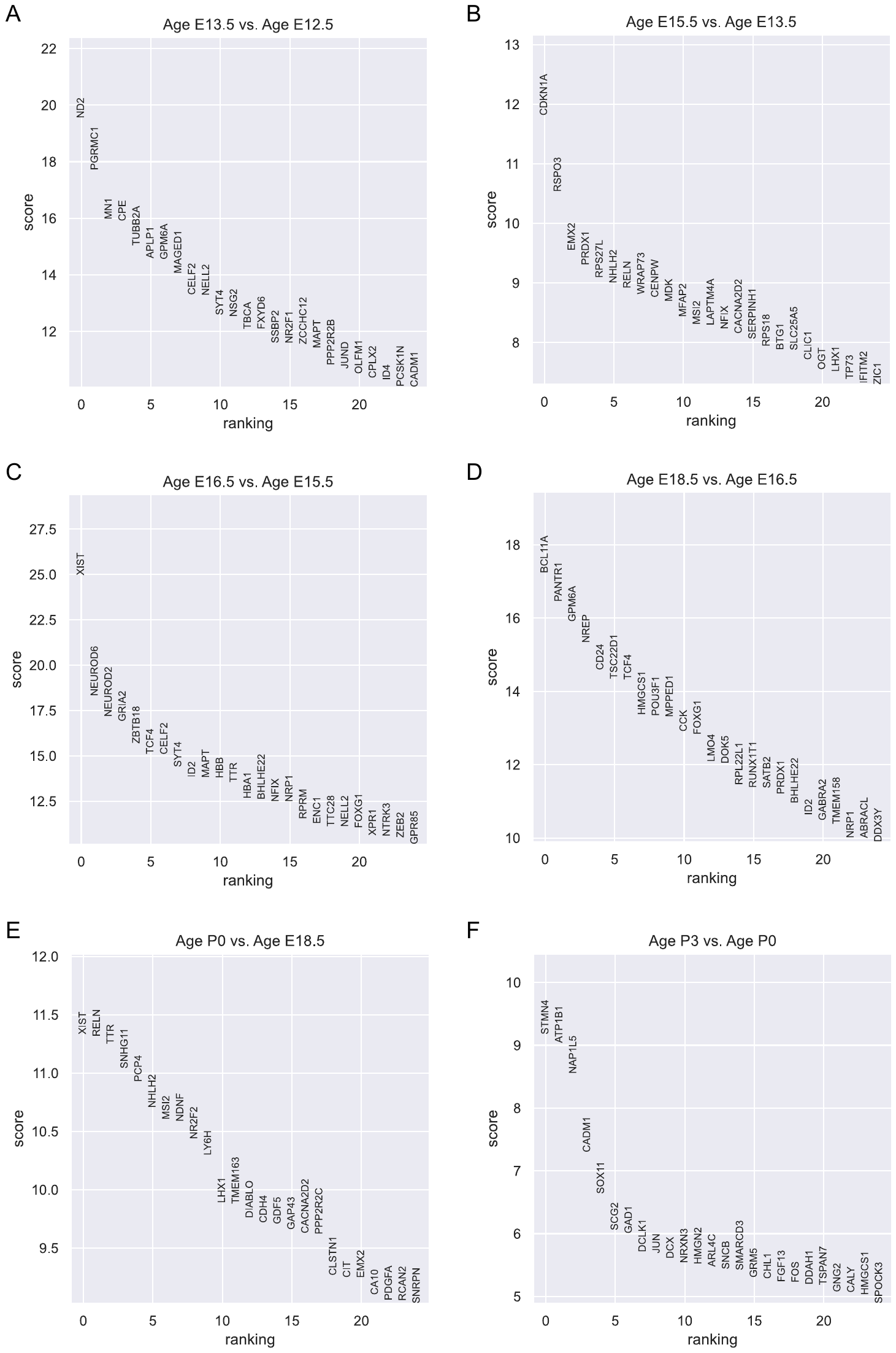

**Figure S40. Differentially expressed genes (DEGs) between adjacent age groups of Other neurons in Forebrain dataset.** Differential expression analysis using Wilcoxon rank-sum is conducted for each pair of adjacent age groups. Genes are ranked according to the z-score underlying the p-value computation and the top 25 rankings are displayed. (A) DEGs between E13.5 and E12.5. (B) DEGs between E15.5 and E13.5. (C) DEGs between E16.5 and E15.5. (D) DEGs between E18.5 and E16.5. (E) DEGs between P0 and E18.5. (F) DEGs between P3 and P0.

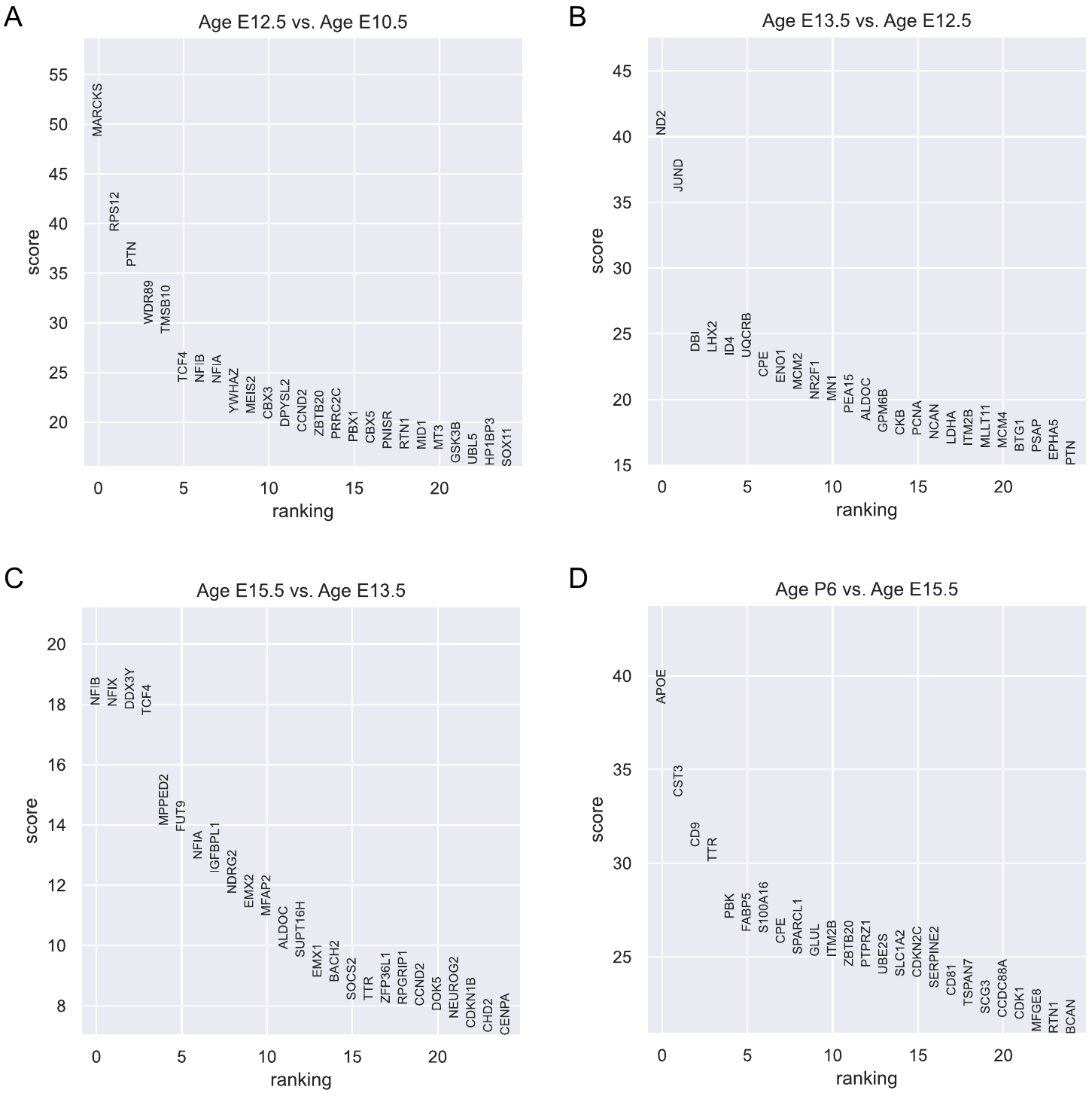

**Figure S41. Differentially expressed genes (DEGs) between adjacent age groups of RGC in Forebrain dataset.** Differential expression analysis using Wilcoxon rank-sum is conducted for each pair of adjacent age groups. Genes are ranked according to the z-score underlying the p-value computation and the top 25 rankings are displayed. (A) DEGs between E12.5 and E10.5. (B) DEGs between E13.5 and E12.5. (C) DEGs between E15.5 and E13.5. (D) DEGs between P6 and E15.5.

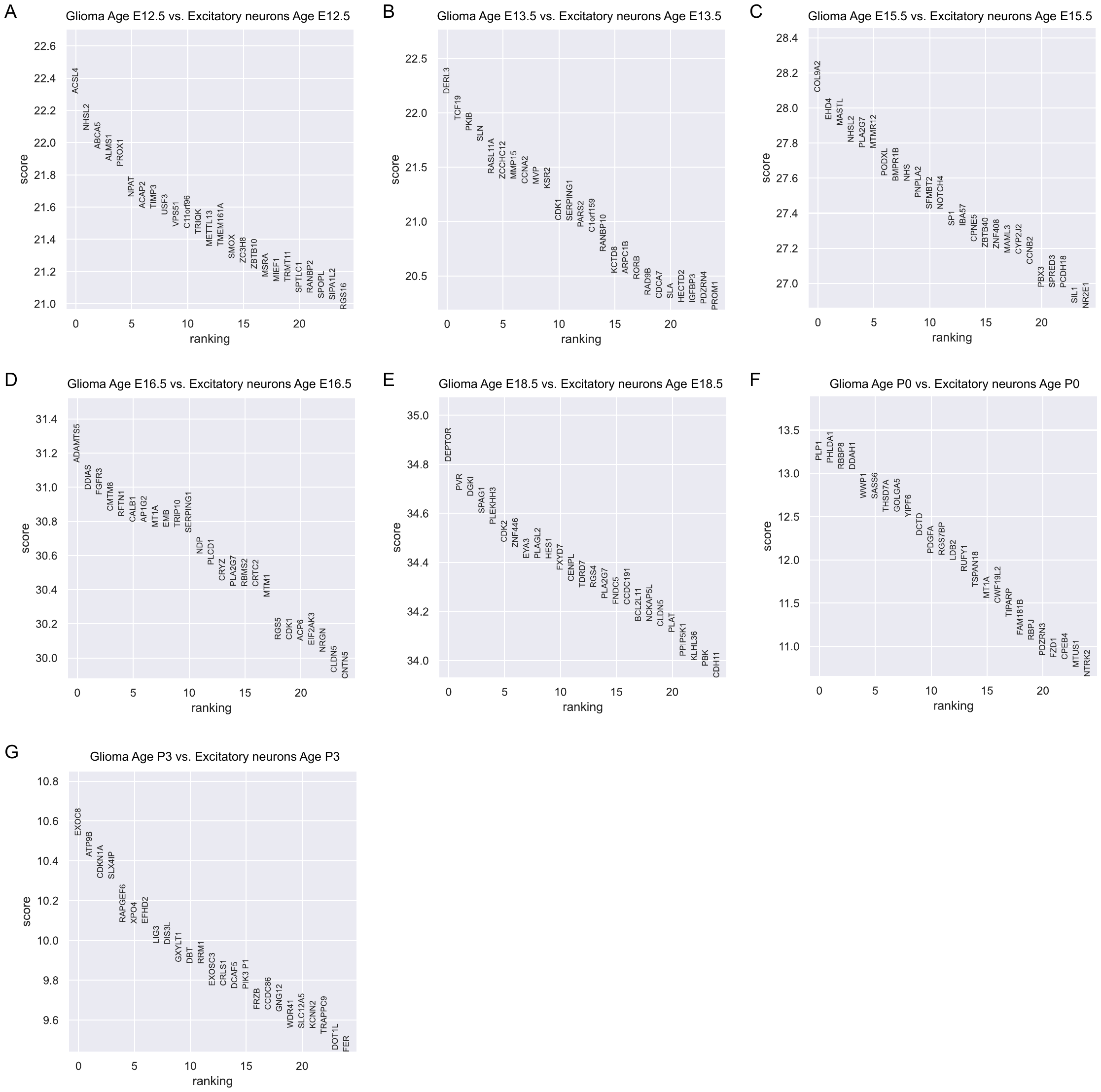

**Figure S42. Differentially expressed genes (DEGs) between glioma and Excitatory neurons in Forebrain dataset.** Differential expression analysis using Wilcoxon rank-sum is conducted for each age group. Genes are ranked according to the z-score underlying the p-value computation and the top 25 rankings are displayed. (A) DEGs for E12.5. (B) DEGs for E13.5. (C) DEGs for E15.5. (D) DEGs for E16.5. (E) DEGs for E18.5. (F) DEGs for P0. (G) DEGs for P3.

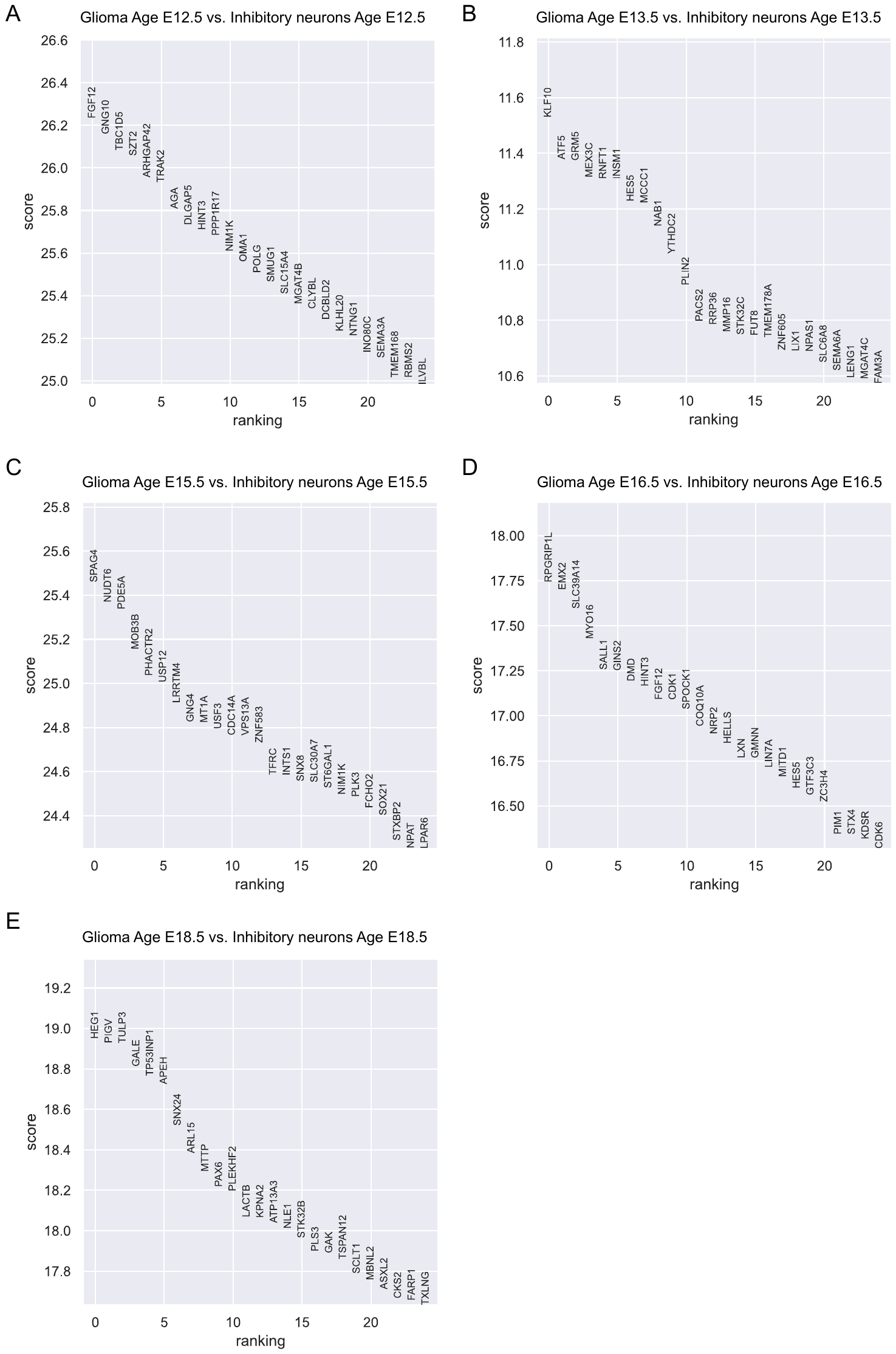

**Figure S43. Differentially expressed genes (DEGs) between glioma and Inhibitory neurons in Forebrain dataset.** Differential expression analysis using Wilcoxon rank-sum is conducted for each age group. Genes are ranked according to the z-score underlying the p-value computation and the top 25 rankings are displayed. (A) DEGs for E12.5. (B) DEGs for E13.5. (C) DEGs for E15.5. (D) DEGs for E16.5. (E) DEGs for E18.5.

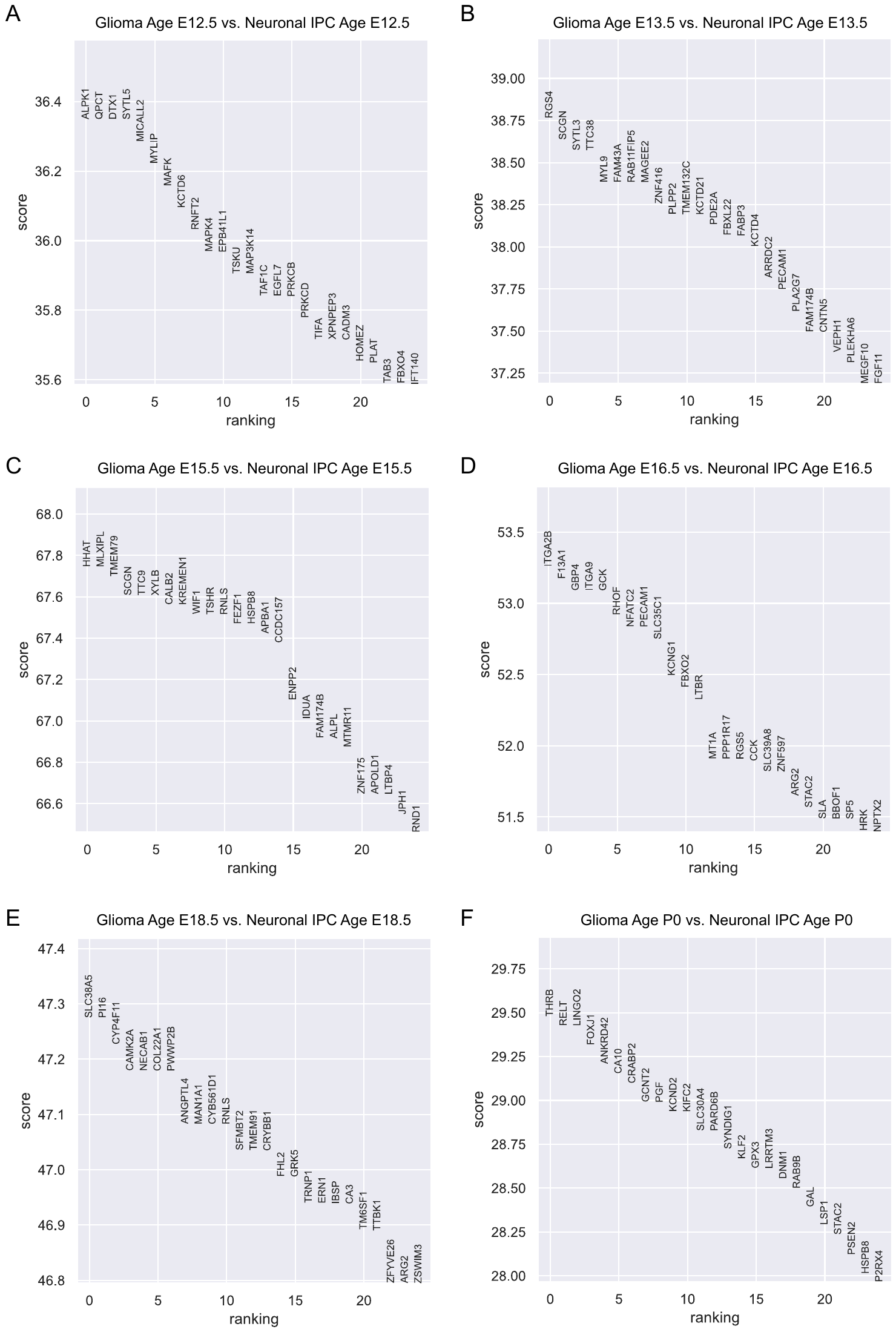

**Figure S44. Differentially expressed genes (DEGs) between glioma and Neuronal IPC in Forebrain dataset.** Differential expression analysis using Wilcoxon rank-sum is conducted for each age group. Genes are ranked according to the z-score underlying the p-value computation and the top 25 rankings are displayed. (A) DEGs for E12.5. (B) DEGs for E13.5. (C) DEGs for E15.5. (D) DEGs for E16.5. (E) DEGs for E18.5. (F) DEGs for P0.

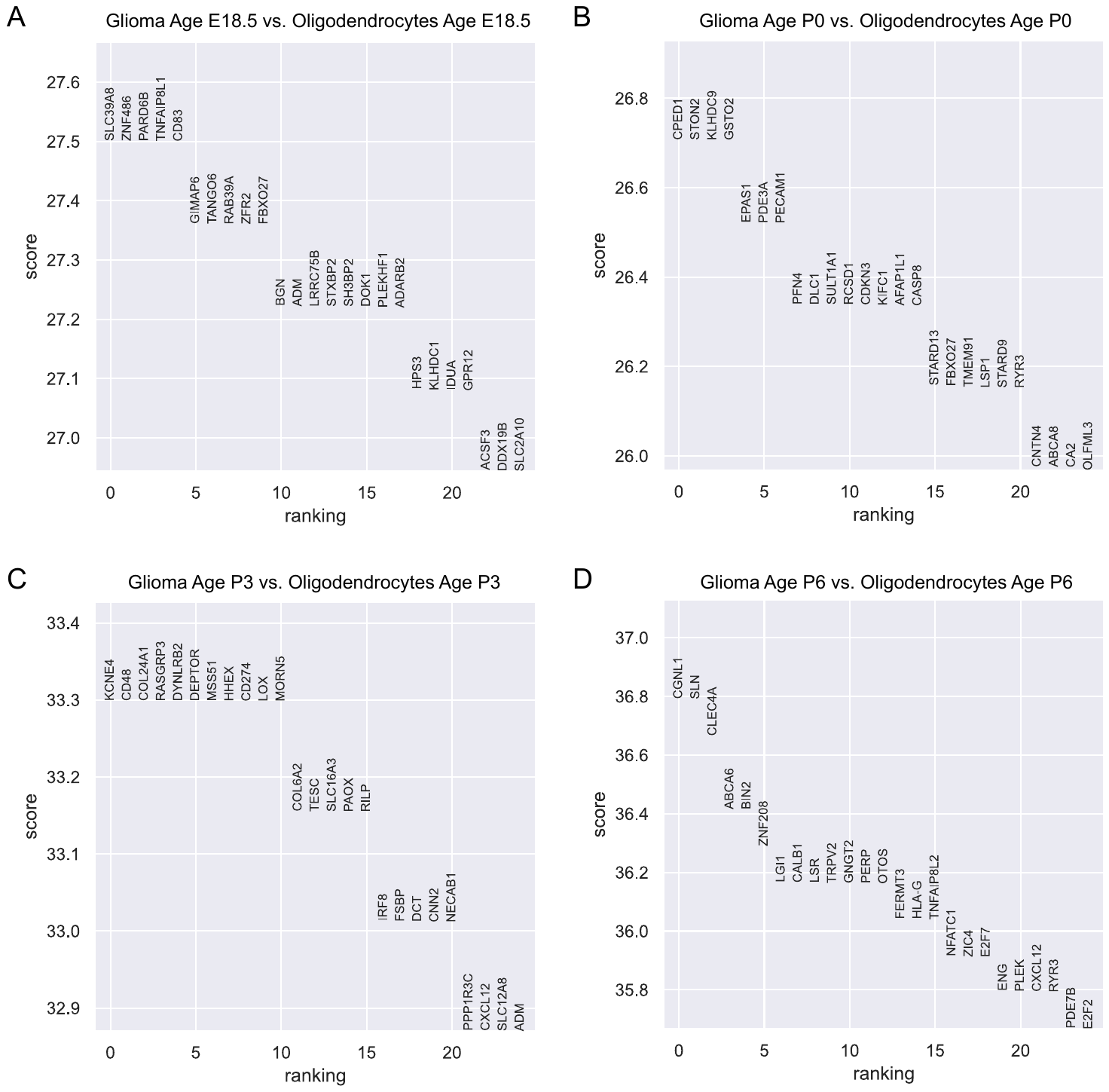

**Figure S45. Differentially expressed genes (DEGs) between glioma and Oligodendrocytes in Forebrain dataset.** Differential expression analysis using Wilcoxon rank-sum is conducted for each age group. Genes are ranked according to the z-score underlying the p-value computation and the top 25 rankings are displayed. (A) DEGs for E18.5. (B) DEGs for P0. (C) DEGs for P3. (D) DEGs for P6.

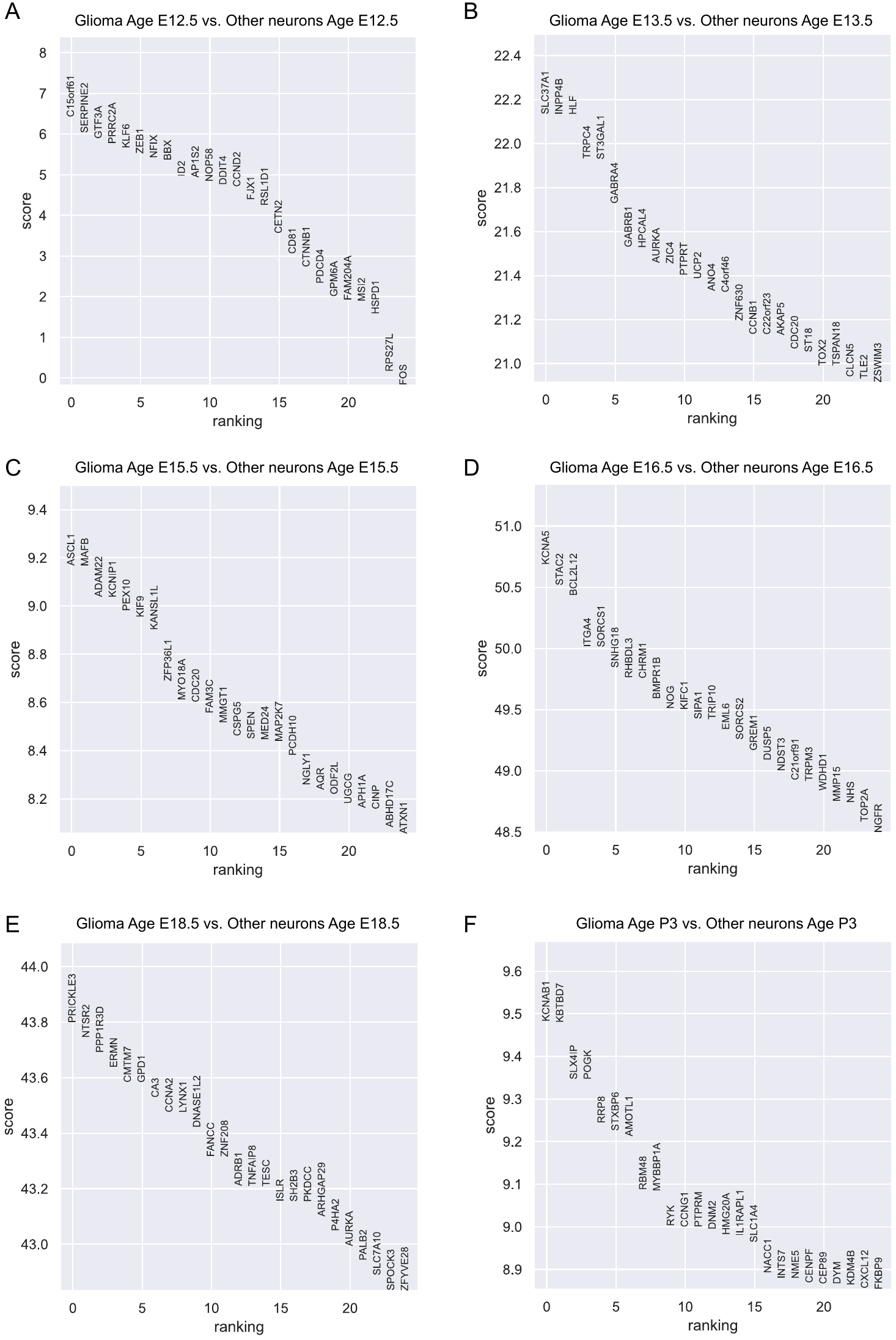

**Figure S46. Differentially expressed genes (DEGs) between glioma and Other neurons in Forebrain dataset.** Differential expression analysis using Wilcoxon rank-sum is conducted for each age group. Genes are ranked according to the z-score underlying the p-value computation and the top 25 rankings are displayed. (A) DEGs for E12.5. (B) DEGs for E13.5. (C) DEGs for E15.5. (D) DEGs for E16.5. (E) DEGs for E18.5. (F) DEGs for P3.

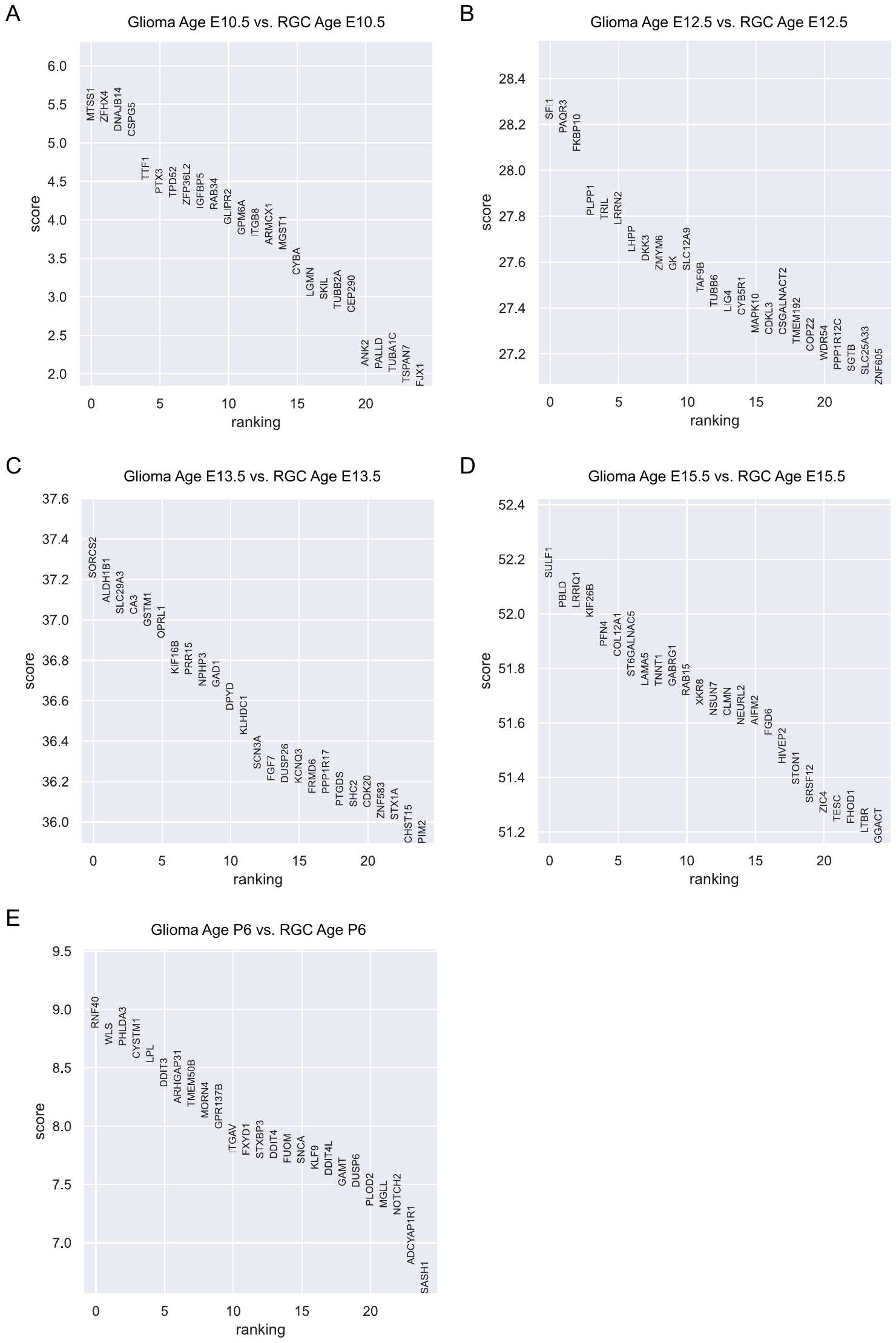

**Figure S47. Differentially expressed genes (DEGs) between glioma and RGC in Forebrain dataset.** Differential expression analysis using Wilcoxon rank-sum is conducted for each age group. Genes are ranked according to the z-score underlying the p-value computation and the top 25 rankings are displayed. (A) DEGs for E10.5. (B) DEGs for E12.5. (C) DEGs for E13.5. (D) DEGs for E15.5. (E) DEGs for P6.

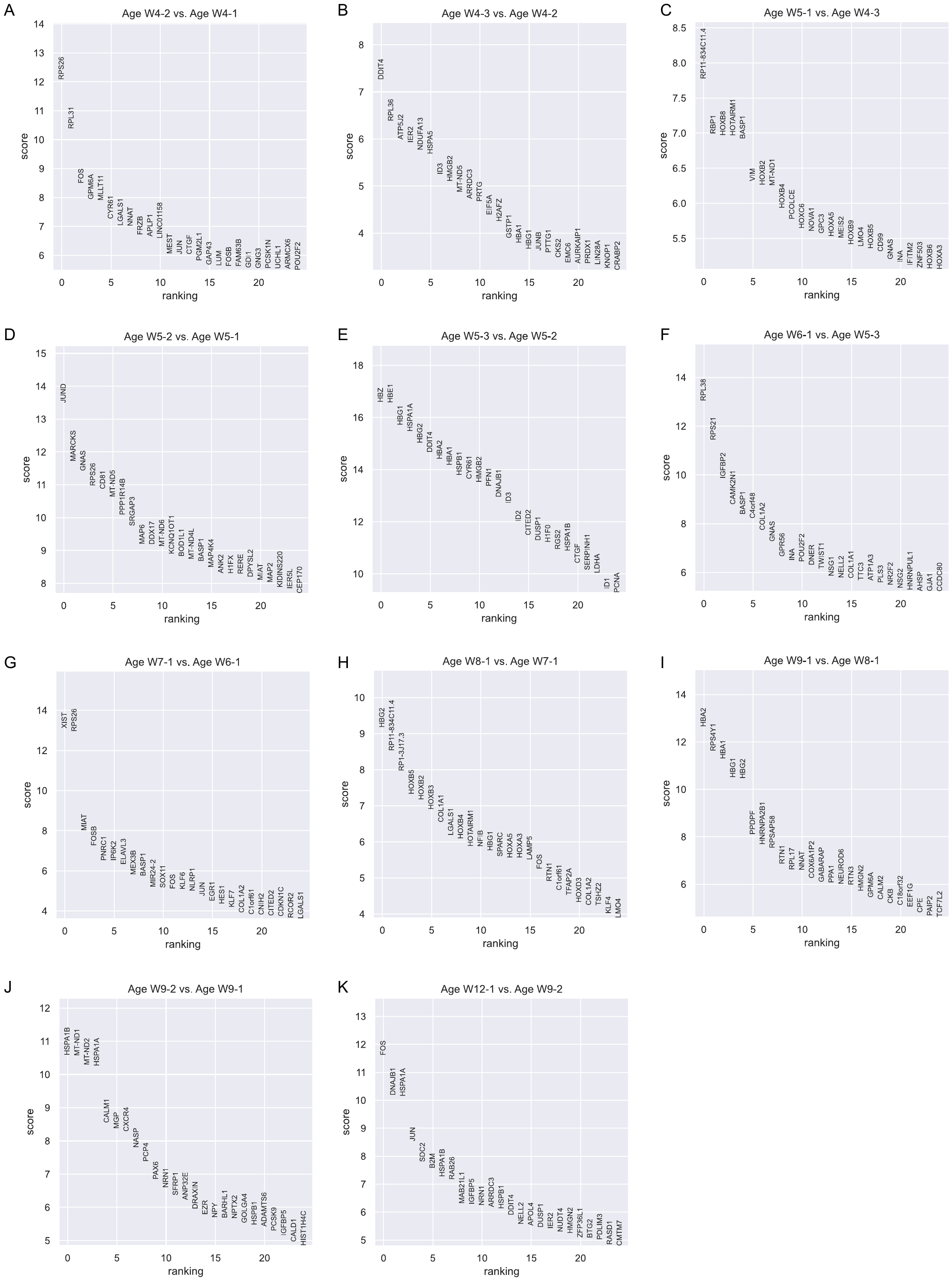

**Figure S48. Differentially expressed genes (DEGs) between adjacent age groups of GABA N 9 in GSE155121 dataset.** Differential expression analysis using Wilcoxon rank-sum is conducted for each pair of adjacent age groups. Genes are ranked according to the z-score underlying the p-value computation and the top 25 rankings are displayed. (A) DEGs between W4-2 and W4-1. (B) DEGs between W4-3 and W4-2. (C) DEGs between W5-1 and W4-3. (D) DEGs between W5-2 and W5-1. (E) DEGs between W5-3 and W5-2. (F) DEGs between W6-1 and W5-3. (G) DEGs between W7-1 and W6-1. (H) DEGs between W8-1 and W7-1. (I) DEGs between W9-1 and W8-1. (J) DEGs between W9-2 and W9-1. (K) DEGs between W12-1 and W9-2.

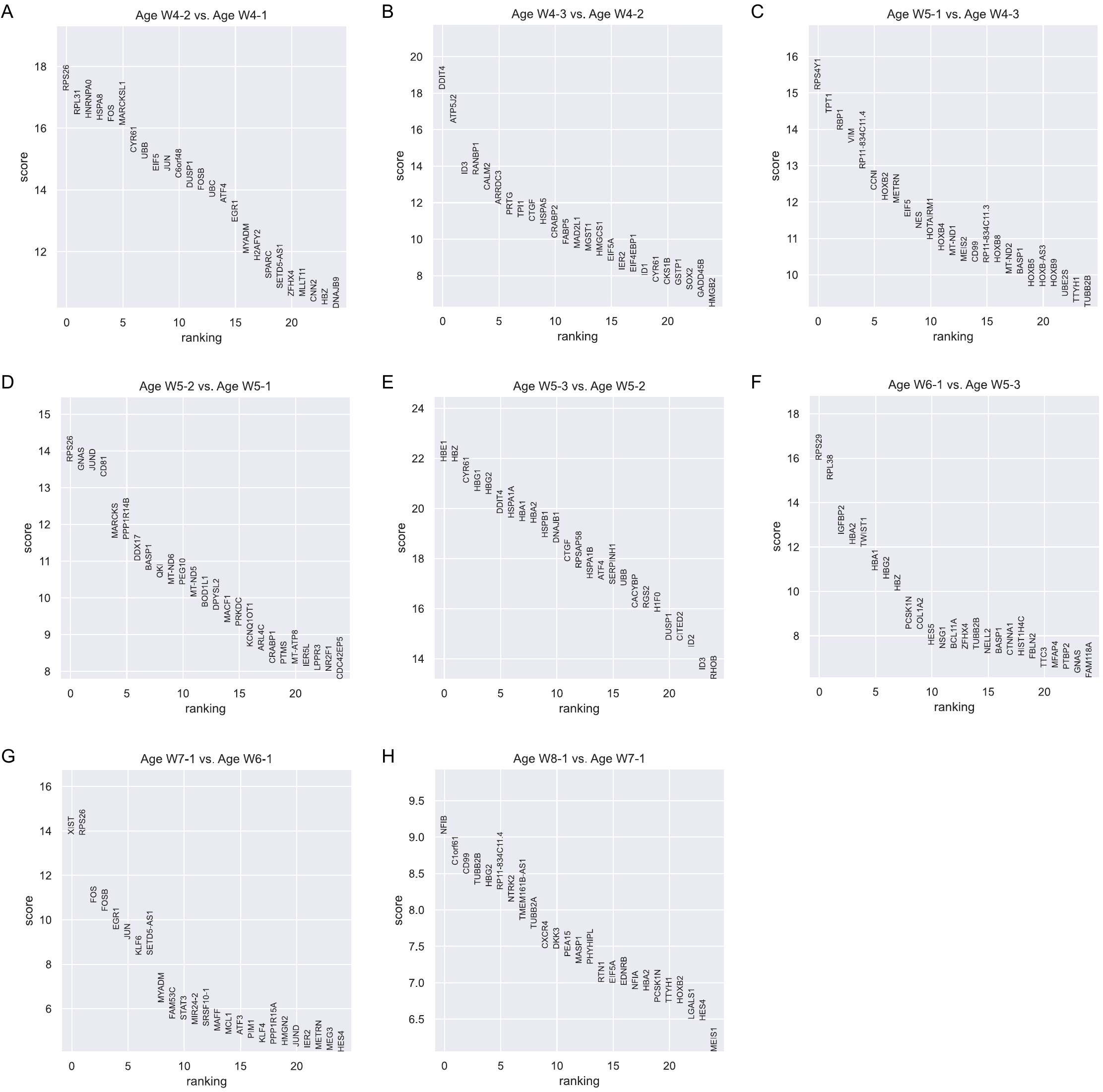

**Figure S49. Differentially expressed genes (DEGs) between adjacent age groups of NSC 12 in GSE155121 dataset.** Differential expression analysis using Wilcoxon rank-sum is conducted for each pair of adjacent age groups. Genes are ranked according to the z-score underlying the p-value computation and the top 25 rankings are displayed. (A) DEGs between W4-2 and W4-1. (B) DEGs between W4-3 and W4-2. (C) DEGs between W5-1 and W4-3. (D) DEGs between W5-2 and W5-1. (E) DEGs between W5-3 and W5-2. (F) DEGs between W6-1 and W5-3. (G) DEGs between W7-1 and W6-1. (H) DEGs between W8-1 and W7-1.

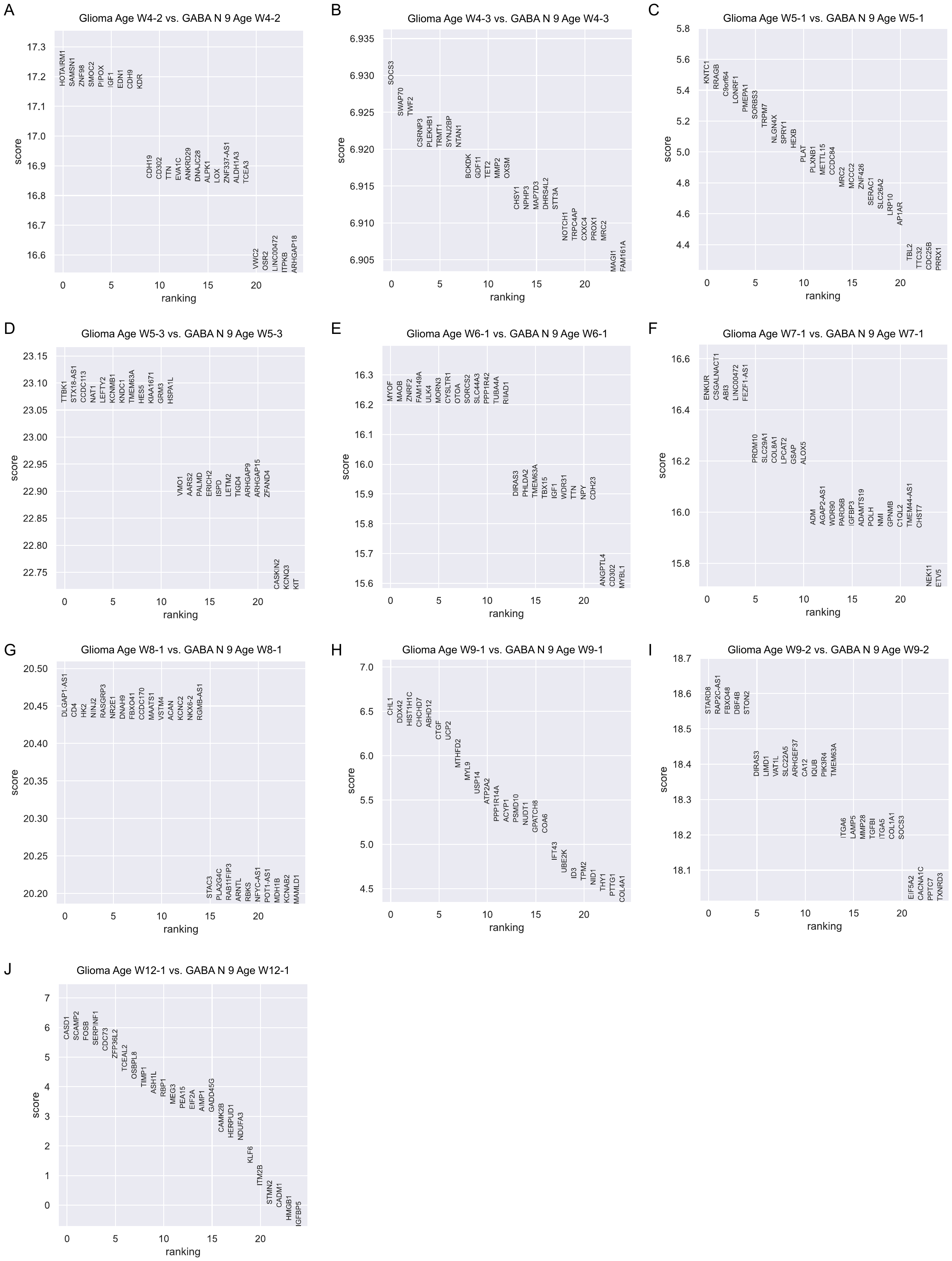

**Figure S50. Differentially expressed genes (DEGs) between glioma and GABA N 9 in GSE155121 dataset.** Differential expression analysis using Wilcoxon rank-sum is conducted for each age group. Genes are ranked according to the z-score underlying the p-value computation and the top 25 rankings are displayed. (A) DEGs for W4-2. (B) DEGs for W4-3. (C) DEGs for W5-1. (D) DEGs for W5-3. (E) DEGs for W6-1. (F) DEGs for W7-1. (G) DEGs for W8-1. (H) DEGs for W9-1. (I) DEGs for W9-2. (J) DEGs for W12-1.

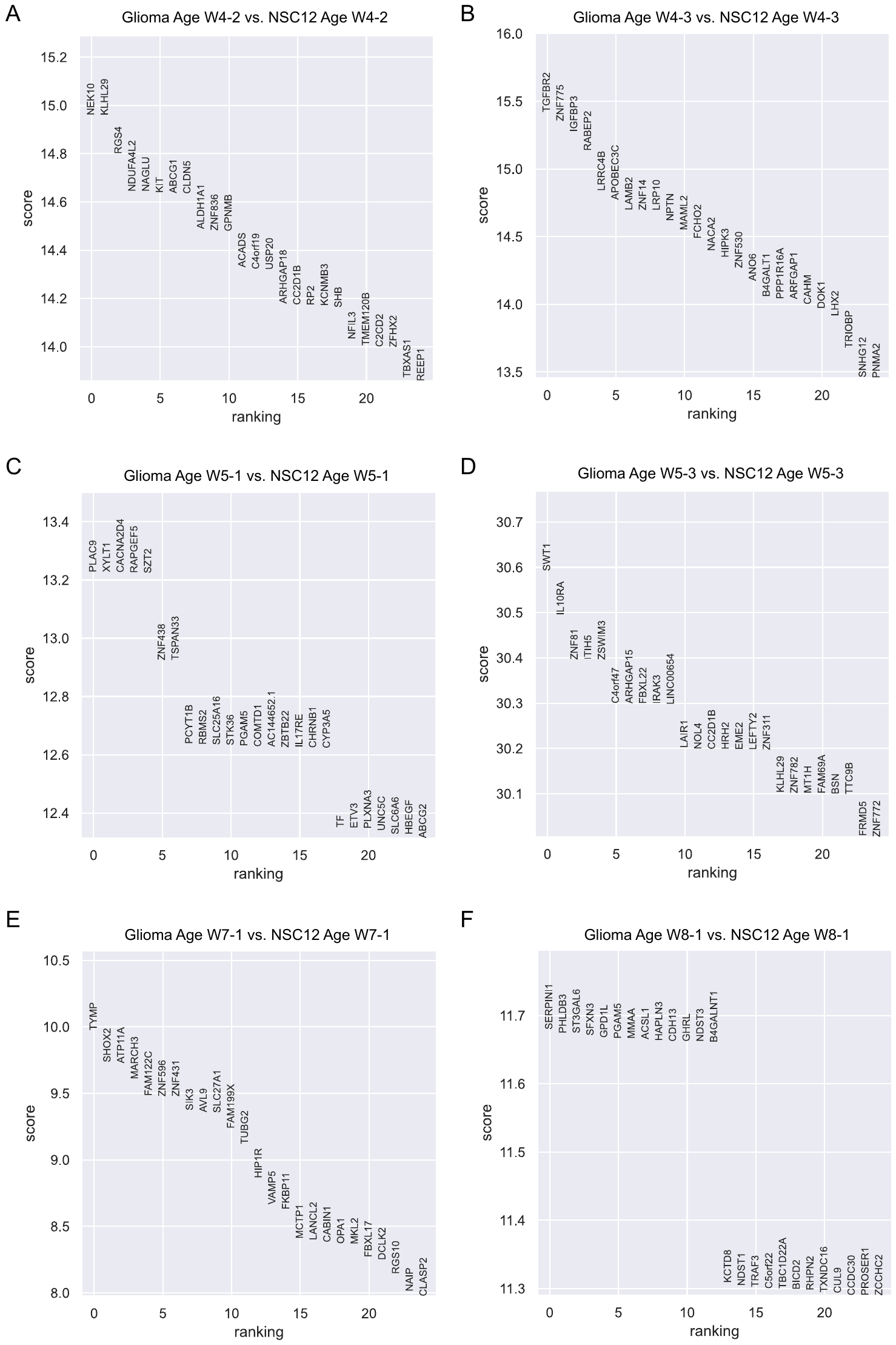

**Figure S51.** Differentially expressed genes (DEGs) between glioma and NSC 12 in GSE155121 dataset. Differential expression analysis using Wilcoxon rank-sum is conducted for each age group. Genes are ranked according to the z-score underlying the p-value computation and the top 25 rankings are displayed. (A) DEGs for W4-2. (B) DEGs for W4-3. (C) DEGs for W5-1. (D) DEGs for W5-3. (E) DEGs for W7-1. (F) DEGs for W8-1.
